# Revisiting the Briggs ancient DNA damage model: a fast regression method to estimate postmortem damage

**DOI:** 10.1101/2023.11.06.565746

**Authors:** Lei Zhao, Rasmus Amund Heriksen, Abigail Daisy Ramsøe, Rasmus Nielsen, Thorfinn Sand Korneliussen

## Abstract

**Motivation:** One essential initial step in the analysis of ancient DNA is to authenticate its ancientness to ensure reliable conclusions. That is, meticulously assessing whether next-generation sequencing reads exhibit ancient characteristics, with a particular focus on the postmortem damage (PMD) signal induced by cytosine deamination in the fragments termini. We present a novel statistical method implementation in a fast multithreaded program ngsBriggs that enables the rapid quantification of PMD by calculation of the Briggs ancient damage model parameters (Briggs parameters).

**Results:** Using a fast multinomial regression approach, ngsBriggs accurately models the Briggs parameters, quantifying the PMD signal from single and double-stranded DNA regions. We revisit and extend the original Briggs model, with ngsBriggs modeling PMD signals for contemporary sequencing platforms. Furthermore, ngsBriggs asserts itself as a reliable and consistent tool, by accurately estimating the Briggs parameters across a variety of contamination levels. The classification accuracy of ngsBriggs significantly exceeds the current tool available when discerning ancient-from modern sequencing reads to decontaminate samples. Our novel method and implementation ngsBriggs outperforms existing tools regarding computational speed and accuracy, establishing its practicality and usability. Our tool, ngsBriggs offers a practical and accurate toolset for researchers seeking to authenticate ancient DNA and improve the quality of their data.

**Availability:** https://github.com/lz398/metadamage_briggs

## 1 Introduction

Ancient DNA (aDNA) refers to the preserved genetic material of ancient organisms. Analyzing aDNA has proven to be an essential tool for researchers to study the past. It has, for example, aided a deeper understanding of ancestral population history [1, 2] and the dynamics of ancient ecosystems [7, 12].

During the last few decades, the field of aDNA has seen an increase in both the quality and quantity of data, with the number of published ancient genomes surpassing 10,000 at the end of 2022 [15].

The primary structure of DNA consists of a linear sequence of nucleotides (A, T, C or G) connected together by a phosphate backbone. DNA is double-stranded - the two strands are connected by hydrogen bonds between the nucleotides. Due to the passage of time and prolonged exposure to various environmental conditions, DNA undergoes several conformational alterations, known as postmortem damage (PMD). PMD in the double-stranded DNA molecule manifests mainly as nicks and deamination. A nick is a break in the backbone of either strand in the fragment. This can cause structural instability, which mediates a complete break where two smaller sub-fragments are formed. The process does not happen at the same place in the backbone of both strands, thus there might be single-stranded DNA in the termini of the fragment. The single-stranded region of the ancient molecule is termed the overhang, which can be defined as either the 5’ or 3’ overhang. Another hallmark of aDNA PMD is deamination of cytosine, which converts cytosine to uracil, which happens with an increased frequency in the single-stranded part of the fragment compared to the double-stranded region, when sequencing a deaminated cytosine, uracil will be sequenced as thymine creating an apparent C*→*T substitution. Lastly, DNA from the (post-) depositional environment can contaminate the endogenous DNA, such that the resulting surviving DNA fragments might have multiple sources. [4, 23]). These characteristics make the truly ancient DNA distinguishable from its modern counterpart. This disparity facilitates the further development of methods to authenticate the ancient content, allowing researchers to ensure that their sequencing data is genuinely ancient.

The process of obtaining DNA sequences from biological material follows a laboratory protocol that can be divided into 1) DNA extraction and purification, 2) library preparation, and 3) DNA sequencing. The first step is sample-specific, whereas the second is sequencing platform-specific. The most common sequencing approach is the sequencing-by-synthesis, which requires that each original DNA fragment is ligated with known adapter sequences. The double-stranded library preparation includes several steps [3, 17]:

1. **Blunt-end Repair:** Removal of the 3’ overhang of the molecule through 3’*→* 5’ exonuclease activity of *T4 DNA Polymerase* while also catalyzing the 5’ overhangs fill-in synthesizing the DNA in 5’*→* 3’ direction (See part I of **Figure** 1) [20, **22]**. With the subsequent 5’ phosphorylation of *T4 Polynucleotide Kinase* required for adapter ligation.

**Figure 1.**
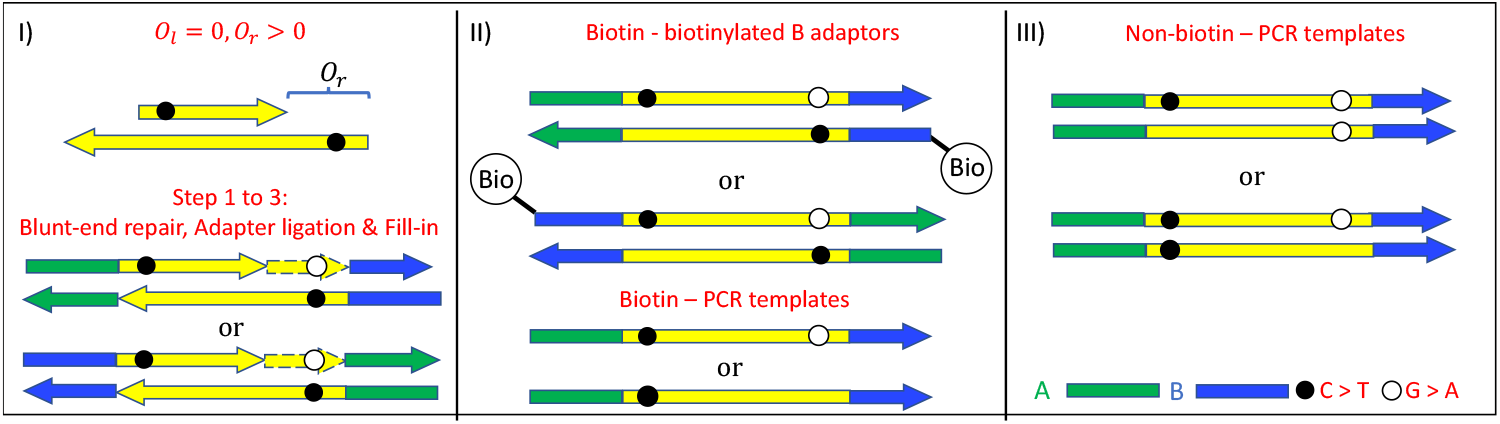
Illustrative representation of the laboratory protocols and resulting deamination pattern, for this scenario, the 3’ overhang is removed during the blunt-end repairs (*O*_*l*_ = 0, left overhang), while the complementary substitution is observed in the 5’ overhang (*O*_*r*_ > 0, right overhang). *I)* Initial steps in the double-stranded library preparation, shared across the two distinct DNA sequencing platforms. *II*) The biotin model, with the fragments with the biotinylated B adapter captured by streptavidin beads with solely the ancient strands with A-to-B orientation serving as PCR templates. *III*) In the non-biotin model, the ancient strands with A-to-B orientation and the ancient strands with B-to-A orientation are equally likely to serve as PCR templates (for simplicity the output PCR templates are depicted with 5’ A-to-B 3’ direction). This single scenario (*O*_*l*_ = 0,*O*_*r*_ > 0) illustration is depicted with a greater amount of detail in Quadrant II of SI **Figure** 2.
2. **Adapter Ligation:** Random ligation of the adapters (A or B) with *T4 DNA Ligase* catalyzing the phosphodiester bond with the blunt end. Those fragments ligated solely with the A or B adapter are non-functional, with only fragments ligated with both A and B adapters contributing to later processes.
3. **Adapter Fill-in:** Followed by *Bst DNA Polymerase, Large Fragment* with strand displacement activity extending the nick present on one strand between the adapter and template (adapter fill-in) (not visualized in part I of **Figure** 1). This strand displacement activity will in the presence of single-stranded nicks in double-stranded inserts perform a similar downstream displacement during the DNA synthesis (nick-fill, as visualized with nicks in both strands SI **Figure** 3; nicks in a single strand SI **Figure** 4)
4. **PCR and denaturing:** The resulting double-stranded DNA is denatured, and both strands may act as the templates of PCR in accordance with different PCR protocols (e.g. emulsion PCR and Illumina sequencing) described below.

In the Briggs’ 2007 article [3], the authors mathematically model the effect of PMD, deducing four parameters (*λ, δ*_*d*_, *δ*_*s*_, *ν*) (denoted throughout as Briggs parameters):

*λ*: the parameter related to the length distribution of the 5’ overhang.

*δ*_*d*_: the deamination level in the double-stranded region.

*δ*_*s*_: the deamination level in the single-stranded region.

*ν*: the nick frequency.

The Briggs article models the effect of the historical Roche 454 sequencing platform solely using one strand for the PCR, by fixating the other using a biotinylated adapter, which will, in the context of this article, be called the biotin model (see part II of **Figure** 1).

Unlike the emulsion PCR approach adopted by the 454 platform, libraries built for Illumina sequencing do not use biotinylated adapters.

Due to the differences in PCR templates between the two methods, using a biotin model to describe deamination patterns of reads generated on modern Illumina platforms is not suitable. In this paper, we extend the idea of the biotin model (part II of **Figure** 1), and develop a non-biotin model for Illumina-sequenced deamination (part III of **Figure** 1) [17].

Some methods have been developed for determining if sequencing reads are produced from ancient DNA. Some use overall nucleotide differences to a reference (e.g. mapDamage [9]) at each position in a read but do not rely on an explicit model of damages. In contrast, mapDamage 2.0 [11] directly infers three of the four Briggs parameters by using a Bayesian Markov chain Monte Carlo approach (MCMC). Other methods, e.g. PMDTools [21] compute a test statistic at the sequence read level for discriminating between reads that exhibit postmortem damage characteristics and those that do not.

To allow for the efficient estimation and calculation of the Briggs parameters, we present a novel statistical method called ngsBriggs that utilizes multinomial regression.

Our method estimates the four parameters of the Briggs model based on a set of sequencing reads. As such, ngsBriggs is the most current and relevant tool for estimating deamination patterns of modern Illumina ancient DNA libraries. Furthermore, within the same framework (and by using an external estimate of the contamination fraction), we can also compute, for each read, a probability of it originating from endogenous ancient DNA. This ability to differentiate between the truly ancient reads and the modern contamination, combines the functionalities of previous bioinformatical tools in one coherent framework.

## 2 Materials and Methods

Both the biotin and non-biotin models attribute any observed PMD signals to nick’s placement, the degree of deamination within both the single- and double-stranded regions, and the length of the 5’ overhangs. The inferred *λ* determines the distribution of the 5’ overhang lengths (both the left and right 5’ overhangs, as defined in the original Briggs article [3], share the same distribution):

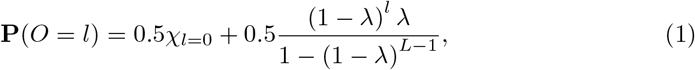

where *χ* is an indicator function, *L* is the focal fragment’s length and *l* is the overhang length. Notice that we make no distinction between the left (denoted as *O*_*l*_) and right (denoted as *O*_*r*_) overhang length distribution and assume that the biochemical process that generates the 5’ and 3’ overhangs are similar competing processes. However, we would only observe the 5’ overhang due to the limitations of the double-stranded library preparation, at which point blunt-end repair removes any 3’ overhang and only retains 5’ overhangs. Furthermore, we require the combined total length of the left 5’ overhang and the right 5’ overhang on the same aDNA fragment that cannot exceed *L* − 2 (which was also assumed in the original Briggs paper [3]). In practice, reads with long overhangs are not observed due to the configuration being too unstable and thus the fragment would not be chemically feasible.

In theory, nicks can occur at any position of an aDNA fragment. However, the nicks in the single-stranded region will not be observed, as these simply decrease the overhang length. As such our model will solely consider the nicks within the double-stranded region. Nicks are assumed to occur uniformly along the genome, with a rate (*ν*) per site. In the original fragment (not the observed sequencing products) we might have multiple nicks; however, after library preparation, all nucleotides downstream from the first nick will be removed and replaced according to the opposite strand (See SI **Figure** 3 and 4 for cases when both strands of the fragment have nicks and only one strand of the fragment has nicks; details of nick placement are discussed in SI **Subsection** 2.3). In the model, we therefore focus on the first nick in the double-stranded region on each strand, and we only show the first nick in the relevant figures and will refer to the first nick as “the nick”.

### 2.1 Output PCR Templates

After the ligation of adapters to the original aDNA fragments, the biotin model, which follows the protocol with biotinylated adapters, will only utilize the original aDNA strands with the adapter direction 5’ A-to-B 3’ as the downstream PCR templates (as shown in part II of **Figure** 1). In the non-biotin case, the original strand with the A-to-B direction will be used directly as the PCR template, while the B-to-A one will be reverse-complemented before serving as the downstream PCR template. The original ancient strands and their reverse complement have an equal chance to serve as the downstream PCR templates (as shown in part III of **Figure** 1).

### 2.2 Inference of parameters of the Briggs model

When ngsBriggs employs multinomial regression to infer the Briggs parameters, it will create and utilize a mismatch matrix across all reads. The mismatch matrix is the position-specific nucleotide substitution count relative to a reference genome (one example is shown in **Table** 2 of the SI). Clearly, the count of deamination-specific substitutions C→T and G→A in the mismatch matrix will be inflated by sequencing errors and true biological variation. However, ngsBriggs corrects for this by quantifying these errors via counting the nucleotides that differ from the expected C→T and corresponding G→A at each cyclic position, assuming all nucleotide substitutions unrelated to PMD at the same cyclic position have an equal likelihood of occurring across all sequence reads.

Using these substitution counts, the effect of the sequencing errors and true biological variations in the mismatch matrix can be removed before inferring the parameters (SI **Section** 3.1.4).

We can classify the probabilities of four spatial relationships of the focal position (*n*), potential nick, and the left or right 5’ overhang:

1. Focal position *n* is within the double-strand region and downstream (position *n*, downstream and upstream are in the sense of 5’ end to 3’ end) of the possible first nick on this strand. Denoted as *p*_1_(*n*; *L*).
2. Focal position *n* is within the right 5’ overhang region. Denoted as *p*_2_(*n*; *L*).
3. Focal position *n* is within the double-strand region and upstream of the possible first nick (this also includes the case without any nick). Denoted as *p*_3_(*n*; *L*).
4. Focal position *n* is within the left 5’ overhang region. Denoted as *p*_4_(*n*; *L*).

The four spatial relationship probabilities are functions of not only *n* and the focal strand’s length *L*, but also the four damage parameters, *λ, δ*_*d*_, *δ*_*s*_, and *ν* as seen in SI **Section** 3.1, with the probabilities satisfying *p*_1_(*n*; *L*) + *p*_2_(*n*; *L*) + *p*_3_(*n*; *L*) + *p*_4_(*n*; *L*) = 1.

If no sequencing errors or contaminating DNA strands are considered, a deaminated C*→*T given the reference nucleotide is C at position *n* (counted from 5’ end) on a randomly chosen original ancient strand (a potential template for both models) can only be observed when the spatial relationship satisfies either type 3 (*p*_3_(*n*; *L*), with *n* being in the double-stranded region with the deamination level *δ*_*d*_) or type 4 (*p*_4_(*n*; *L*), with *n* in the single-stranded region, and deamination level *δ*_*s*_). Hence, the chance of observing a C*→*T given a reference nucleotide of C is *p*_3_(*n*; *L*)*δ*_*d*_ + *p*_4_(*n*; *L*)*δ*_*s*_. Similarly, the chance of observing a G*→* A given a reference nucleotide, G, is *p*_1_(*n*; *L*)*δ*_*d*_ + *p*_2_(*n*; *L*)*δ*_*s*_.

However, when the focal position *n* is on a randomly chosen reverse complement of the original strand (a potential PCR template only for the non-biotin model), a C*→*T (or a G*→*A) is equivalent to a G*→*A (or a C*→*T) at position *L − n* + 1 on the original strand. Therefore the probability of observing either C*→*T or a G*→*A, at position *n* on the complementary strand, is given as *p*_1_(*L − n* + 1; *L*)*δ*_*d*_ + *p*_2_(*L − n* + 1; *L*)*δ*_*s*_ and *p*_3_(*L−* n + 1; *L*)*δ*_*d*_ + *p*_4_(*L− n* + 1; *L*)*δ*_*s*_, respectively.

Given these relationships, we can define the theoretical deamination frequencies for the model-specific PCR templates (see **Figure** 1) with the following formulae.

**Equations** 2 and 3 represent the biotin model (abbreviated as b), with **Equations** 4 and 5 representing the non-biotin model (abbreviated as nb).

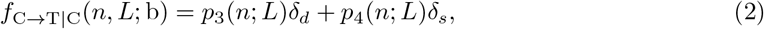

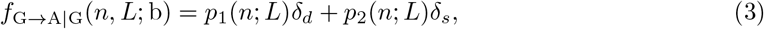

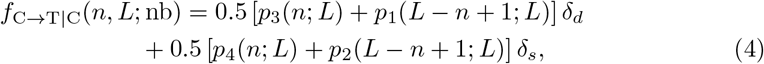

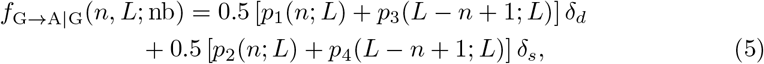

where *f*_X*→*Y|X_(*n, L*; *m*) denotes the frequency of nucleotide change X*→*Y given X at position *n* in a fixed-fragment-length (*L*) sample of a specified model (*m*). Throughout the rest of this work, we will use the notation *f*_X*→*Y|X_(*n, L*) if the relevant formulae are applied to both models.

The above theoretical frequencies can be further incorporated with sequencing errors and potential contamination (as shown in Sections 3.1.1-3.1.4 of the SI). For consistency and simplicity, we will still use the same notations to represent the corresponding frequencies with errors and contamination. Using the mismatch matrix, ngsBriggs conducts the regression to maximize the following log-likelihood,

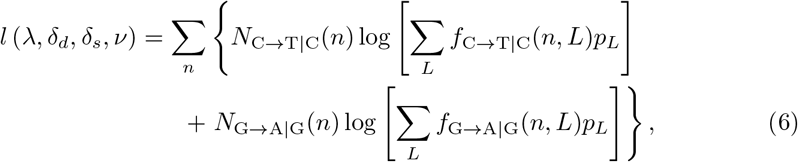

where *N*_X*→*Y|X_(*n*) is the actual counts of nucleotide X*→*Y given X at position *n* of a randomly chosen fragment from the mismatch matrix, and *p*_*L*_ is the proportion of fragments of length *L* in the sample after the PCR.

### 2.3 Ancient read probability in the presence of contamination

Once the four Briggs model parameters have been inferred, we can also compute a posterior probability of each read being ancient for samples contaminated with modern human DNA. It should be noted that the existing methods, e.g., PMDtools, can only distinguish ancient reads from their modern counterparts based on the observed PMD signals, hence it will be difficult for them to identify those reads originating from ancient material but without PMD patterns. In contrast, ngsBriggs combines information on length distribution and PMD signals. By assuming the differences in lengths between modern and ancient DNA reads, it gains more power to identify ancient reads, even without evidence of PMD.

We expect the length distribution of ancient reads to be distinguishable from that of modern reads (ancient reads are generally shorter) and have an elevated C*→* T frequency at the ends; this leads us to the following two assumptions:

1, By assuming that the lengths of the endogenous and modern contamination follow distinguishable constrained normal distributions *𝒩*_*b*_ (*μ*_*a*_, *σ*_*a*_) and *𝒩*_*b*_ (*μ*_*m*_, *σ*_*m*_) (The assumption of normality is strong but for efficient computation, it can be viewed to define a score to distinguish reads. In **Figure** 3 and SI **Figures** 24, we have proved it works for different cases, e.g., log-normal length distributions, which violate this assumption.), we can estimate the values of (*μ*_*a*_, *σ*_*a*_, *μ*_*m*_, *σ*_*m*_) if the overall modern contamination amount *r* is provided.

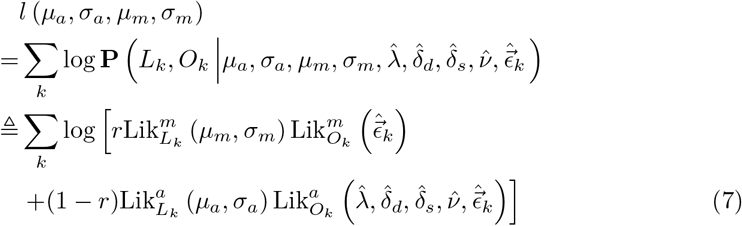

where **P** (*L*_*k*_, *O*_*k*_| *· · ·*) represents the joint probability of ancient strand length *L*_*k*_ and the nucleotide misincorporation pattern *O*_*k*_. Here 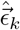 is the sequencing error per position of strand *k* provided by the Phred-scaled base quality score, and 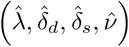 are the inferred damage parameters based on the mismatch matrix. The notation Lik : represents the likelihood function associated with the subscripted data (i.e., either *L*_*k*_ or(*O*_*k*_), and the superscript specifies whether the fragment is assumed to be ancient (*a*) or modern (*m*). The derivation of Lik: is in Sections 4.1 and 4.2 of the SI.

2, Based on the estimates 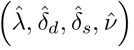 and 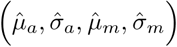, we can now calculate the posterior probability of being ancient for each strand given the observed nucleotide misincorporation pattern and strand length, e.g., for strand *k*, the posterior probability can be written as follows,

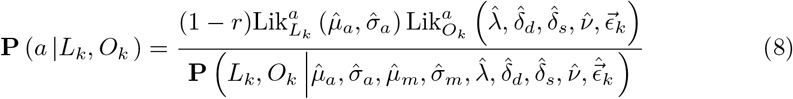

The detailed derivation of this expression is given in **Section** 4 of the SI.

### 2.4 Published data and simulated files

#### Simulated files for inference of Briggs parameters

To determine the accuracy of our regression framework, we used ngsBriggs inference on simulated files. Using the simulation software NGSNGS [10] we generated 100 files for the biotin- and 100 for the non-biotin model, equally separated into five groups with a varying number of reads, i.e. 10^3^, 10^4^, 10^5^, 10^6^ and 10^7^ to test different scenarios. All of these files were simulated with a set of “default” deamination parameters as estimated in the original Briggs article [3], i.e., 0.36, 0.0097, 0.68, 0.024 (*λ, δ*_*d*_, *δ*_*s*_, *ν*, SI **Section** 5 for more details).

#### Published data for different populations for inference of Briggs parameters

We also applied our program on previously published ancient samples across different time periods, i.e., 121 individuals from Damgaard (2018) [6] (termed DA), 219 individuals from Allentoft (2022) [1] (NEO), 67 individuals from Allentoft (2015) [2] (RISE), and finally 379 individuals from Margaryan 2020 [16] (VK). As such, we could test our inference models on samples with potentially different deamination patterns due to age, environmental conditions, and library preparation.

#### Simulated files for contamination scenarios

To investigate the effect of contamination, we simulated files with a mixture of ancient- and modern sequencing reads, using contamination proportions of 10%, 20%, 30%, 40%, or 50%. For each of the contamination levels, we simulated numerous PMD signals (either biotin or non-biotin) by varying the overhang length *λ* and single-stranded deamination rate *δ*_*s*_ as a way to signify different levels of ancientness (SI **Section** 5).

## 3 Results

### 3.1 Inferring Briggs parameters on simulated files

We evaluate the performance of the new regression method by comparing the inferred values of the Briggs parameters (*λ,δ*_*s*_,*δ*_*d*_ and *ν*) to the true values (from the simulations, SI **Section** 5), by normalizing the root mean square difference with the true value (NRMSE) as seen in **Figure** 2, denoting an error measurement, across the multiple replicates.

**Figure 2.**
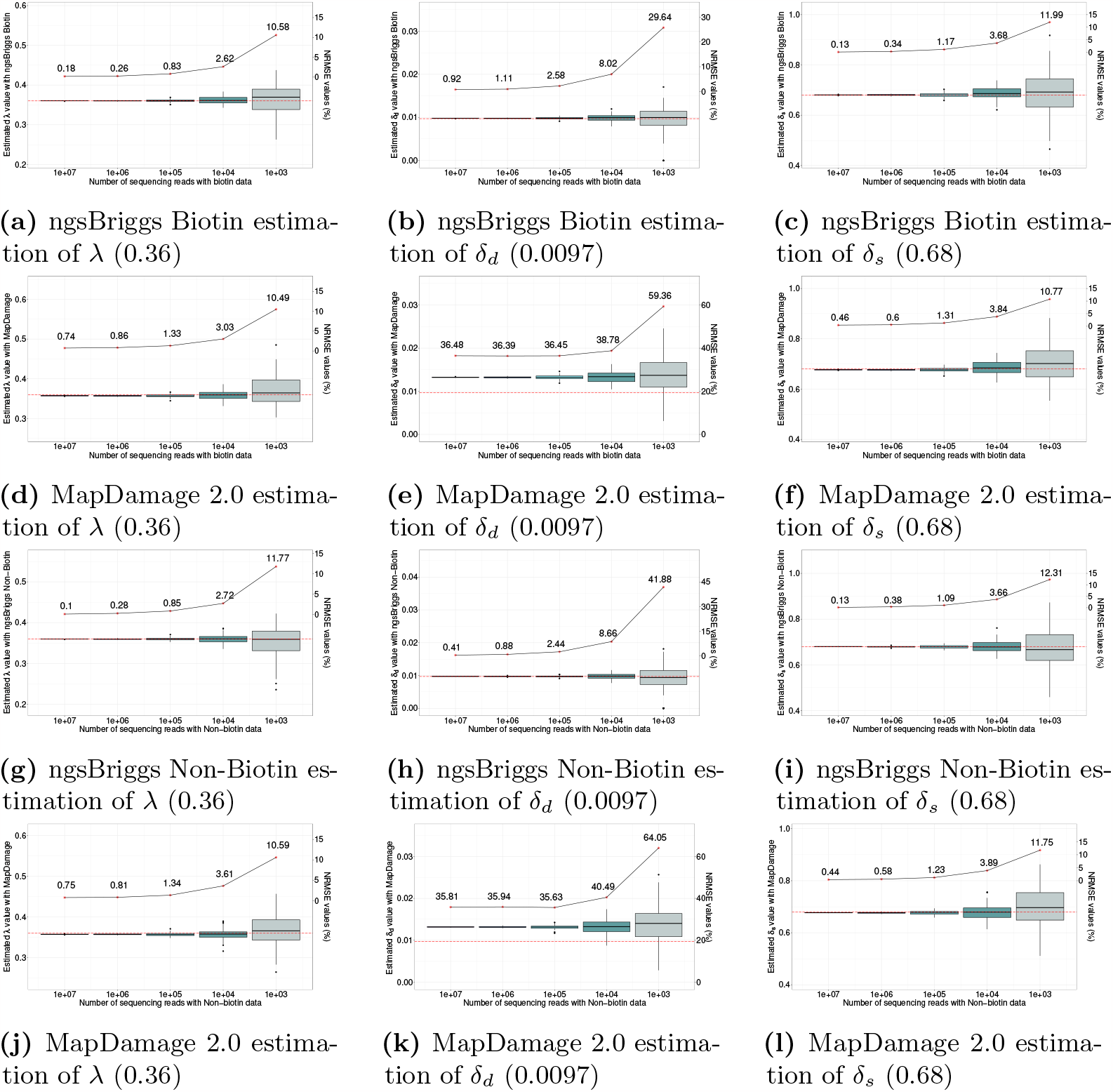
Each subfigure contains two separate y-axes representing different measurements. The left y-axis refers to the estimated values of *λ, δ*_*d*_, *δ*_*s*_ from the simulated data. The true parameter value is shown in each subfigure as a red horizontal dotted line and is also given in the figure caption. The right y-axis represents the NRMSE value. The x-axis shows the number of sequencing reads.

**Figure 3.**
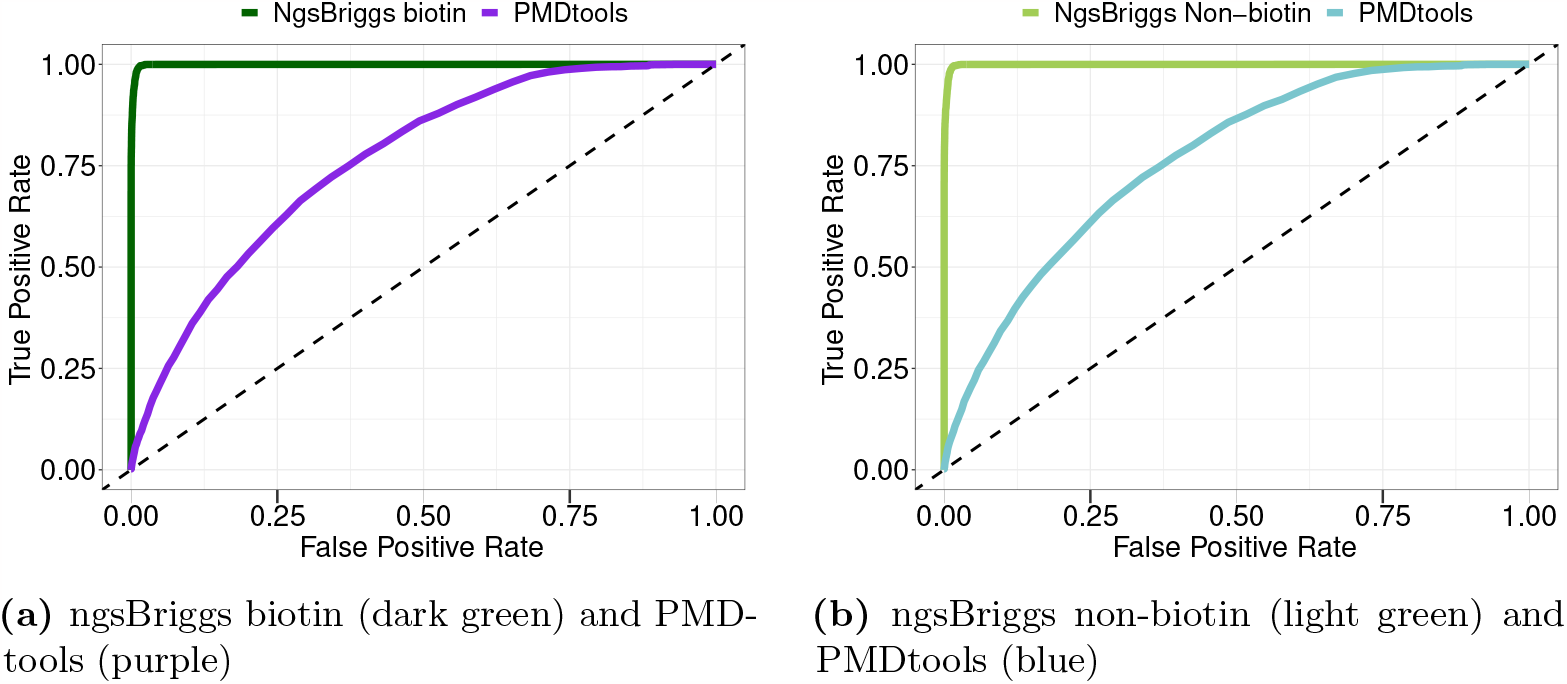
ROC curve illustrating the performance of the classification models, PMDtools and both ngsBriggs models.

We also compare the performance of ngsBriggs to mapDamage 2.0 under all scenarios. For both methods, we observe that with a lower number of reads (i.e., a lower sequencing depth), the interquartile range- and normalized root mean square error increases, signifying more uncertainty in the inferred parameters compared to the true deamination patterns.

Across all parameters, we observe that ngsBriggs for both the biotin (**Figure** 2a to 2c), and non-biotin simulated data (**Figure** 2g to 2i), exhibits a lower NRMSE compared to mapDamage 2.0 (**Figure** 2d to 2f and 2j to 2l). We observe the largest difference between the mapDamage 2.0 and ngsBriggs for the *δ*_*d*_ parameter (**Figure** 2b, and 2e), where mapDamage 2.0 exhibits lower accuracy and precision than ngsBriggs in all scenarios. The estimates of *ν* have the highest statistical uncertainty for both the biotin and non-biotin model (SI **Figure** 10 and 14).

### 3.2 Inferring Briggs parameters on empirical data

Scatter plots of inferred parameters from mapDamage 2.0 and ngsBriggs for all the ancient data (RISE, DA, VK and NEO) can be found in (SI **Figure** 16). We observed a clear correlation between the corresponding statistics for ngsBriggs and mapDamage 2.0. However, given that the data is empirical, we do not know the true values of the parameters, and hence cannot judge which method is more accurate.

Grouping all the ancient samples in different datasets, we conducted Jonckheere-Terpstra test [11] and observed significant increasing trends of ngsBriggs estimated *λ, δ*_*d*_ and *δ*_*s*_ as the radiocarbon date of the sample increases (see SI **Figure** 19 and 20). These observations may support the hypothesis that with increasing archaeological age, ancient specimens tend to accumulate deamination. However, there is likely to be a large contribution from location-specific preservation or project-specific treatment.

With regard to the efficiency of the tools, across all populations, we observed when creating a mismatch matrix that ngsBriggs is several magnitudes faster than both mapDamage 2.0 and PMDtools. When inferring the parameters, ngsBriggs is likewise magnitudes faster than mapDamage 2.0 (**Table** 1 and SI **Table** 4).

**Table 1.**
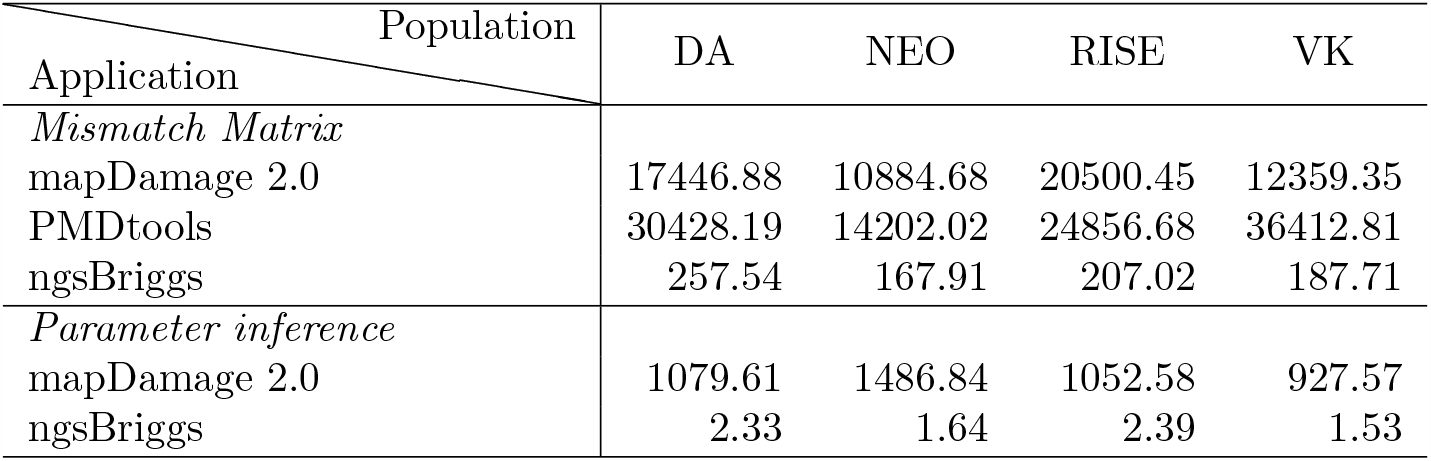
The first three lines represent the mean wall-clock running time used to generate the nucleotide matrices of 5’ C→T and 3’ G A deamination frequencies. The next two lines represent the wall clock running time of the Briggs parameter inference.

| Application | Population |  |  |  |
| --- | --- | --- | --- | --- |
|  | DA | NEO | RISE | VK |
| *Mismatch Matrix* |  |  |  |  |
| mapDamage 2.0 | 17446.88 | 10884.68 | 20500.45 | 12359.35 |
| PMDtools | 30428.19 | 14202.02 | 24856.68 | 36412.81 |
| ngsBriggs | 257.54 | 167.91 | 207.02 | 187.71 |
| *Parameter inference* |  |  |  |  |
| mapDamage 2.0 | 1079.61 | 1486.84 | 1052.58 | 927.57 |
| ngsBriggs | 2.33 | 1.64 | 2.39 | 1.53 |

### 3.3 Benchmarking in the presence of contamination

To measure the effect of contamination with modern human DNA, we compared parameter estimates of mapDamage 2.0, which assumes all reads are of ancient origin, to the ngsBriggs estimates, with- or without providing prior knowledge of contamination level (*ϵ*, SI **Section** 6.5).

When including contamination we observe stable estimates of *λ* across all investigated scenarios but have a tendency to be biased. We observe a significant impact from contamination on the three other estimated parameters *δ*_*d*_, *δ*_*s*_ and *ν*. MapDamage 2.0 estimates of *δ*_*d*_, *δ*_*s*_ decreases, due to its inability to take into account contamination from modern sources. When assuming no contamination with ngsBriggs we observe the same trend but allowing for *ϵ >* 0 it becomes possible to obtain essentially unbiased and accurate results (SI **Figure** 21 and 22).

### 3.4 Discrimination of ancient and modern reads

We assessed the performance of ngsBriggs when discriminating between ancient and modern reads by comparing them to PMDtools. This was done by using the Receiver Operating Characteristic (ROC) curve plotting the True Positive Rate (TPR) against the False Positive Rate (FPR) at different classification thresholds (**Figure** 3). Points above the diagonal signify better classification than could be obtained randomly. With a perfect classification, the line would have a TPR of 1 and FPR of 0 across all thresholds.

We benchmarked the tools as depicted in **Figure** 3 with the simulated files with 10% contamination, with the ancient component (SI **Section** 5.3) having the default deamination settings, similar to the ones used for benchmarking in **Figure** 2, provided by the original Briggs article [3] (*λ* : 0.36, *δ*_*d*_ : 0.0097, *δ*_*s*_ : 0.68, *ν* : 0.024). The sequence reads from the modern component (SI **Section** 5.3) follows *𝒩* (130,5) with an upper limit of 145, whereas the ancient sequence reads follow a constrained log-normal distribution(4,0.5) with a lower- and upper limit of 30 to 125 respectively. Additional deamination scenarios with variations in the *λ* and *δ*_*s*_ parameters are presented in SI **Section** 6.5.5.

We observe that PMDtools in most cases can distinguish between ancient DNA reads containing the PMD signal and human contaminants with sequencing error masking as C*→*T or G*→*A. However, the remaining ancient DNA fragments with a small fragment length but without deamination signal remain unclassified. As described in SI **Section** 4, ngsBriggs combines the strand length information and PMD signal to infer a potential sequence read length distribution for the contaminants. The ROC curve corresponding to the ngsBriggs method in **Figure** 3 is closer to the top-left corner. This indicates ngsBriggs almost perfectly discriminates between modern and ancient reads under the explored simulation conditions.

### 3.5 Runtime

When measuring the wall-clock time for the analyses of all empirical data sets, ngsBriggs was considerably faster than mapDamage 2.0, both when creating the mismatch matrix and when inferring the Briggs parameters (**Table** 1).

This time disparity is a consequence of the different frameworks of mapDamage 2.0, PMDtools and ngsBriggs. mapDamage 2.0 [11] combines Python and R and requires multiple independent steps to compute the aDNA-specific metrics (deamination frequency, statistical estimations, and visualizing the results). PMDtools processes and calculates the PMD using standard output produced by samtools [5] which parses the SAM file. The extra step of processing the samtools output accounts for most of the time difference. However, ngsBriggs computes all these metrics in one step without requiring multiple I/O operations.

## 4 ngsBriggs Implementation

The presented methods are implemented in the metadamage/metaDMG framework [18] as a standalone fast multi-threaded C/C++ function with htslib as a dependency. The code and documentation is available on https://github.com/lz398/metadamage briggs. We use the BFGS algorithm for parameter estimation in our regression model. For computing the mismatch matrix, we support the MD:Z tag in the AUX part of the samtools specification [14]. Additionally, the sub-functionality for computing the posterior probability that a read is ancient extends the AUX section of the alignment by adding a new non-standard tag AN:f:**AN**cient probability.

## 5 Discussion

The tool presented in this paper, ngsBriggs, introduces a novel approach for accurately quantifying the postmortem signal in reads sequenced by both older and modern sequencing platforms. Additionally, ngsBriggs represents a significant advancement in the authentication of ancient samples, as it combines the functionalities of mapDamage 2.0 and PMDtools for estimating damage parameters (*λ,δ*_*d*_,*δ*_*s*_ and *ν*) and computing posterior probabilities to discriminate ancient from modern reads. ngsBriggs accomplishes this with a significantly faster wall-clock running time compared to previous methods (**Table** 1). These factors make ngsBriggs highly suitable for large-scale high-throughput analyses of aDNA data.

By accounting for sequencing errors and true biological variation, ngsBriggs ensured accurate estimates of the Briggs parameters. Once the PMD parameters are inferred, researchers can effectively decontaminate and refine a given sample, removing reads with a low computed probability of being ancient. This enables researchers to make efficient use of the limited archaeological samples. During DNA extracting and library preparation, these finite ancient samples are inevitably destroyed, raising ethical implications. Although these ethical concerns might not necessarily impede scientific progress, our tool makes it possible for researchers to consider this, by mitigating the need for repeated and often destructive sampling thus contributing to the continued sustainable paleogenomics practice.

External estimates of contamination levels can mitigate the negative effect of exogenous DNA on the PMD signal, and ngsBriggs retains high accuracy during the inference of Briggs parameters and the assignment of read-specific contamination probabilities when such estimates are available.

For the simulated files, we observed greater accuracy for both ngsBriggs biotin and non-biotin models compared to mapDamage 2.0.

Our current implementation framework shows accurate inference of the Briggs parameters, and we show through extensive simulations, also using the Briggs model, that we are able to obtain essentially unbiased estimates of our statistics. An obvious limitation is that the postmortem damage signal is poorly understood, and the biochemical properties that are modeled directly by the four Briggs parameters will still pose an issue.

In the presence of contamination, we achieved more accurate estimates from ngsBriggs when providing a known contamination level. Hence, by leveraging ngsBriggs alongside tools like ContaMix [8], Schmutzi [19] or ANGSD [13], it is possible to estimate PMD features with high accuracy, even in the presence of significant contamination.

One potential issue with ngsBriggs arises in the presence of contamination when the assumption of distinguishable length distribution for the endogenous and modern contamination DNA is violated. When employing sequencing platforms like NovaSeq X with a 50 nucleotide cycle length, most sequenced reads, aDNA and potential modern contamination alike, will exhibit a similar length distribution, thus violating this assumption. While ngsBriggs may be successful in distinguishing some modern and ancient reads, the presented advantage as seen with the ROC curves would diminish.

As previously stated, ngsBriggs is implemented as part of a metaDMG framework to extend the mismatch matrix and the Briggs parameter inference functionalities to each taxonomic identifier (”taxid”) found in metagenomic data aligned to a reference database. By leveraging ngsBriggs within the metaDMG framework, researchers can authenticate and gain valuable insights into the taxonomic composition of ancient environmental DNA samples. This could enable researchers to delve deeper into the timeline of ancient samples and potentially gain new insight. This will broaden the scope of research in multiple fields, including paleogenomics, phylogenomics, metagenomics and historical ecology.

## Acknowledgements

We thank Dr. Lasse Vinner, Dr. Hugh McColl and Dr. Søren Overballe for many discussions regarding the laboratory protocols.

