## Supplementary document for "Revisiting the Briggs ancient DNA damage model: a fast regression method to estimate postmortem damage"

Supplementary Material

Lei Zhao, Rasmus Amund Henriksen, Abigail Ramsøe,
Rasmus Nielsen and Thorfinn Sand Korneliussen

6.5.1 Contamination rate's influence on parameter inference of 0.024, 0.36, 0.68 and 0.0097 36
6.5.2 Contamination rate's influence on parameter inference of 0.024, 0.7, 0.68 and 0.0097 . 36
6.5.3 Contamination rate's influence on parameter inference of 0.024, 0.1, 0.68 and 0.0097 . 37

**7 Future persepectives** **41**

### 1 Introduction

Ancient DNA (aDNA) exhibits distinct characteristics compared to modern DNA, including shorter fragment lengths, higher error rates, and deamination, with an increased frequency of the transition, cytosine to thymine ($C \rightarrow T$) in the 5' termini of the DNA fragments. These post-mortem damages (PMD) have been quantified by several studies developing methods assessing the deamination and its impact. In this study, we present the software ngsBriggs git commit *b5ed6b8* (`https://github.com/lz398/metadamage_briggs`, written in C/C++ with htslib as the sole dependency Bonfield *et al.*, 2021), designed to:

- Estimate four parameters ($\lambda$, $\delta_d$, $\delta_s$, $\nu$, as originally defined by Briggs *et al.* (2007), describing the deamination pattern while taking into account several scenarios, namely data with- or without sequencing errors or contamination.
- Compute a sequence read specific posterior probability to assert if the read originates from an ancient specimen or modern-day contaminant. With this computation, the sequencing reads can be discriminated and effectively used to decontaminate the aligned samples.

This program accomplishes this by modeling two different, yet related double-stranded DNA deamination models representing the original sequencing approach (Briggs *et al.*, 2007) and the contemporary one (Meyer and Kircher, 2010). This supplementary material first describes the overall procedure (**Figure** 1) and the distinction between the two deamination models (**Section** 2), hereafter referred to as the biotin and non-biotin model. Following the definition of both models, **Section** 3 and 4 document the mathematical modelling used for both features of ngsBriggs (as depicted in **Figure** 1). Following the methodological explanation, **Section** 5 gives examples of commands used for simulation, benchmarking and analysis throughout the work, and **Section** 6 presents extraordinary figures and tables to support our conclusions posed in the main manuscript, specifically that ngsBriggs, is not only capable of quantifying the PMD patterns with contemporary sequencing platforms, but also can maintain a greater accuracy when inferring the parameters and discriminating reads under various scenarios. Finally, in **Section** 7 we briefly put into context how ngsBriggs fit into the larger framework of metagenomic analysis of ancient DNA as mentioned in the main manuscript.

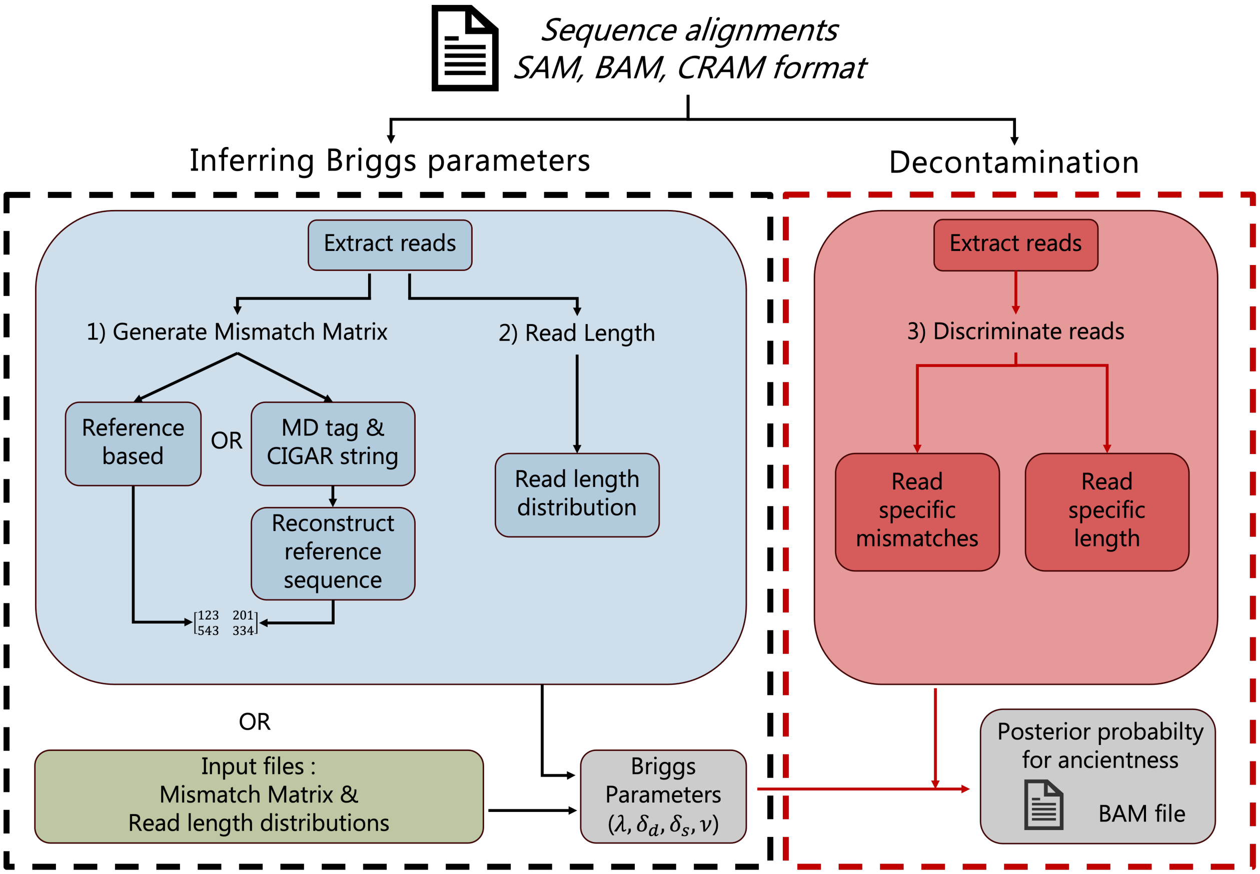

**Figure 1:** A workflow chart illustrating the different features of ngsBriggs. From the aligned sequence reads in a sequence Alignment/Map format (*.sam,.bam,.cram*) ngsBriggs can infer the Briggs parameters and utilize these to decontaminate samples. The **black** dotted box highlights two approaches to infer the parameters of the PMD signal (see **Section** 2 and 3) by generating a mismatch matrix (with a simplistic visualization shown, with **Table** 2 showing an accurate version) and read length distribution. The first approach uses the aligned reads (**blue** box), which itself can identify mismatches from a provided reference genome or the information stored within the MD tag and CIGAR string. The second approach infers the parameters from input files containing the mismatch matrix and a length distribution (**green** box). The **red** dotted line highlights the steps, ngsBriggs utilize to decontaminate samples (see **Section** 4), by generating sequence read-specific mismatches and lengths, which in conjunction with the inferred parameters are employed to compute read-specific posterior probabilities for ancientness. With the grey boxes representing the output.

#### 2 Overview of aDNA Deamination Models

In this section, mathematical formulae are derived for both the biotin- and non-biotin models. Both models assume that the observed deamination pattern is a joint effect of 5' overhangs (both left and right) caused by aDNA fragmentation, aDNA deaminations in the double- and single-stranded regions, the possible nick placement and the PCR library preparation protocols. As described in the main text, the biotin and non-biotin models differ in the library preparation protocols, which results in different templates for the downstream PCR process. In the biotin model, only the original ancient strands will serve as the PCR templates, while in the non-biotin model, both the original strands and their reverse-complement can act as the templates (visualized in **Figure** 2). The different PCR templates will affect the observed deamination patterns and the form of the likelihood functions for the estimation of the parameters, which we will discuss in **Section** 3. In this section, we will focus on the model features of aDNA fragmentation, aDNA deamination, and nick placement, which both models share.

##### 2.1 Notations

Before going into the details, we need to introduce the notations adopted in the models. They are listed in **Table** 1.

| | |
|---|---|
| $\lambda$ | Geometric distribution parameter related to the length |
| $\delta_d$ | Deamination rate per nucleotide site in the double strand region |
| $\delta_s$ | Deamination rate per nucleotide site in the single strand region |
| $\nu$ | nick rate per nucleotide site |
| $O_l$ | The length of the left 5' overhang |
| $O_r$ | The length of the right 5' overhang |
| $\epsilon_n$ | Sequencing error at position $n$ |
| $N_{n,\mathrm{X}\to\mathrm{Y}}$ | Counts of X(ref) $\to$ Y(read) from the mismatch matrix. |
| $f_{\mathrm{X}\to\mathrm{Y}\mid\mathrm{X}}(n, L; m)$ | Frequency of nucleotide change X(ref)$\to$Y(read) with model $m$ at position $n$ |
| $\chi_n^k$ | Indicator functions at position $n$ of focal strand $k$ |
| $\chi_{n,i}^k$ | Indicator functions at position $n$ within region $i$ of the focal strand $k$ |
| $\delta_{\mathrm{eff},n,\mathrm{X}\to\mathrm{Y}}$ | Theoretical deamination rate of X$\to$Y at position $n$ |
| $P_k^a$ | The posterior probability of strand $k$ being ancient |
| $f_L$ | Probability of fragment having length $L$ |

**Table 1:** Notations for different model variables

##### 2.2 5' Overhangs

Within the aDNA samples, DNA molecules exist in the form of short fragments. As shown in **Figure** 2 and 3, such fragments have possible double-stranded regions and single-stranded regions. The double-stranded library preparation only considers the reads with double-stranded regions. The single-stranded regions are assumed to possibly appear at both ends of the aDNA fragment, to which we refer as **overhangs**. If an overhang contains a 5' end of one strand in the focal fragment, we call it **5' overhang** (See Quadrant I of **Figure** 2 and 3 where both strands of the fragment have 5' overhangs), otherwise, if an overhang happens to have a 3' end of a strand, we call it **3' overhang** (See Quadrant III of **Figure** 2 and 3 where both strands have 3' overhangs). In ancient materials, deamination process accumulates observed C$\to$T substitutions, and the deamination rate in the single-stranded region, $\delta_s$ is believed to be much higher than the corresponding rate, $\delta_d$, in the double-stranded region, since the double-stranded region is more stable. The difference between $\delta_s$ and $\delta_d$ is assumed to be a driving factor to form the observed deamination pattern (Briggs *et al.*, 2007): As discussed in the main text, the end-repair treatment removes any possible 3' overhang, and retains the 5' overhangs as the only single-stranded aDNA regions that contribute to the downstream analyses (as shown in **Figure** 2 and 3). We further conceptually define the **left 5' overhang** of a specific strand as the 5' overhang on this strand; while its **right 5' overhang** denotes the 5' overhang on the other strand of the same aDNA fragment. The distribution of the length of the left 5' overhang, $O_l$ and that of the right 5' overhang, $O_r$ are assumed to be identical and as follows,

$$\mathbf{P}\left(O_l = l \,|\lambda, L\right) = \mathbf{P}\left(O_r = l \,|\lambda, L\right) = \frac{1}{2}\frac{\lambda\left(1-\lambda\right)^l}{1-\left(1-\lambda\right)^{L-1}} + \frac{1_{l=0}}{2}. \tag{1}$$

The above probability function can be interpreted as a weighted mixture of a Kronecker delta and a rescaled geometric distribution. The Kronecker delta part represents the case when there is originally a 3' overhang at the focal end of the fragment (Since the 3' overhang will be removed during the end-repair treatment, this case can be viewed as forming a 5' overhang of the length 0). In contrast, the rescaled geometric distribution part is the situation where the focal end is a 5' overhang, whose length is limited in the range $[0, L-2]$. The chances that the focal end is a 5' overhang or a 3' end are the same and equal 50%.

We further assume the total length of left 5' overhang $O_l$ and right 5' overhang $O_r$ can not exceed $L-2$, the joint distribution of $O_l$ and $O_r$ can then be defined as follows,

$$\mathbf{P}\left(O_l = l, O_r = r \,|\lambda, L\right) = \frac{\mathbf{P}\left(O_l = l \,|\lambda, L\right)\mathbf{P}\left(O_r = r \,|\lambda, L\right)}{S}, \tag{2}$$

where $S$ is the scaling factor to standardise the probability and can be calculated as follows,

$$S = \sum_{l+r\leq L-2} \mathbf{P}\left(O_l = l \,|\lambda, L\right)\mathbf{P}\left(O_r = r \,|\lambda, L\right) = \frac{3}{4} + \frac{1}{4}\frac{1-\left(1-\lambda\right)^{L-1} - (L-1)\lambda\left(1-\lambda\right)^{L-1}}{\left[1-\left(1-\lambda\right)^{L-1}\right]^2}. \tag{3}$$

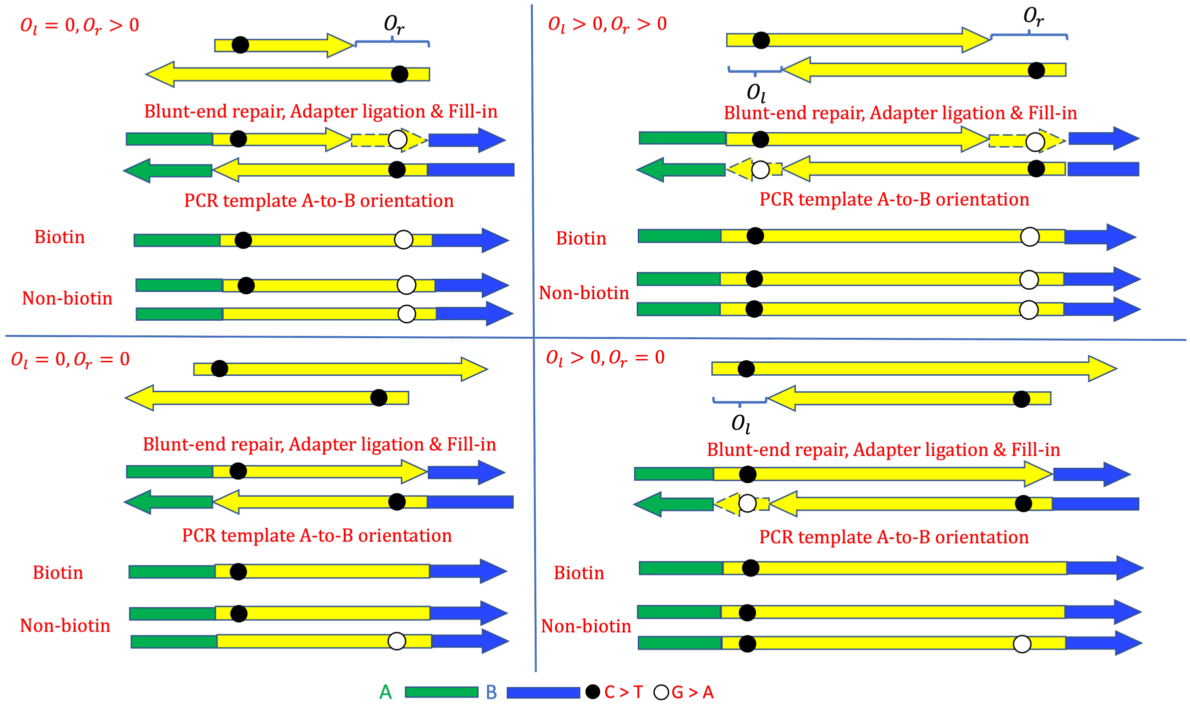

**Figure 2:** An illustration of postmortem damage patterns induced only by the overhangs with different parameter settings: The plots are arranged in a quadrant sense. Quadrants I, II, III and IV represent the situations when the lengths of left and right 5' overhangs satisfy $O_l, O_r > 0$ ; $O_r > O_l = 0$ ; $O_l = O_r = 0$, and $O_l > O_r = 0$, respectively. 3' overhangs will be discarded, and in the cases where 5' overhangs exist, their complements are generated such that the corresponding single-stranded parts of the ancient fragments will be repaired. Adaptors A and B are then assumed to randomly attach to the ends of the fragments. Only the fragments with both adaptors A and B attached contribute to the downstream PCR. For simplicity, we only show the results when adaptors A and B are attached to the left and the right of the focal fragment, correspondingly. But equal chances can be that adaptors A and B are attached the other way around. In the biotin model, only the original ancient strands with A-to-B orientation serve as the PCR templates; while in the non-biotin model, the A-to-B original ancient strands and the reverse-complement of the B-to-A original ancient strands will both serve as the PCR templates.

#### 2.3 Nicks Placement

The deamination process described above can only explain the true damage signals accumulated in the aDNA samples, i.e., C→T. The complementary changes, i.e., G→A, which are also intensively observed, will be introduced by the T4 DNA polymerase end-repair treatment. Without nicks in the aDNA fragment, the end-repair treatment will introduce the complement G→A on the single-stranded regions, or more specifically, on the right 5' overhangs (see **Figure** 2). Nicks can help to place G→A substitutions on the double-stranded regions (see **Figure** 3). The terminology nick is defined as the small crack on the backbone of the aDNA molecules, which is caused by the breakup of the phosphodiester bond between adjacent nucleotides. Nicks are also consequences of postmortem damage. In theory, nicks can occur at each position of an aDNA fragment, however, the nicks in the single-stranded regions will effectively shorten the overhangs. We thus focus on the nicks in the double-stranded regions. As shown in **Figure** 3, during the end-repair procedure, any sequence downstream of the first nick of the focal strand will be replaced by a new one generated in accordance with its opposite strand, which can lead to complementary G→A in the double-stranded regions.

In the following parts, we split the descriptions of nick placement for the biotin and non-biotin models.

The reason for this split is in the biotin model, only one strand of each fragment will contribute to the downstream analyses, while in the non-biotin model, both strands matter for the downstream analyses. In the biotin model, we only need to care about the nick on a randomly chosen strand, but in the non-biotin model, the nick positions on both strands of the same fragment may interact with each other.

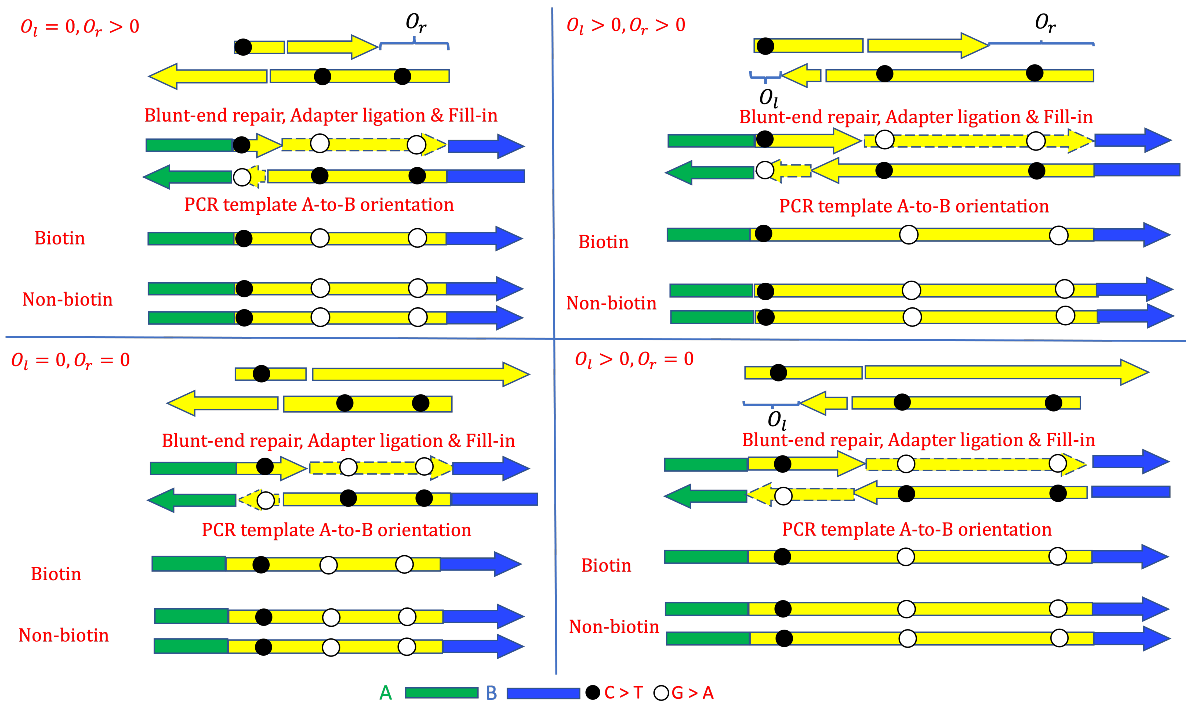

**Figure 3:** An illustration of postmortem damage patterns induced by the overhangs and nicks with different parameter settings. The plots arrangement and the assumptions of 5' overhangs and adaptors are as addressed in **Figure** 2. The nicks on both strands satisfy 1, on the double-strand region, 2, there is no other nick upstream of the focal nick position on both strands. Any nucleotides downstream of the nicks will be filled up by the reverse-complement of the other strand.

**Figure** 4 is similar to **Figure** 3 except that only observing a nick on one of the two strands.

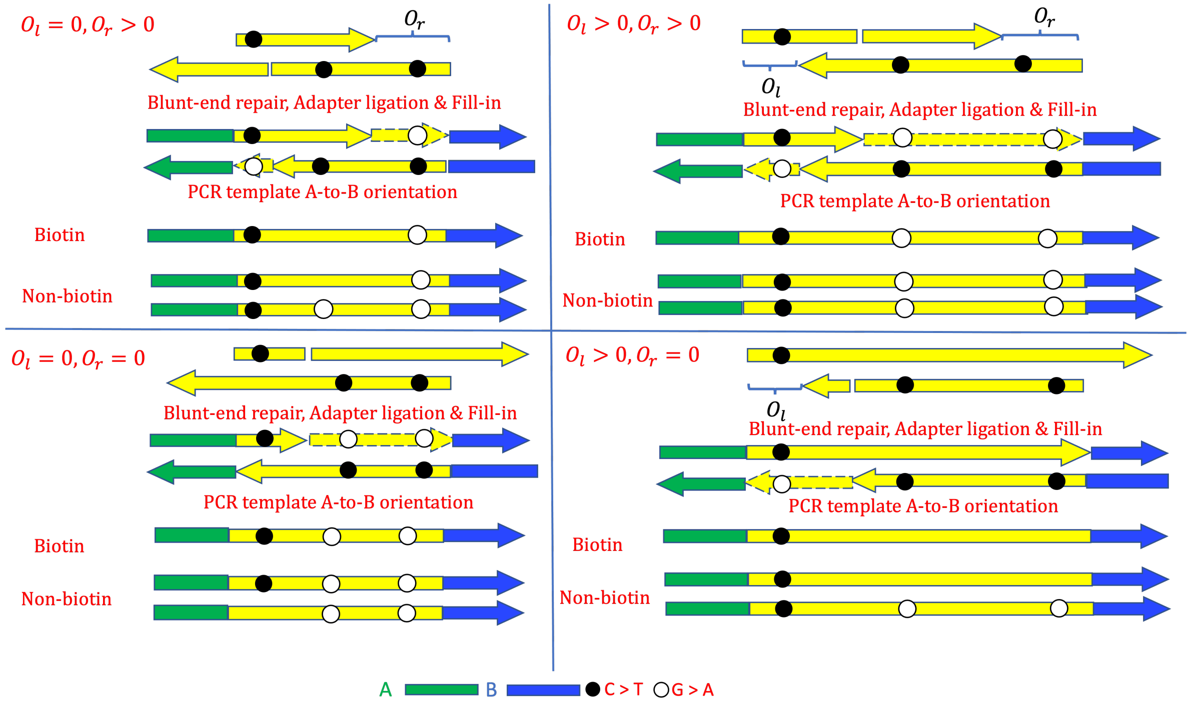

**Figure 4:** An illustration of postmortem damage patterns only induced by the overhangs and a single nick with different parameter settings The plots arrangement and the assumptions of 5′-overhangs and adaptors are as addressed in **Figure** 2.

##### 2.3.1 Biotin model

The nick placement model of the biotin model is based on the supplement Information of (Briggs *et al.*, 2007). The chance that a randomly chosen strand (with the length of its double strand region $L_{ds} = L - l - r$) from the raw fragments before PCR is a No-nick fragment is

$$\mathrm{P}\left(\text{No-nick}\right) = (1-\nu)^{L_{ds}-1}\,, \tag{4}$$

and the probability that the first nick of this randomly chosen fragment is at double-stranded position $n_{ds}$ $(1 \le n_{ds} < L_{ds})$, is

$$\mathrm{P}\left(n_{ds}\right) = \nu\,(1-\nu)^{L_{ds}-2}\,, \tag{5}$$

where the probability is calculated based on the assumption that no nick should occur upstream of this nick in the sense of both strands of the focal fragment.

As shown in **Figure** 5 and addressed in (Briggs *et al.*, 2007) and above, any sequence downstream of the first nick on the focal strand will be replaced by a new one generated in accordance with its complementary strand, and if the focal strand carries no nick, its part of the right 5' overhang will also be filled up by the new sequence generated according to the opposite strand. For simplicity, the No-nick fragment is effectively viewed as the fragment whose first nick hits the last position $L_{ds}$ of the double-stranded region. Hence any strand in the biotin model will have an effective nick position and the rescaled probability of nick position in the biotin model can be given as follows,

$$\mathrm{P}\left(n_{ds}\,|\nu, L_{ds}\,\right) = \begin{cases} \frac{\nu}{(L_{ds}-1)\nu+(1-\nu)}, & 1 \le n_{ds} < L_{ds}, \\ \frac{1-\nu}{(L_{ds}-1)\nu+(1-\nu)}, & n_{ds} = L_{ds}, \end{cases} \tag{6}$$

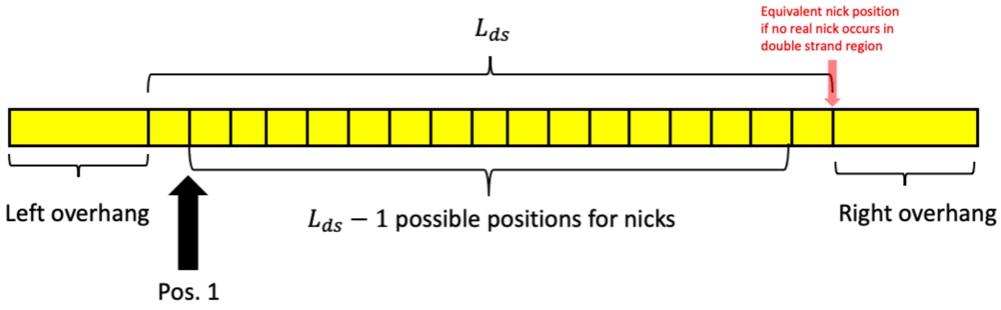

**Figure 5:** Different from the previous illustrations, this figure represent the original double-stranded fragment, since this is the only region where its possible to place a nick. As nick occurs between nucleotide sites, only $L_{ds}-1$ possible positions for nick in one strand. If the nick has not occurred within the $L_{ds}-1$ positions, one could equivalently view a nick occurring at the first position of the right overhang. This figure also shows if the nick on one strand is fixed at position $n_{ds}$, all the possible nick positions $m_{ds}$ on the other strand according to the models' assumptions.

##### 2.3.2 Non-biotin model

In the non-biotin model, we will focus on both strands of the aDNA fragment. The marginal distribution of the nick position $n_{ds}$ on a randomly chosen strand is given by **Equation** 6. Given $n_{ds}$, we define the conditional probability of the nick position on the other strand, $m_{ds}$, as a geometric distribution as follows,

$$\mathrm{P}\left(m_{ds}\,|\nu, L_{ds}, n_{ds}\right) = \begin{cases} (1-\nu)^{n}, & m_{ds} = L_{ds}, \\ \nu\,(1-\nu)^{m_{ds}+n-L_{ds}}, & L_{ds}-n \leq m_{ds} < L_{ds}, \\ 0, & 1 \leq m_{ds} < L_{ds}-n, \end{cases} \tag{7}$$

where $n = \min\left(n_{ds}, L_{ds}-1\right)$, $m_{ds}$ and $n_{ds}$ are counted from their own strands' 5' end.

The reason that $\mathrm{P}\left(m_{ds}\,|\nu, L_{ds}, n_{ds}\right) = 0$ when $1 \leq m_{ds} < L_{ds} - n$ in **Equation** 7 is based on the assumption addressed before "no nick should occur upstream of this nick in the sense of both strands of the focal fragment" (See **Figure** 6). Henriksen *et al.* (2023) adopted **Equations** 6 and 7 to place the nicks in the non-biotin model simulations.

Because of **Equations** 6 and 7, we can also calculate the marginal distribution of the nick position $m_{ds}$ on the corresponding opposite strand.

$$\mathrm{P}\left(m_{ds}\,|\nu, L_{ds}\right) = \sum_{n_{ds}} \mathrm{P}\left(m_{ds}\,|\nu, L_{ds}, n_{ds}\right) \mathrm{P}\left(n_{ds}\,|\nu, L_{ds}\right) \tag{8}$$

$$= \begin{cases} \frac{\nu}{(L_{ds}-1)\nu+(1-\nu)}, & 1 \leq m_{ds} < L_{ds} \\ \frac{1-\nu}{(L_{ds}-1)\nu+(1-\nu)}, & m_{ds} = L_{ds} \end{cases}, \tag{9}$$

which indicates $m_{ds}$ and $n_{ds}$ share the identical marginal distribution. And this conclusion indicates that there is no need to distinguish the nick position distributions on both strands of the same fragment, which helps to build the regression likelihoods of the non-biotin model in **Section** 3.1.2.

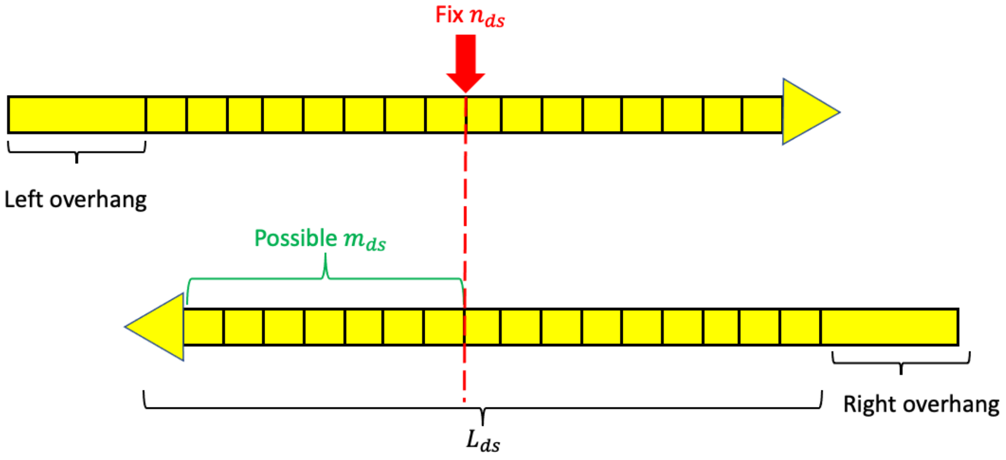

**Figure 6:** Possible nick placement: In both models, nicks are assumed to be effectively put between adjacent nucleotide sites in the double strand region. There are $L_{ds} - 1$ possible nick positions on each strand. In accordance with the models' assumptions, we only focus on the first nicks on both strands, and if there is a nick on one strand, there should not be nicks found upstream of the nick position on the other strand. If no nick occurs, one could view a nick being put at the first position after the $L_{ds} - 1$ possible nick positions.

### 3 Regression Likelihood

The previous methods inferring the Briggs model's parameters often go through each read $i$ and each position $j$, and try to optimize a full log-likelihood function of the following format, e.g., (Ginolhac *et al.*, 2011)

$$l = \sum_{i=1}^{\#\text{read}} \sum_{j=1}^{\#\text{pos.}_i} l_{ji},$$

To maximize such a function, advanced technologies like Markov chain Monte Carlo (MCMC) are applied but the previous methods still suffer from relatively large computational time and memory especially when a large quantity of data is supplied. We thus developed a method that first theoretically calculates the postmortem damage rate (C $\to$ T and G $\to$ A rates) at a specific set of cyclic positions (first few positions near the 5' end and 3' end) of aDNA fragments, and then utilizes a multinomial regression to estimate the Briggs parameters.

The proposed regression method requires a summarised mismatch matrix as follows.

| Dir. | Pos. | AA | AC | AG | AT | CA | CC | CG | CT | GA | GC | GG | GT | TA | TC | TG | TT |
|---|---|---|---|---|---|---|---|---|---|---|---|---|---|---|---|---|---|
| 5' | 1 | 295783 | 70 | 59 | 67 | 48 | 158603 | 43 | 46031 | 41 | 38 | 204183 | 35 | 76 | 59 | 77 | 294787 |
| 5' | 2 | 295110 | 38 | 41 | 38 | 44 | 174339 | 43 | 29978 | 56 | 50 | 204314 | 36 | 64 | 52 | 58 | 295739 |
| 5' | 3 | 294837 | 104 | 92 | 101 | 79 | 184374 | 81 | 19874 | 105 | 62 | 204037 | 62 | 144 | 182 | 155 | 295711 |
| | | | | | | | ... | | ... | ... | | | | | | | |
| 3' | 1 | 295377 | 102 | 94 | 98 | 59 | 204295 | 61 | 598 | 44568 | 79 | 159001 | 66 | 101 | 94 | 97 | 295310 |
| 3' | 2 | 295561 | 117 | 108 | 102 | 58 | 203462 | 67 | 675 | 29313 | 59 | 175467 | 49 | 108 | 98 | 90 | 294666 |
| 3' | 3 | 295221 | 94 | 97 | 92 | 50 | 203909 | 59 | 749 | 19044 | 67 | 185258 | 60 | 104 | 79 | 85 | 295032 |
| | | | | | | | ... | | ... | ... | | | | | | | |

**Table 2:** An example of the mismatch matrix, the column AC denotes the counts of nucleotides across the focal sample whose reference are As but the reads show Cs at different cyclic positions. The counts of X (ref) to Y (read) at cyclic position $n$ in this table will be denoted as $N_{n,\text{X}\to\text{Y}}$ in **Section** 3.

#### 3.1 General Results for Both Models

In both models (biotin and non-biotin models), The observed postmortem damage rate (C $\to$ T or G $\to$ A rate) at a specific cyclic position depends on the spatial relationships of this position, potential nick, and both 5' overhangs. Considering one of the original strands in both models, we classify such spatial relationships into four different types:

1. The focal position $n$ is within the double-strand region and downstream of the possible first nick on this strand;

2. the focal position $n$ is within the right 5' overhang region;

3. the focal position $n$ is within the double-strand region and upstream of the possible first nick (this also includes the case where there is no nicks);

4. the focal position $n$ is within the left 5' overhang region.

We define the probabilities of the above four mutually exclusive types of spatial relationships as $p_1(n;L)$, $p_2(n;L)$, $p_3(n;L)$, and $p_4(n;L)$, respectively. If the focal position $n$, nick position, and the 5' overhangs have relation type 3 (or 1), the observed C $\to$ T (or the complementary G $\to$ A) will have a rate same as the true deamination within the double-strand region, $\delta_d$; while if the spatial distribution of $n$, nick position, and the 5' overhangs are of the relation type 4 (or 2), the observed C $\to$ T (or the complementary G $\to$ A) will have a rate $\delta_s$, which is the true deamination rate on the single-strand region of the ancient DNA fragment.

Based on the mechanisms described in **Section** 2 and the assumption that the aDNA fragments are of the fixed length $L$, we have $p_1(n;L)$calculated as follows,

$$\begin{aligned}
p_1(n;L) =& \sum_{l=0}^{n-1}\sum_{r=0}^{L-n} \mathbf{P}\left(O_l = l, O_r = r \,|\lambda, L, O_l + O_r \leq L-2\,\right) \mathbf{P}\left(p_n \leq n-1 \,|\nu, \text{seq}, O_l = l, O_r = r\,\right) \\
=& C\left[1-(1-\lambda)^{L-1}+\lambda\right]^2 \frac{(n-1)\nu}{(L-1)\nu+(1-\nu)} \\
&+C\lambda\left[1-(1-\lambda)^{L-1}+\lambda\right]\sum_{l=1}^{n-1}(1-\lambda)^l \frac{(n-l-1)\nu}{(L-1-l)\,\nu+(1-\nu)} \\
&+C\lambda\left[1-(1-\lambda)^{L-1}+\lambda\right]\sum_{r=1}^{L-n}(1-\lambda)^r \frac{(n-1)\nu}{(L-1-l)\,\nu+(1-\nu)} \\
&+C\lambda^2\sum_{l=1}^{n-1}\sum_{r=1}^{L-n}(1-\lambda)^{l+r} \frac{(n-l-1)\nu}{(L-1-l-r)\,\nu+(1-\nu)}
\end{aligned} \tag{10}$$

where $n$ is the focal nucleotide position counted from the 5' end. $p_n$ represents the first nick position within the fragment, and

$$C = \frac{1}{4S\left[1-(1-\lambda)^{L-1}\right]^2}, \tag{11}$$

the definition of $C$ is only for simplification, $S$ is defined in previous **Equation** 3, and the final results of **Equation** 10 are splitted into four lines representing the cases where $l = r = 0$, $l > r = 0$, $r > l = 0$ and $l$, $r > 0$, respectively.

Similarly, $p_2$, $p_3$, and $p_4$ are given as follows,

$$\begin{aligned}
p_2(n;L) &= \sum_{r=L-n+1}^{L-2} \sum_{l=0}^{L-2-r} \mathbf{P}\left(O_l = l, O_r = r \,|\lambda, L, O_l + O_r \le L-2\right) \\
&= C \sum_{r=L-n+1}^{L-2} \lambda\left(1-\lambda\right)^r \left[1-\left(1-\lambda\right)^{L-1}+\lambda\right] + C \sum_{r=L-n+1}^{L-2} \sum_{l=1}^{L-2-r} \lambda^2 \left(1-\lambda\right)^{r+l} \\
&= C\left[2-\left(1-\lambda\right)^{L-1}\right]\left[\left(1-\lambda\right)^{L-n+1}-\left(1-\lambda\right)^{L-1}\right] - C(n-2)\lambda(1-\lambda)^{L-1}, \\
p_3(n;L) &= \mathbf{P}\left(n \text{ in ds}\right) - p_1(n;L) \\
&= C\left[2-\left(1-\lambda\right)^{L-1}-\left(1-\lambda\right)^{n}\right]\left[2-\left(1-\lambda\right)^{L-1}-\left(1-\lambda\right)^{L-n+1}\right] - C\lambda^2\left(1-\lambda\right)^{L-1} \\
&\quad - 1_{n=1 \text{ or } L} C\lambda\left(1-\lambda\right)^{L-1}\left[1-\left(1-\lambda\right)^{L-1}\right] - p_1(n;L), \\
p_4(n;L) &= \sum_{l=n}^{L-2} \sum_{r=0}^{L-2-l} \mathbf{P}\left(O_l = l, O_r = r \,|\lambda, L, O_l + O_r \le L-2\right) \\
&= C\left[2-\left(1-\lambda\right)^{L-1}\right]\left[\left(1-\lambda\right)^{n}-\left(1-\lambda\right)^{L-1}\right] - C(L-n-1)\lambda\left(1-\lambda\right)^{L-1}.
\end{aligned}$$

In the following context where all fragments have the same fixed length $L$, we will write $p_j(n;L)$ as $p_j(n)$ ($j = 1,2,3,4$) for short.

##### 3.1.1 Biotin Model with Fixed Length $L$

In the biotin model, only the strands that belong to the original ancient fragments can serve as the PCR templates, we thus can calculate the probability of observing a T (or A) given reference is a C (or G) at position $n$ of the PCR outputs under the biotin model, denoted as $\delta_{\text{eff},n,\text{G}\to\text{A}}$ and $\delta_{\text{eff},n,\text{C}\to\text{T}}$. $\delta_{\text{eff},n,\text{G}\to\text{A}}$ and $\delta_{\text{eff},n,\text{C}\to\text{T}}$ can be interpreted as the frequencies of the observed C→T and G→A at the cyclic position $n$, given the reference is C and G, correspondingly.

$$\delta_{\text{eff},n,\text{G}\to\text{A}} = \delta_s p_2(n) + \delta_d p_1(n), \tag{12}$$

$$\delta_{\text{eff},n,\text{G}\to\text{G}} = 1 - \delta_{\text{eff},n,\text{G}\to\text{A}}, \tag{13}$$

$$\delta_{\text{eff},n,\text{C}\to\text{T}} = \delta_s p_4(n) + \delta_d p_3(n), \tag{14}$$

$$\delta_{\text{eff},n,\text{C}\to\text{C}} = 1 - \delta_{\text{eff},n,\text{C}\to\text{T}}. \tag{15}$$

Here $\delta_{\text{eff},n,\text{C}\to\text{C}}$ and $\delta_{\text{eff},n,\text{G}\to\text{G}}$ represent the probabilities of the match of the observed nucleotide and its reference under the biotin model at the position $n$, conditional on the corresponding reference is C or G.

##### 3.1.2 Non-biotin Model with Fixed Length $L$

Unlike the biotin model where all the selected PCR templates are the original ancient strands, in the non-biotin model, the PCR templates can be either the original ancient strands or the reverse-complement of the original ancient strands. Each of the cases has a probability of 50%. Take a G → A change at position $n$ on the double-strand part of a PCR template as an example: In the non-biotin model, it can be introduced by either the case that the position $n$ of an original strand bears a complementary deamination, G → A, and satisfies the spatial relation type 1 and the original strand directly plays a role as a PCR template or the case that position $L-n+1$ of an original strand harbours a true deamination, C → T, and satisfies the spatial relation type 3 and the original strand's reverse-complement serves as a PCR template. C → T on the double-strand part of a PCR template, and G $\to$ A or C $\to$ T on the single-strand part of a PCR template can also be interpreted in the similar way. We hence calculate the observed frequency of T (or A) given reference is a C (or G) at position $n$ of the PCR outputs under the non-biotin model, denoted as $\tilde{\delta}_{\text{eff},n,\text{G}\to\text{A}}$ and $\tilde{\delta}_{\text{eff},n,\text{C}\to\text{T}}$,

$$\tilde{\delta}_{\text{eff},n,\text{G}\to\text{A}} = \frac{\delta_s}{2}\left[p_2(n) + p_4(L-n+1)\right] + \frac{\delta_d}{2}\left[p_1(n) + p_3(L-n+1)\right], \tag{16}$$

$$\tilde{\delta}_{\text{eff},n,\text{G}\to\text{G}} = 1 - \tilde{\delta}_{\text{eff},n,\text{G}\to\text{A}}, \tag{17}$$

$$\tilde{\delta}_{\text{eff},n,\text{C}\to\text{T}} = \frac{\delta_s}{2}\left[p_2(L-n+1) + p_4(n)\right] + \frac{\delta_d}{2}\left[p_1(L-n+1) + p_3(n)\right], \tag{18}$$

$$\tilde{\delta}_{\text{eff},n,\text{C}\to\text{C}} = 1 - \tilde{\delta}_{\text{eff},n,\text{C}\to\text{T}}, \tag{19}$$

where $\tilde{\delta}_{\text{eff},n,\text{C}\to\text{C}}$ and $\tilde{\delta}_{\text{eff},n,\text{G}\to\text{G}}$ represent the probabilities of the match of the observed nucleotide and its reference in the non-biotin model, conditional on the corresponding references are given.

It is obvious that the observed deamination pattern of the non-biotin model is more symmetric than that of the biotin model since from the above equations, we have the following relationship,

$$\tilde{\delta}_{\text{eff},n,\text{C}\to\text{T}} = \tilde{\delta}_{\text{eff},L-n+1,\text{G}\to\text{A}}, \tag{20}$$

while the biotin model does not have the same property.

###### 3.1.3 Models with Variant Lengths

It is highly unlikely that all the reads in an ancient sample have the same length, we now consider a more realistic sample of aDNA fragments with a length distribution $f_L$ ($f_L$ corresponds to the probability that a fragment has the length $L$, and $\sum_L f_L = 1$) under both models. The observed G $\to$ A and C $\to$ T rates at position $n$ can be viewed as $f_L$ weighted average of the results in **Subsections** 3.1.1 or 3.1.2, and such rates under the biotin model with a known length distribution $f_L$ are as follows,

$$\delta_{\text{eff},n,\text{G}\to\text{A}} = \delta_s \sum_L p_2(n;L) f_L + \delta_d \sum_L p_1(n;L) f_L, \tag{21}$$

$$\delta_{\text{eff},n,\text{G}\to\text{G}} = 1 - \delta_{\text{eff},n,\text{G}\to\text{A}}, \tag{22}$$

$$\delta_{\text{eff},n,\text{C}\to\text{T}} = \delta_s \sum_L p_4(n;L) f_L + \delta_d \sum_L p_3(n;L) f_L, \tag{23}$$

$$\delta_{\text{eff},n,\text{C}\to\text{C}} = 1 - \delta_{\text{eff},n,\text{C}\to\text{T}}, \tag{24}$$

while the corresponding rates under the non-biotin model with a known length distribution $f_L$ are given as,

$$\tilde{\delta}_{\text{eff},n,\text{G}\to\text{A}} = \frac{\delta_s}{2} \sum_L \left[p_2(n;L) + p_4(L-n+1;L)\right] f_L + \frac{\delta_d}{2} \sum_L \left[p_1(n;L) + p_3(L-n+1;L)\right] f_L, \tag{25}$$

$$\tilde{\delta}_{\text{eff},n,\text{G}\to\text{G}} = 1 - \tilde{\delta}_{\text{eff},n,\text{G}\to\text{A}}, \tag{26}$$

$$\tilde{\delta}_{\text{eff},n,\text{C}\to\text{T}} = \frac{\delta_s}{2} \sum_L \left[p_2(L-n+1;L) + p_4(n;L)\right] f_L + \frac{\delta_d}{2} \sum_L \left[p_1(L-n+1;L) + p_3(n;L)\right] f_L, \tag{27}$$

$$\tilde{\delta}_{\text{eff},n,\text{C}\to\text{C}} = 1 - \tilde{\delta}_{\text{eff},n,\text{C}\to\text{T}}, \tag{28}$$

###### 3.1.4 Models with Sequencing Errors and Modern Contamination

Sequencing errors and modern contamination can also be incorporated into the models.

The cyclic position-specific sequencing errors will be estimated based on the mismatch matrix. To avoid the interference of the deamination, we use reference nucleotide A and T's changes to calculate the errors, i.e.,

$$\epsilon_n = \frac{\sum_{\mathrm{X}\neq\mathrm{A}} N_{n,\mathrm{A}\to\mathrm{X}} + \sum_{\mathrm{Y}\neq\mathrm{T}} N_{n,\mathrm{T}\to\mathrm{Y}}}{\sum_{\mathrm{X}} N_{n,\mathrm{A}\to\mathrm{X}} + \sum_{\mathrm{Y}} N_{n,\mathrm{T}\to\mathrm{Y}}}, \tag{29}$$

where $N_{n,\mathrm{X}\to\mathrm{Y}}$ are the counts of X(ref) $\to$ Y(read) from the mismatch matrix.

Furthermore, suppose a modern contamination rate $r$ is provided by some external software, such as Schmutzi (Renaud *et al.*, 2015), ContaMix (Fu *et al.*, 2013) or ANGSD (Korneliussen *et al.*, 2014), we can calculate the observed G $\to$ A and C $\to$ T rates at position $n$ under both models incorporated with sequencing errors and modern contamination as follows,

$$\delta_{\mathrm{eff},n,\mathrm{G}\to\mathrm{A},r} = (1-r)\left(1-\frac{4}{3}\epsilon_n\right)\delta_{\mathrm{eff},n,\mathrm{G}\to\mathrm{A}} + \frac{\epsilon_n}{3}, \tag{30}$$

$$\delta_{\mathrm{eff},n,\mathrm{G}\to\mathrm{G},r} = -(1-r)\left(1-\frac{4}{3}\epsilon_n\right)\delta_{\mathrm{eff},n,\mathrm{G}\to\mathrm{A}} + 1 - \epsilon_n, \tag{31}$$

$$\delta_{\mathrm{eff},n,\mathrm{C}\to\mathrm{T},r} = (1-r)\left(1-\frac{4}{3}\epsilon_n\right)\delta_{\mathrm{eff},n,\mathrm{C}\to\mathrm{T}} + \frac{\epsilon_n}{3}, \tag{32}$$

$$\delta_{\mathrm{eff},n,\mathrm{C}\to\mathrm{C},r} = -(1-r)\left(1-\frac{4}{3}\epsilon_n\right)\delta_{\mathrm{eff},n,\mathrm{C}\to\mathrm{T}} + 1 - \epsilon_n, \tag{33}$$

$$\tilde{\delta}_{\mathrm{eff},n,\mathrm{G}\to\mathrm{A},r} = (1-r)\left(1-\frac{4}{3}\epsilon_n\right)\tilde{\delta}_{\mathrm{eff},n,\mathrm{G}\to\mathrm{A}} + \frac{\epsilon_n}{3}, \tag{34}$$

$$\tilde{\delta}_{\mathrm{eff},n,\mathrm{G}\to\mathrm{G},r} = -(1-r)\left(1-\frac{4}{3}\epsilon_n\right)\tilde{\delta}_{\mathrm{eff},n,\mathrm{G}\to\mathrm{A}} + 1 - \epsilon_n, \tag{35}$$

$$\tilde{\delta}_{\mathrm{eff},n,\mathrm{C}\to\mathrm{T},r} = (1-r)\left(1-\frac{4}{3}\epsilon_n\right)\tilde{\delta}_{\mathrm{eff},n,\mathrm{C}\to\mathrm{T}} + \frac{\epsilon_n}{3}, \tag{36}$$

$$\tilde{\delta}_{\mathrm{eff},n,\mathrm{C}\to\mathrm{C},r} = -(1-r)\left(1-\frac{4}{3}\epsilon_n\right)\tilde{\delta}_{\mathrm{eff},n,\mathrm{C}\to\mathrm{T}} + 1 - \epsilon_n. \tag{37}$$

To distinguish different notations in the above formulae, we refer to $\delta_{\mathrm{eff},n,\mathrm{X}\to\mathrm{Y},r}$ or $\tilde{\delta}_{\mathrm{eff},n,\mathrm{X}\to\mathrm{Y},r}$ as the theoretical X $\to$ Y change frequency with errors and contamination, while $\delta_{\mathrm{eff},n,\mathrm{X}\to\mathrm{Y}}$ or $\tilde{\delta}_{\mathrm{eff},n,\mathrm{X}\to\mathrm{Y}}$ is the version without errors and contamination from **Subsection** 3.1.1, 3.1.2 or 3.1.3. But they will all be presented later as $\delta_{\mathrm{eff},n,\mathrm{X}\to\mathrm{Y}}$ or $\tilde{\delta}_{\mathrm{eff},n,\mathrm{X}\to\mathrm{Y}}$ for short.

#### 3.2 Likelihood

In this subsection, we give the formulas of the multinomial regression likelihoods of both models as follows,

$$\begin{aligned} l = \sum_n \big[ & N_{n,\mathrm{G}\to\mathrm{A}} \log \delta_{\mathrm{eff},n,\mathrm{G}\to\mathrm{A}} + N_{n,\mathrm{G}\to\mathrm{G}} \log \delta_{\mathrm{eff},n,\mathrm{G}\to\mathrm{G}} \\ & + N_{n,\mathrm{C}\to\mathrm{T}} \log \delta_{\mathrm{eff},n,\mathrm{C}\to\mathrm{T}} + N_{n,\mathrm{C}\to\mathrm{C}} \log \delta_{\mathrm{eff},n,\mathrm{C}\to\mathrm{C}} \big], \end{aligned} \tag{38}$$

$$\begin{aligned} \tilde{l} = \sum_n \Big[ & N_{n,\mathrm{G}\to\mathrm{A}} \log \tilde{\delta}_{\mathrm{eff},n,\mathrm{G}\to\mathrm{A}} + N_{n,\mathrm{G}\to\mathrm{G}} \log \tilde{\delta}_{\mathrm{eff},n,\mathrm{G}\to\mathrm{G}} \\ & + N_{n,\mathrm{C}\to\mathrm{T}} \log \tilde{\delta}_{\mathrm{eff},n,\mathrm{C}\to\mathrm{T}} + N_{n,\mathrm{C}\to\mathrm{C}} \log \tilde{\delta}_{\mathrm{eff},n,\mathrm{C}\to\mathrm{C}} \Big]. \end{aligned} \tag{39}$$

where the form of $\delta_{\mathrm{eff},n,\mathrm{G}\to\mathrm{A}}$, $\delta_{\mathrm{eff},n,\mathrm{C}\to\mathrm{T}}$, $\tilde{\delta}_{\mathrm{eff},n,\mathrm{G}\to\mathrm{A}}$, and $\tilde{\delta}_{\mathrm{eff},n,\mathrm{C}\to\mathrm{T}}$ can be adopted according to different models described in the **Subsections** of 3.1.1, 3.1.2, 3.1.3 and 3.1.4. As was addressed previously, any parameter with a tilde symbol is related to the non-biotin model, otherwise the parameter is relevant to the biotin model. $N_{n,\mathrm{G}\to\mathrm{A}}$ and $N_{n,\mathrm{C}\to\mathrm{T}}$ are the counts of G(ref) $\to$ A(read) and C(ref) $\to$ T(read) reads at position $n$ across all the fragments, while $N_{n,\mathrm{G}\to\mathrm{G}}$ and $N_{n,\mathrm{C}\to\mathrm{C}}$ are the counts of the reads (G or C) staying the same as the reference (regarded as truth) at position $n$.

$$\left(\hat{\lambda}, \hat{\delta}_d, \hat{\delta}_s, \hat{\nu}\right) = \underset{\lambda,\delta_d,\delta_s,\nu}{\arg\max}\, l, \tag{40}$$

$$\left(\hat{\tilde{\lambda}}, \hat{\tilde{\delta}}_d, \hat{\tilde{\lambda}}_s, \hat{\tilde{\nu}}\right) = \underset{\tilde{\lambda},\tilde{\delta}_d,\tilde{\delta}_s,\tilde{\nu}}{\arg\max}\, \tilde{l}, \tag{41}$$

The approximated covariance matrices of both models, namely $V$ and $\tilde{V}$, can be given by the following Fisher's information matrices,

$$V = -\left.\begin{pmatrix} \frac{\partial^2 l}{\partial\lambda^2} & \frac{\partial^2 l}{\partial\lambda\partial\delta_d} & \frac{\partial^2 l}{\partial\lambda\partial\delta_s} & \frac{\partial^2 l}{\partial\lambda\partial\nu} \\ \frac{\partial^2 l}{\partial\lambda\partial\delta_d} & \frac{\partial^2 l}{\partial\delta^2} & \frac{\partial^2 l}{\partial\delta_d\partial\delta_s} & \frac{\partial^2 l}{\partial\delta_d\partial\nu} \\ \frac{\partial^2 l}{\partial\lambda\partial\delta_s} & \frac{\partial^2 l}{\partial\delta_d\partial\delta_s} & \frac{\partial^2 l}{\partial\delta_s^2} & \frac{\partial^2 l}{\partial\delta_s\partial\nu} \\ \frac{\partial^2 l}{\partial\lambda\partial\nu} & \frac{\partial^2 l}{\partial\delta_d\partial\nu} & \frac{\partial^2 l}{\partial\delta_s\partial\nu} & \frac{\partial^2 l}{\partial\nu^2} \end{pmatrix}^{-1}\right|_{(\lambda,\delta_d,\delta_s,\nu)=\left(\hat{\lambda},\hat{\delta}_d,\hat{\delta}_s,\hat{\nu}\right)}, \tag{42}$$

$$\tilde{V} = -\left.\begin{pmatrix} \frac{\partial^2 \tilde{l}}{\partial\tilde{\lambda}^2} & \frac{\partial^2 \tilde{l}}{\partial\tilde{\lambda}\partial\tilde{\delta}_d} & \frac{\partial^2 \tilde{l}}{\partial\tilde{\lambda}\partial\tilde{\delta}_s} & \frac{\partial^2 \tilde{l}}{\partial\tilde{\lambda}\partial\tilde{\nu}} \\ \frac{\partial^2 \tilde{l}}{\partial\tilde{\lambda}\partial\tilde{\delta}_d} & \frac{\partial^2 \tilde{l}}{\partial\tilde{\delta}_d{}^2} & \frac{\partial^2 \tilde{l}}{\partial\tilde{\delta}_d\partial\tilde{\delta}_s} & \frac{\partial^2 \tilde{l}}{\partial\tilde{\delta}_d\partial\tilde{\nu}} \\ \frac{\partial^2 \tilde{l}}{\partial\tilde{\lambda}\partial\tilde{\delta}_s} & \frac{\partial^2 \tilde{l}}{\partial\tilde{\delta}_d\partial\tilde{\delta}_s} & \frac{\partial^2 \tilde{l}}{\partial\tilde{\delta}_s{}^2} & \frac{\partial^2 \tilde{l}}{\partial\tilde{\delta}_s\partial\tilde{\nu}} \\ \frac{\partial^2 \tilde{l}}{\partial\tilde{\lambda}\partial\tilde{\nu}} & \frac{\partial^2 \tilde{l}}{\partial\tilde{\delta}_d\partial\tilde{\nu}} & \frac{\partial^2 \tilde{l}}{\partial\tilde{\delta}_s\partial\tilde{\nu}} & \frac{\partial^2 \tilde{l}}{\partial\tilde{\nu}^2} \end{pmatrix}^{-1}\right|_{\left(\tilde{\lambda},\tilde{\delta}_d,\tilde{\delta}_s,\tilde{\nu}\right)=\left(\hat{\tilde{\lambda}},\hat{\tilde{\delta}}_d,\hat{\tilde{\delta}}_s,\hat{\tilde{\nu}}\right)}. \tag{43}$$

$V$ and $\tilde{V}$ are for obtaining the asymptotic confidence intervals for the optimal estimates of $\left(\hat{\lambda}, \hat{\delta}_d, \hat{\delta}_s, \hat{\nu}\right)$ and $\left(\hat{\tilde{\lambda}}, \hat{\tilde{\delta}}_d, \hat{\tilde{\delta}}_s, \hat{\tilde{\nu}}\right)$.

### 4 Posterior Probability of Being Ancient

For a modern-contaminated sample whose overall contamination rate $r$ is given by some external software (Schmutzi (Renaud *et al.*, 2015) or ContaMix (Fu *et al.*, 2013)), another functionality of the presented tool is calculating the posterior probability of being ancient for each given ancient strand based on the mismatch observations of the first and last few positions and the strand's length.

#### 4.1 Likelihood based on observed mismatches

Once the four damaging parameters are estimated, the likelihood of a specific strand being ancient or modern based on the observed mismatches is defined in this subsection.

If the focal strand $k$ is modern, all the observed mismatches, denoted as $\vec{o^k}$, are assumed to be caused solely by sequencing errors $\vec{\epsilon^k}$, extracted from the base calling scores of the strand $k$. Given strand $k$ is a modern contamination, the likelihood of $\vec{o^k}$ for both biotin (the plain version) and non-biotin (the tilde version) are the same and as follows,

$$\mathrm{Lik}^m_{\vec{o^k}}\left(\vec{\epsilon^k}\right) = \widetilde{\mathrm{Lik}}^m_{\vec{o^k}}\left(\vec{\epsilon^k}\right) = \prod_n \left[\left(\frac{\epsilon^k_n}{3}\right)^{\sum_{\mathrm{X}\neq\mathrm{Y}} \chi^k_n(\mathrm{X}\to\mathrm{Y})} \left(1-\epsilon^k_n\right)^{\sum_{\mathrm{X}} \chi^k_n(\mathrm{X}\to\mathrm{X})}\right], \tag{44}$$

where the position index $n$ only takes the value corresponding to the first and last 15 cyclic position of the focal strand $k$.

Given the focal strand is from ancient endogenous materials, the observed mismatch can be due to sequencing errors as well as the damages caused by deaminations, which are featured by the inferred $(\hat{\lambda}, \hat{\lambda}_d, \hat{\delta}_s, \hat{\nu})$.

If we further assume the library preparation protocol satisfies the biotin model, the likelihood of $\vec{o^k}$ can be writtern as below,

$$
\begin{aligned}
&\mathrm{Lik}^a_{\vec{o^k}}\left(\lambda, \delta_d, \delta_s, \nu, \vec{\epsilon^k}\right) \\
=&\mathbf{E}\left\{\prod_n\left[\left(\frac{\epsilon^k_n}{3}+\delta_s-\frac{4\epsilon^k_n\delta_s}{3}\right)^{\chi^k_{n,1}(\mathrm{C}\to\mathrm{T})}\left(\frac{\epsilon^k_n}{3}\right)^{\sum_{\mathrm{X}\neq\mathrm{C,T}}\chi^k_{n,1}(\mathrm{C}\to\mathrm{X})}\left(1-\epsilon^k_n-\delta_s+\frac{4\epsilon^k_n\delta_s}{3}\right)^{\chi^k_{n,1}(\mathrm{C}\to\mathrm{C})}\right.\right. \\
&\times\left(\frac{\epsilon^k_n}{3}+\delta_s-\frac{4\epsilon^k_n\delta_s}{3}\right)^{\chi^k_{n,2}(\mathrm{G}\to\mathrm{A})}\left(\frac{\epsilon^k_n}{3}\right)^{\sum_{\mathrm{X}\neq\mathrm{G,A}}\chi^k_{n,2}(\mathrm{G}\to\mathrm{X})}\left(1-\epsilon^k_n-\delta_s+\frac{4\epsilon^k_n\delta_s}{3}\right)^{\chi^k_{n,2}(\mathrm{G}\to\mathrm{G})} \\
&\times\left(\frac{\epsilon^k_n}{3}+\delta_d-\frac{4\epsilon^k_n\delta_d}{3}\right)^{\chi^k_{n,3}(\mathrm{C}\to\mathrm{T})}\left(\frac{\epsilon^k_n}{3}\right)^{\sum_{\mathrm{X}\neq\mathrm{C,T}}\chi^k_{n,3}(\mathrm{C}\to\mathrm{X})}\left(1-\epsilon^k_n-\delta_d+\frac{4\epsilon^k_n\delta_d}{3}\right)^{\chi^k_{n,3}(\mathrm{C}\to\mathrm{C})} \\
&\times\left(\frac{\epsilon^k_n}{3}+\delta_d-\frac{4\epsilon^k_n\delta_d}{3}\right)^{\chi^k_{n,4}(\mathrm{G}\to\mathrm{A})}\left(\frac{\epsilon^k_n}{3}\right)^{\sum_{\mathrm{X}\neq\mathrm{G,A}}\chi^k_{n,4}(\mathrm{G}\to\mathrm{X})}\left(1-\epsilon^k_n-\delta_d+\frac{4\epsilon^k_n\delta_d}{3}\right)^{\chi^k_{n,4}(\mathrm{G}\to\mathrm{G})} \\
&\times\left(1-\epsilon^k_n\right)^{\sum_{\mathrm{X=A,T}}\chi^k_n(\mathrm{X}\to\mathrm{X})+\chi^k_{n,1}(\mathrm{G}\to\mathrm{G})+\chi^k_{n,3}(\mathrm{G}\to\mathrm{G})+\chi^k_{n,2}(\mathrm{C}\to\mathrm{C})+\chi^k_{n,4}(\mathrm{C}\to\mathrm{C})} \\
&\left.\left.\times\left(\frac{\epsilon^k_n}{3}\right)^{\sum_{\substack{\mathrm{X}\neq\mathrm{Y}\\ \mathrm{X=A,T}}}\chi^k_n(\mathrm{X}\to\mathrm{Y})+\sum_{\mathrm{Y}\neq\mathrm{G}}\left[\chi^k_{n,1}(\mathrm{G}\to\mathrm{Y})+\chi^k_{n,3}(\mathrm{G}\to\mathrm{Y})\right]+\sum_{\mathrm{Y}\neq\mathrm{C}}\left[\chi^k_{n,2}(\mathrm{C}\to\mathrm{Y})+\chi^k_{n,4}(\mathrm{C}\to\mathrm{Y})\right]}\right]\right\}, \qquad (45)
\end{aligned}
$$

where $\mathbf{E}\{\cdot\}$ is the expectation in the sense of both overhangs and nick placement given $(\hat{\lambda}, \hat{\lambda}_d, \hat{\delta}_s, \hat{\nu})$. $\chi^k_n(\cdot)$ and $\chi^k_{n,i}(\cdot)$ $(i = 1, 2, 3, 4)$ are all indicator functions. $\chi^k_n(\cdot) = 1$ if and only if the specified nucleotide change is observed at position $n$ and $n$ is within the first 15 and last 15 cyclic positions of a strand, $\chi^k_n(\cdot) = 0$ otherwise; $\chi^k_{n,i}(\cdot) = 1$ if and only if the specified nucleotide change is observed at position $n$, $n$ is within the first 15 and last 15 cyclic positions and $n$ is also in the region indicated by $i$, $\chi^k_{n,i}(\cdot) = 0$ otherwise. $i$ = 1, 2, 3, 4 represent that position $n$ is within the left 5' overhang, within the right 5' overhang, within the double-stranded region but before the possible nick, and within the double-stranded region but after the possible nick, respectively. $\frac{\epsilon}{3} + \delta - \frac{4\epsilon\delta}{3} = \frac{\epsilon}{3}(1-\delta) + (1-\epsilon)\delta$ and $1 - \epsilon - \delta + \frac{4\epsilon\delta}{3} = \frac{\epsilon}{3}\delta + (1-\epsilon)(1-\delta)$ represent the multiplicative effects of pure deamination and sequencing errors.

If we assume the library preparation protocol is based on the non-biotin model, given the focal strand $k$ is from ancient materials, the likelihood of the observed mismatches $\vec{o^k}$, $\widetilde{\mathrm{Lik}}^a_{\vec{o^k}}$, can be different from $\mathrm{Lik}^a_{\vec{o^k}}$ in **Equation** 45, since in this case, the focal strand has 50% of the chances to be reverse-complement of an original strand,

$$
\begin{aligned}
&\widetilde{\mathrm{Lik}}^{a}_{\vec{o^k}}\left(\lambda, \delta_d, \delta_s, \nu, \vec{\epsilon^k}\right) = 0.5\mathrm{Lik}^{a}_{\vec{o^k}}\left(\lambda, \delta_d, \delta_s, \nu, \vec{\epsilon^k}\right) + 0.5\mathbf{E}\left\{\prod_n \Bigg[\right. \\
&\left(\frac{\epsilon^k_{L-n+1}}{3} + \delta_s - \frac{4\epsilon^k_{L-n+1}\delta_s}{3}\right)^{\tilde{\chi}^k_{n,1}(\mathrm{C}\to\mathrm{T})} \left(\frac{\epsilon^k_{L-n+1}}{3}\right)^{\sum_{\mathrm{X}\neq\mathrm{C,T}} \tilde{\chi}^k_{n,1}(\mathrm{C}\to\mathrm{X})} \left(1 - \epsilon^k_{L-n+1} - \delta_s + \frac{4\epsilon^k_{L-n+1}\delta_s}{3}\right)^{\tilde{\chi}^k_{n,1}(\mathrm{C}\to\mathrm{C})} \\
&\times \left(\frac{\epsilon^k_{L-n+1}}{3} + \delta_s - \frac{4\epsilon^k_{L-n+1}\delta_s}{3}\right)^{\tilde{\chi}^k_{n,2}(\mathrm{G}\to\mathrm{A})} \left(\frac{\epsilon^k_{L-n+1}}{3}\right)^{\sum_{\mathrm{X}\neq\mathrm{G,A}} \tilde{\chi}^k_{n,2}(\mathrm{G}\to\mathrm{X})} \left(1 - \epsilon^k_{L-n+1} - \delta_s + \frac{4\epsilon^k_{L-n+1}\delta_s}{3}\right)^{\tilde{\chi}^k_{n,2}(\mathrm{G}\to\mathrm{G})} \\
&\times \left(\frac{\epsilon^k_{L-n+1}}{3} + \delta_d - \frac{4\epsilon^k_{L-n+1}\delta_d}{3}\right)^{\tilde{\chi}^k_{n,3}(\mathrm{C}\to\mathrm{T})} \left(\frac{\epsilon^k_{L-n+1}}{3}\right)^{\sum_{\mathrm{X}\neq\mathrm{C,T}} \tilde{\chi}^k_{n,3}(\mathrm{C}\to\mathrm{X})} \left(1 - \epsilon^k_{L-n+1} - \delta_d + \frac{4\epsilon^k_{L-n+1}\delta_d}{3}\right)^{\tilde{\chi}^k_{n,3}(\mathrm{C}\to\mathrm{C})} \\
&\times \left(\frac{\epsilon^k_{L-n+1}}{3} + \delta_d - \frac{4\epsilon^k_{L-n+1}\delta_d}{3}\right)^{\tilde{\chi}^k_{n,4}(\mathrm{G}\to\mathrm{A})} \left(\frac{\epsilon^k_{L-n+1}}{3}\right)^{\sum_{\mathrm{X}\neq\mathrm{G,A}} \tilde{\chi}^k_{n,4}(\mathrm{G}\to\mathrm{X})} \left(1 - \epsilon^k_{L-n+1} - \delta_d + \frac{4\epsilon^k_{L-n+1}\delta_d}{3}\right)^{\tilde{\chi}^k_{n,4}(\mathrm{G}\to\mathrm{G})} \\
&\times \left(1 - \epsilon^k_{L-n+1}\right)^{\sum_{\mathrm{X=A,T}} \tilde{\chi}^k_n(\mathrm{X}\to\mathrm{X}) + \tilde{\chi}^k_{n,1}(\mathrm{G}\to\mathrm{G}) + \tilde{\chi}^k_{n,3}(\mathrm{G}\to\mathrm{G}) + \tilde{\chi}^k_{n,2}(\mathrm{C}\to\mathrm{C}) + \tilde{\chi}^k_{n,4}(\mathrm{C}\to\mathrm{C})} \\
&\times \left(\frac{\epsilon^k_{L-n+1}}{3}\right)^{\sum_{\substack{\mathrm{X}\neq\mathrm{Y} \\ \mathrm{X=A,T}}} \tilde{\chi}^k_n(\mathrm{X}\to\mathrm{Y}) + \sum_{\mathrm{Y}\neq\mathrm{G}}\left[\tilde{\chi}^k_{n,1}(\mathrm{G}\to\mathrm{Y}) + \tilde{\chi}^k_{n,3}(\mathrm{G}\to\mathrm{Y})\right] + \sum_{\mathrm{Y}\neq\mathrm{C}}\left[\tilde{\chi}^k_{n,2}(\mathrm{C}\to\mathrm{Y}) + \tilde{\chi}^k_{n,4}(\mathrm{C}\to\mathrm{Y})\right]} \left.\Bigg]\right\}, \qquad (46)
\end{aligned}
$$

where $\tilde{\chi}^k_n(\cdot)$ and $\tilde{\chi}^k_{n,i}(\cdot)$ $(i = 1, 2, 3, 4)$ are all indicator functions defined similarly as $\chi^k_n(\cdot)$ and $\chi^k_{n,i}(\cdot)$ $(i = 1, 2, 3, 4)$ respectively. The only difference is that the hat versions focus on the relationships of the nicks and overhangs on the reverse-complement of the focal strand. Hence their corresponding sequencing errors has the subscript $L - n + 1$, which reflects the position on the focal strand.

#### 4.2 Likelihood based on strand length

Apart from the information delivered by nucleotide mismatches, the length of the focal strand can also help to distinguish whether the strand is ancient or not. By assuming the length distributions of the ancient endogenous strands and the modern contamination strands are distinguishable, and can be approximated by the bounded discrete normal distributions $\mathcal{N}_b\left(\mu_a, \sigma_a; l_{\min}, l_{\max}\right)$ and $\mathcal{N}_b\left(\mu_m, \sigma_m; l_{\min}, l_{\max}\right)$, we can give the likelihoods based on the observation that strand $k$ is of the length $L_k$,

$$
\mathrm{Lik}^a_{L_k}\left(\mu_a, \sigma_a\right) \triangleq f_b\left(L_k; \mu_a, \sigma_a, l_{\min}, l_{\max}\right), \qquad (47)
$$

$$
\mathrm{Lik}^m_{L_k}\left(\mu_m, \sigma_m\right) \triangleq f_b\left(L_k; \mu_m, \sigma_m, l_{\min}, l_{\max}\right), \qquad (48)
$$

where $f_b(l; \mu, \sigma, l_{\min}, l_{\max})$ is the probability function of a bounded discrete normal distribution, and its format is defined as below:

$$
f_b(l; \mu, \sigma, l_{\min}, l_{\max}) = \begin{cases} 0 & l < l_{\min}, \\ \frac{\Phi(l+0.5;\mu,\sigma) - \Phi(l-0.5;\mu,\sigma)}{\Phi(l_{\max}+0.5;\mu,\sigma) - \Phi(l_{\min}-0.5;\mu,\sigma)} & l_{\min} \leq l \leq l_{\max}, \\ 0 & l > l_{\max}, \end{cases} \qquad (49)
$$

where $\Phi\left(x; \mu, \sigma\right)$ is the cumulative distribution function of the normal distribution $\mathcal{N}\left(\mu, \sigma\right)$.

#### 4.3 Posterior probability of being ancient

Once the four damaging parameters are estimated, $(\hat{\mu}_a, \hat{\sigma}_a, \hat{\mu}_m, \hat{\sigma}_m)$ can be further obtained by optimizing the following log-likelihood function,

$$\begin{aligned} & l(\mu_a, \sigma_a, \mu_m, \sigma_m) \\ = & \sum_k \log \left[ (1-r) \operatorname{Lik}^a_{L_k}(\mu_a, \sigma_a) \operatorname{Lik}^a_{\vec{o^k}} \left( \hat{\lambda}, \hat{\delta}_d, \hat{\delta}_s, \hat{\nu}, \hat{\vec{\epsilon^k}} \right) + r \operatorname{Lik}^m_{L_k}(\mu_m, \sigma_m) \operatorname{Lik}^m_{\vec{o^k}} \left( \hat{\vec{\epsilon^k}} \right) \right], \end{aligned} \tag{50}$$

where $(1-r) \operatorname{Lik}^a_{L_k}(\mu_a, \sigma_a) \operatorname{Lik}^a_{\vec{o^k}} \left( \hat{\lambda}, \hat{\delta}_d, \hat{\delta}_s, \hat{\nu}, \hat{\vec{\epsilon^k}} \right) + r \operatorname{Lik}^m_{L_k}(\mu_m, \sigma_m) \operatorname{Lik}^m_{\vec{o^k}} \left( \hat{\vec{\epsilon^k}} \right)$ represents the joint probability of observing read $k$ length and deamination patterns. More detailedly, $(1-r)$ is prior probability read $k$ is ancient, and $\operatorname{Lik}^a_{L_k}(\mu_a, \sigma_a) \operatorname{Lik}^a_{\vec{o^k}} \left( \hat{\lambda}, \hat{\delta}_d, \hat{\delta}_s, \hat{\nu}, \hat{\vec{\epsilon^k}} \right)$ is the joint probability of observing read $k$ length and deamination patterns given it is ancient. $r$ is prior probability read $k$ is a modern contamination, and $\operatorname{Lik}^m_{L_k}(\mu_m, \sigma_m) \operatorname{Lik}^m_{\vec{o^k}} \left( \hat{\vec{\epsilon^k}} \right)$ represents the joint probability of observing read $k$ length and deamination patterns given it is modern.

Note that to optimize the above log-likelihood function requires to go through all the provided reads, and will take relatively longer compared with the procedure of estimating the four damaging parameters, and ngsBriggs provides a multithreading optimization option for this.

The posterior probability of strand $k$ being ancient, $P^a_k$, can then be given as below,

$$P^a_k = \frac{(1-r) \operatorname{Lik}^a_{L_k}(\hat{\mu}_a, \hat{\sigma}_a) \operatorname{Lik}^a_{\vec{o^k}} \left( \hat{\lambda}, \hat{\delta}_d, \hat{\delta}_s, \hat{\nu}, \hat{\vec{\epsilon^k}} \right)}{(1-r) \operatorname{Lik}^a_{L_k}(\hat{\mu}_a, \hat{\sigma}_a) \operatorname{Lik}^a_{\vec{o^k}} \left( \hat{\lambda}, \hat{\delta}_d, \hat{\delta}_s, \hat{\nu}, \hat{\vec{\epsilon^k}} \right) + r \operatorname{Lik}^m_{L_k}(\hat{\mu}_m, \hat{\sigma}_m) \operatorname{Lik}^m_{\vec{o^k}} \left( \hat{\vec{\epsilon^k}} \right)} \tag{51}$$

#### 4.4 Recalibration of the nucleotide likelihoods

Based on the chosen model, the previously inferred $\left( \hat{\lambda}, \hat{\delta}_d, \hat{\delta}_s, \hat{\nu} \right)$ and $P^a_k$, we can then recalibrate the likelihood of the observed nucleotide at a given position of a specific strand. As given in **Equations** 31-37, if we replace $(1-r)$ and $\epsilon_n$ by their strand $k$ specific counterparts, i.e., $P^a_k$ and $\epsilon^k_n$,

$$\delta_{\text{eff},n,\text{G}\to\text{A},r} = P^a_k \left( 1 - \frac{4}{3}\epsilon^k_n \right) \delta_{\text{eff},n,\text{G}\to\text{A}} + \frac{\epsilon^k_n}{3},\ \delta_{\text{eff},n,\text{G}\to\text{G},r} = -P^a_k \left( 1 - \frac{4}{3}\epsilon^k_n \right) \delta_{\text{eff},n,\text{G}\to\text{A}} + 1 - \epsilon^k_n \tag{52}$$

$$\delta_{\text{eff},n,\text{C}\to\text{T},r} = P^a_k \left( 1 - \frac{4}{3}\epsilon^k_n \right) \delta_{\text{eff},n,\text{C}\to\text{T}} + \frac{\epsilon^k_n}{3},\ \delta_{\text{eff},n,\text{C}\to\text{C},r} = -P^a_k \left( 1 - \frac{4}{3}\epsilon^k_n \right) \delta_{\text{eff},n,\text{C}\to\text{T}} + 1 - \epsilon^k_n \tag{53}$$

$$\tilde{\delta}_{\text{eff},n,\text{G}\to\text{A},r} = P^a_k \left( 1 - \frac{4}{3}\epsilon^k_n \right) \tilde{\delta}_{\text{eff},n,\text{G}\to\text{A}} + \frac{\epsilon^k_n}{3},\ \tilde{\delta}_{\text{eff},n,\text{G}\to\text{G},r} = -P^a_k \left( 1 - \frac{4}{3}\epsilon^k_n \right) \tilde{\delta}_{\text{eff},n,\text{G}\to\text{A}} + 1 - \epsilon^k_n \tag{54}$$

$$\tilde{\delta}_{\text{eff},n,\text{C}\to\text{T},r} = P^a_k \left( 1 - \frac{4}{3}\epsilon^k_n \right) \tilde{\delta}_{\text{eff},n,\text{C}\to\text{T}} + \frac{\epsilon^k_n}{3},\ \tilde{\delta}_{\text{eff},n,\text{C}\to\text{C},r} = -P^a_k \left( 1 - \frac{4}{3}\epsilon^k_n \right) \tilde{\delta}_{\text{eff},n,\text{C}\to\text{T}} + 1 - \epsilon^k_n \tag{55}$$

If the change of X→Y is not listed above, then we do not need to modify the likelihood, namely,

$$\delta_{\text{eff},n,\text{X}\to\text{Y},r} = \tilde{\delta}_{\text{eff},n,\text{X}\to\text{Y},r} = \begin{cases} 1 - \epsilon^k_n, & \text{X} = \text{Y}, \\ \frac{\epsilon^k_n}{3}, & \text{X} \neq \text{Y}. \end{cases} \tag{56}$$

For a specific strand, its length is fixed, the forms of $\delta_{\text{eff},n,\text{X}\to\text{Y}}$ and $\tilde{\delta}_{\text{eff},n,\text{X}\to\text{Y}}$ can be chosen from **Subsection** 3.1.1 and 3.1.2.

If $P^a_k = 1$ and $\delta_{\text{eff},n,\text{G}\to\text{A}} = \delta_{\text{eff},n,\text{C}\to\text{T}} = \tilde{\delta}_{\text{eff},n,\text{G}\to\text{A}} = \tilde{\delta}_{\text{eff},n,\text{C}\to\text{T}} = 1 - \delta_{\text{eff},n,\text{G}\to\text{G}} = 1 - \delta_{\text{eff},n,\text{C}\to\text{C}} = 1 - \tilde{\delta}_{\text{eff},n,\text{G}\to\text{G}} = 1 - \tilde{\delta}_{\text{eff},n,\text{C}\to\text{C}} = 0$ in **Equations** 52-55, which corresponds to the case without sequencing errors and contamination, such equations will degenerate to the same form as **Equation** 56.

#### 5 Data processing commands

Throughout the main article and supplementary results described in **Section** 6 we have simulated, using the tool NGSNGS (Henriksen *et al.*, 2023), various datasets with different characteristics. The quality profiles an length distribution files used for particular simulations are provided by NGSNGS. While every analysis is meticulously conducted on numerous datasets, we have for simplicity only shown one simulation command for the datasets used for the respective analysis below, with the alterations between the commands being limited to the the seed value, certain parameters for both deaminations model, length distribution and number of reads to be simulated. All datasets were simulated from chromosome 22 from the human reference genome build GRCh37 (Church *et al.*, 2011), and stored directly in a sequence alignment format (Li *et al.*, 2009; CB *et al.*, 2022) avoiding any potential short fragments with an exacerbated PMD being unable to align.

##### 5.1 Simulated data for inference of Briggs parameters

For the analysis described in **Section** 6.1 we simulated five groups of datasets, with varying numbers of sequencing reads (*-r*) $10^3$, $10^4$, $10^5$, $10^6$ and $10^7$ respectively. Within each group, a 100 independent samples was simulated with the biotin deamination model (*-m b7*, results in **Section** 6.1.1), and another 100 samples with the non-biotin model (*-m b*, (results in **Section** 6.1.2), with only the seed value varying between each of the datasets. For both deamination models, the Briggs parameters utilized were chosen from the original briggs article (Briggs *et al.*, 2007), i.e. 0.024, 0.36, 0.68, 0.0097 ($\nu$, $\lambda$, $\delta_s$, $\delta_d$).

**Biotin simulated data with $10^3$ reads.**

```
ngsngs -i chr22.fa -r 1000 -t 1 -s 1 -lf Test_Examples/Size_dist_sampling.txt
   -seq SE -ne -q1 Test_Examples/AccFreqL150R1.txt -m b7
   ,0.024,0.36,0.68,0.0097 -f bam -o OutputBiotin
```

**Non-Biotin simulated data with $10^3$ reads.**

```
ngsngs -i chr22.fa -r 1000 -t 1 -s 21 -lf Test_Examples/Size_dist_sampling.txt
   -seq SE -ne -q1 Test_Examples/AccFreqL150R1.txt -m b
   ,0.024,0.36,0.68,0.0097 -f bam -o OutputNonBiotin
```

##### 5.2 Simulated data used by contamination scenario for inferring parameters

In our rigorous analysis of the influence of contamination, we opted to emulate various contamination levels, i.e. 10%, 20%, 30%, 40% and 50% respectively. For each of these groups of contamination rates, we chose to simulate the ancient endogenous and modern exogenous content independently, to more accurately represent their diverse sequence characteristics. Once the simulations had been completed, both components were merged to obtain 20 samples within each group with a total of $10^6$ reads, such that contamination rate of 10% constitutes $9 \cdot 10^5$ ancient reads with $10^5$ modern reads, and similarly a sample with a contamination rate of 50 % were comprised of $5 \cdot 10^5$ ancient- and modern reads.

**Modern-day sequencing data**

Five datasets representing modern contaminants, with $10^5$, $2{\cdot}10^5$, $3{\cdot}10^5$, $4{\cdot}10^5$ and $5{\cdot}10^5$ reads were simulated and under the assumption of modern sequences sharing similar characteristics, repeatedly employed when merging with their ancient counterpart to obtain the desired contamination scenarios.

Modern sequencing platforms are capable of producing sequencing reads at 150 bp or longer, making them easily distinguishable from ancient reads which predominantly are shorter than 100 bp and mostly situated around 60 bp. To assess our ngsBriggs remained accurate in the different contamination scenarios, we deliberately sampled the modern reads from a $\mathcal{N}(100, 5)$ distribution. Thus generating a potential overlap in sequence length across the two components, effectively making it more problematic to solely differentiate the sequence reads based on the length.

```
ngsngs -i chr22.fa -r 100000 -s 100 -t 1 -ld Norm,100,5 -seq SE -q1
    Test_Examples/AccFreqL150R1.txt -f bam -o HumanContamination100k
```

**Ancient reads**

The ancient datasets prior to merging with the modern simulated, were generated in accordance with the command described in **Section** 5.1 with the differences as previously mentioned being deamination parameters, read counts, and seed values. For each of the intended contamination rates $9 \cdot 10^5$ (10% contamination), $8 \cdot 10^5$ (20%), $7 \cdot 10^5$ (30%), $6 \cdot 10^5$ (40%) and $5 \cdot 10^5$ (50%) we simulated three deamination scenarios with the biotin- and three with the non-biotin model. Each scenario contained 20 samples, and included the default Briggs parameters 0.024, 0.36, 0.68, 0.0097 ($\nu$, $\lambda$, $\delta_s$, $\delta_d$; **Section** 6.5.1), and increased For each of the desired number of read groups $9 \cdot 10^5$ (10% contamination), $8 \cdot 10^5$ (20%), $7 \cdot 10^5$ (30%), $6 \cdot 10^5$ (40%) and $5 \cdot 10^5$ (50%) we simulated 20 samples. For each contamination rate we included different deamination models $\lambda$ (0.7; **Section** 6.5.2) and decreased $\lambda$ (0.1; **Section** 6.5.3). With only one example of an increased $\lambda$ parameter with the non-biotin model being provided:

```
ngsngs -i chr22.fa -r 900000 -t 1 -s 220 -lf Test_Examples/Size_dist_sampling.
    txt -seq SE -q1 Test_Examples/AccFreqL150R1.txt -m b,0.024,0.7,0.68,0.0097
    -f bam -o NonBiotin09contamination
```

#### 5.3 Simulated contamination data when estimating read level probability of ancientness

To discriminate the ancient- from modern reads, we used a similar approach to the one described in **Section** 5.2 and simulated five ancient datasets ($9 \cdot 10^5$ reads) for both the biotin and non-biotin model, before reusing one of the modern-day sample ($10^5$ reads) as the exogenous component, yielding merged samples with a total number of $10^6$ reads and fixed contamination rate of 10 %.

The five samples simulated with the biotin model and the five samples simulated with the non-biotin model had a distinct deamination pattern, using default Briggs parameters 0.024, 0.36, 0.68, 0.0097 ($\nu$, $\lambda$, $\delta_s$, $\delta_d$), increased and decreased $\lambda$ (0.7 and 0.1), increased and decreased $\delta_s$ (0.9 and 0.3; **Section** 6.5.5). The example below describes the scenario with an increased $\delta_s$ rate for the biotin model.

```
ngsngs -i chr22.fa -r 900000 -s 2 -t 1 -lf LogNorm405_30125.txt -seq SE -q1
    Test_Examples/AccFreqL150R1.txt -m b7,0.024,0.36,0.9,0.0097 -f bam -o
    Anc900kDeltaS09
```

The ancient fragment length distribution file used was generated from cumulative distribution functions of a constrained lognormal distribution(4,0.5) with a lower- and upper limit of 30 and 125. Similarly the modern distribution $\mathcal{N}$(130,5) was limited to an upper length of 145.

#### 5.4 Creation of mismatch matrix

All three tools ngsBriggs, mapDamage 2.0, and PMDtools all rely upon a common data structure, before conducting their subsequent computational processes, the mismatch matrix (or mismatch matrix, shown in **Table** 2). Before initiating any computation procedures, however, mapDamage 2.0 and ngsBriggs also require each aligned sequenced read to have an MD tag present.

Tool-specific requirements involve providing the reference genome, wherein mapDamage 2.0 generates a mismatch matrix based on this input. As visualized in **Figure** 1, ngsBriggs can despite lacking a reference genome, still perform its computational procedures by generating the mismatch matrix solely based on the MD tag. The MD tag encodes information mismatching bases at a given position of the read (Li *et al.*, 2009; CB *et al.*, 2022). By combining the information stored in the MD tag and the equally important CIGAR string, the reference sequence can be reconstructed and thereby create the mismatch matrix. However several CIGAR string operations, including soft-clipping ('S'), which can be caused by various artifacts, are not represented in the MD tag, and thus these bases won't be considered during reconstruction. Consequently using this alternative approach (right side of part 1 in the **blue** box of **Figure** 1), slight variations to the mapDamage 2.0, PMDtools and ngsBriggs reference-based construction (left side of part 1 in the **blue** box of **Figure** 1) is expected. However, since the soft-clips in the 5' or 3' of the fragment remain unused the C→T frequency shouldn't be influenced.

With one example of the mismatch matrix construction for one individual from the VK population (Margaryan *et al.*, 2020) using mapDamage 2.0 and PMDtools.

```
mapDamage -i VK99.bam -r hg19.fa -d results_VK99
```

```
samtools view VK99.bam | python2 pmdtools.0.60.py --deamination
```

Following the creation of the mismatch matrix, all tools can proceed with additional computational steps. We chose to measure the wall-clock running time for the different steps across all three tools such that we could make comparable time measurements, with the mean wall-clock time shown in the first three lines across the populations in the main article **Table** 1, with the equivalent median wall-clock running times shown in **Table** 4.

#### 5.5 Inference of Briggs parameters

From the mismatch matrix, mapDamage 2.0 can complete the statistical estimations as follows.

```
mapDamage -d results_VK99 --stats-only
```

With the equivalent ngsBriggs command, generating both the mismatch matrix and inferring the parameters using the reference-based approach (*-ref*) assuming a biotin model (*-model b*).

```
./metadamage briggs -bam VK99.bam -model b -ref hg19.fa
```

When inferring these parameters, contamination rates can be considered by supplying *-eps* and the non-biotin model can be selected with *-model nb*, with an example of simulated data with 10 % contamination below.

```
./metadamage briggs -bam NbCont01RegParam.bam -model nb -eps 0.1 -ref chr22.fa
```

#### 5.6 Discriminating ancient from modern reads

When decontaminating samples, discriminating ancient- from modern reads (as visualized in the **red** section of **Figure** 1), the user is required to provide a *.bed* file, specifying chromosomal coordinates. The coordinates serve to specify the primary region from which the reads are sampled, prior to calculating the read specific posterior probability. This feature enables the users to decontaminate not only the entire genome, but also specific regions of interest. These regions could be crucial when performing downstream analysis of specific genes or conserved regions across multiple species. When supplying a *.bed* file, whether for the full genome, or specific regions, the chromosomal coordinates must be in accordance with the reference genomes index file (*.fai*).

```
./metadamage briggs -bam NbCont01RegParam.bam -model b -eps 0.1 -ibam
    NbCont01RegParam.bam -obam NbCont01RegParam_scores.bam -isrecal 1 -ibed
    chr22.bed -chr chr22 -nthread 1
```

The input bed file in this instance (*-ibed*) contains the following information:

```
chr22   1   17244502
```

Similarly, PMDtools can assign each read with a log-likelihood ratio, to discriminate ancient- from modern reads, with an example below.

```
samtools view -h NbCont01RegParam.bam | python2 pmdtools.0.60.py -r chr22.fa
    --header -p
```

#### 5.7 Inferring parameters and decontaminating with mismatch matrix and strand length files

To make consistent and equal wall-clock time evaluations, we adhered to the previously described commands. While we have presented two alternative approaches in ngsBriggs initial stage when inferring the briggs parameters, all three tools are executed with a provided reference genome. Such that they all would theoretically rely on the same input bottleneck. We expect the wall-clock time for ngsBriggs would diminish if solely providing the *.bam* file relying on the MD tag. The fastest running time of ngsBriggs is expected to be observed by solely providing the the mismatch matrix and strand length input files as visualized in the (**green** box) in **Figure** 1, since it removes the need for utilizing each sequence read information which would be the most computationally demanding part of inferring the parameters. With examples of the commands shown below.

The equivalent commands as presented in the previous sections, when solely inferring parameters, without and with provided contamination ratio would be:

```
./metadamage briggs -tab MisMatch.txt -model nb -len StrandLength.txt
./metadamage briggs -tab MisMatch.txt -model nb -len StrandLength.txt -eps 0.1
```

Assigning each read with a probability would still require the alignment file to generate the additional probability tag for each read within the file.

```
./metadamage briggs -tab MisMatch.txt -model b -len StrandLength.txt -eps 0.1
    -ibam NbCont01RegParam.bam -obam NbCont01RegParam_scores.bam -isrecal 1 -
    ibed chr22.bed -nthread 1
```

The MisMatch.txt file has the structure as shown in **Table** 2 and the StrandLength.txt is simply a two-column tab-separated file with the read length and count.

### 6 Supplementary Results

The results presented in the main article predominantly encapsulate benchmarking metrics pertaining to the simulated data. Whereas the subsequent sections delve into a more comprehensive comparative analysis of the performance of mapDamage 2.0 and ngsBriggs both in the context of simulated- and empirical datasets.

#### 6.1 Inferring Briggs parameters on simulated data across deamination models

Another prominent feature of aDNA is the inherent limited number of sequencing reads. To assess the robustness of ngsBriggs inference method, under these conditions with low endogenous content, we used datasets with $10^3$, $10^4$, $10^5$, $10^6$, $10^7$ reads respectively (**Section** 5.1)

##### 6.1.1 Biotin Model

For the inferred $\lambda$ (**Figure** 7), $\delta_d$ (**Figure** 8), and $\delta_s$ (**Figure** 9) parameters, an overarching trend becomes evident. With an increased number of reads, both mapDamage 2.0 and ngsBriggs biotin model exhibit a reduced variance, simultaneously with a more pronounced disparity between the estimated parameters across the two tools.

**Inferring $\lambda$ parameter**

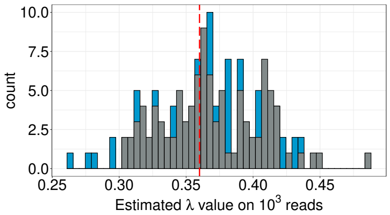

**(a)** $10^3$ reads, ngsBriggs biotin (RMSE = 0.038096) and mapDamage 2.0 (RMSE = 0.037766).

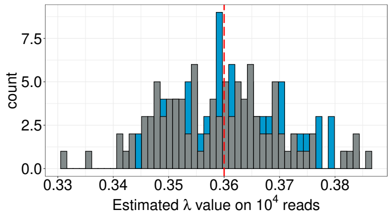

**(b)** $10^4$ reads, ngsBriggs biotin (RMSE = 0.009448) and mapDamage 2.0 (RMSE = 0.010904).

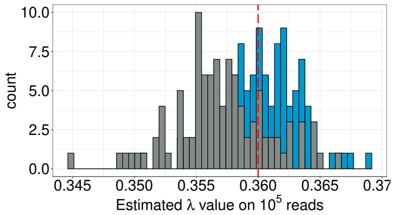

**(c)** $10^5$ reads, ngsBriggs biotin (RMSE = 0.002988) and mapDamage 2.0 (RMSE = 0.004787).

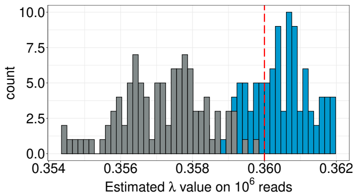

**(d)** $10^6$ reads, ngsBriggs biotin (RMSE = 0.000934) and mapDamage 2.0 (RMSE = 0.003078).

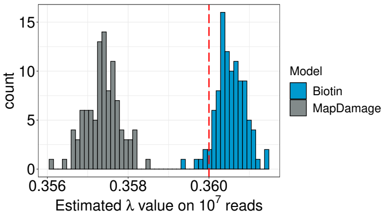

**(e)** $10^7$ reads, ngsBriggs biotin (RMSE = 0.000641) and mapDamage 2.0 (RMSE = 0.002659).

**Figure 7:** Inferred $\lambda$ values using ngsBriggs Biotin and mapDamage 2.0 model across datasets with a different number of reads.

##### Inferring $\delta_d$ parameter

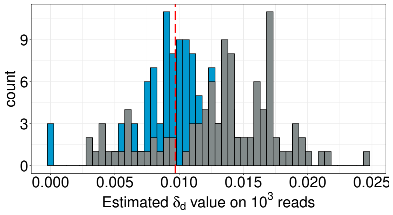

**(a)** $10^3$ reads, ngsBriggs biotin (RMSE = 0.002875) and mapDamage 2.0 (RMSE = 0.005758)

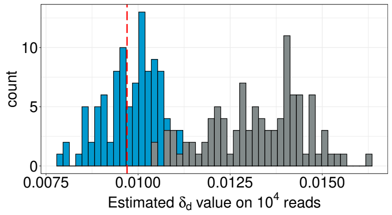

**(b)** $10^4$ reads, ngsBriggs biotin (RMSE = 0.000778) and mapDamage 2.0 (RMSE = 0.003762)

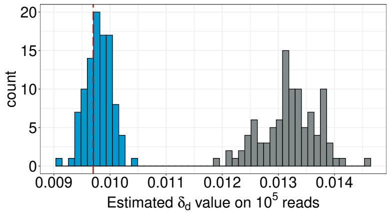

**(c)** $10^5$ reads, ngsBriggs biotin (RMSE = 0.000250) and mapDamage 2.0 (RMSE = 0.003536)

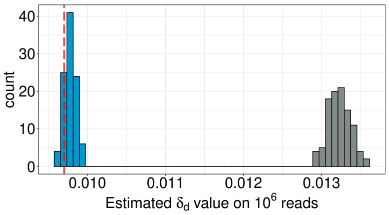

**(d)** $10^6$ reads, ngsBriggs biotin (RMSE = 0.000107) and mapDamage 2.0 (RMSE = 0.003530)

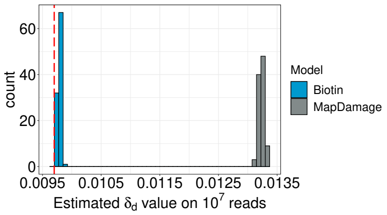

**(e)** $10^7$ reads, ngsBriggs biotin (RMSE = 8.8892e-05) and mapDamage 2.0 (RMSE = 3.5387e-03)

**Figure 8:** Inferred $\delta_d$ values using ngsBriggs Biotin and mapDamage 2.0 model for the different number of reads.

##### Inferring $\delta_s$ parameter

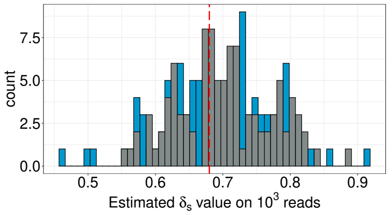

**(a)** $10^3$ reads, ngsBriggs Biotin (RMSE = 0.081506) and mapDamage 2.0 (RMSE = 0.073267)

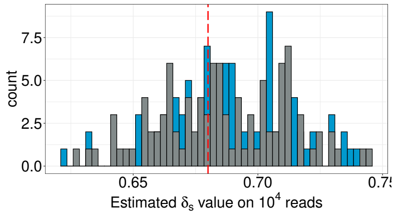

**(b)** $10^4$ reads, ngsBriggs Biotin (RMSE = 0.025015) and mapDamage 2.0 (RMSE = 0.026095)

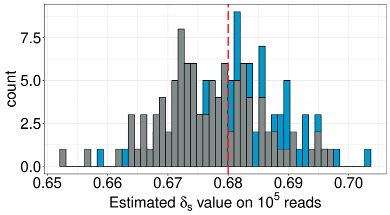

**(c)** $10^5$ reads, ngsBriggs Biotin (RMSE = 0.007957) and mapDamage 2.0 (RMSE = 0.008926)

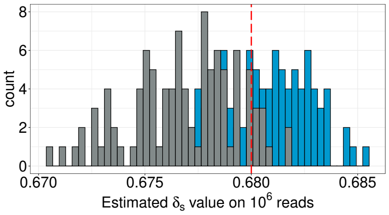

**(d)** $10^6$ reads, ngsBriggs Biotin (RMSE = 0.002331) and mapDamage 2.0 (RMSE = 0.004052)

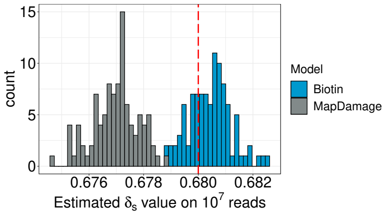

**(e)** $10^7$ reads, ngsBriggs Biotin (RMSE = 0.000898) and mapDamage 2.0 (RMSE = 0.003142)

**Figure 9:** Inferred $\delta_s$ values using ngsBriggs Biotin and mapDamage 2.0 model for the different number of reads.

##### Inferring $\nu$ parameter

The distributions for the estimated $\nu$ parameter, which is exclusively estimated by ngsBriggs, exhibit higher NRMSE values (**Figure** 10a) as opposed to the NRMSE values presented in **Figure** 2 in the main article. Albeit we observe a decrease in RMSE value as we increase the number of reads, suggesting a higher accuracy, the distributions also depict a greater disparity between the inferred values and the true value, with $10^7$ reads we observe a complete lack of overlap (**Figure** 10f). This has previously only been observed for the mapDamage 2.0 tool. However we believe based on the low RMSE value for the $10^7$ read distribution, that the lack of overlap can be ascribed to mathematical limitations in form of decimal precision when conducting the different equations as described in previous sections.

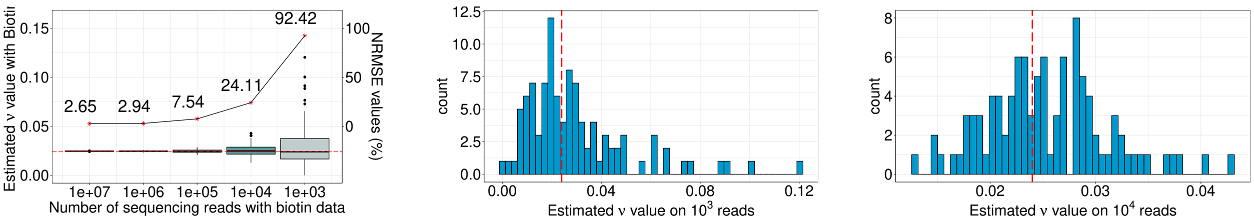

**(a)** ngsBriggs Biotin estimation of $\nu$ (0.024)
**(b)** $10^3$ reads, ngsBriggs Biotin (RMSE = 0.022180)
**(c)** $10^4$ reads, ngsBriggs Biotin (RMSE = 0.005787)

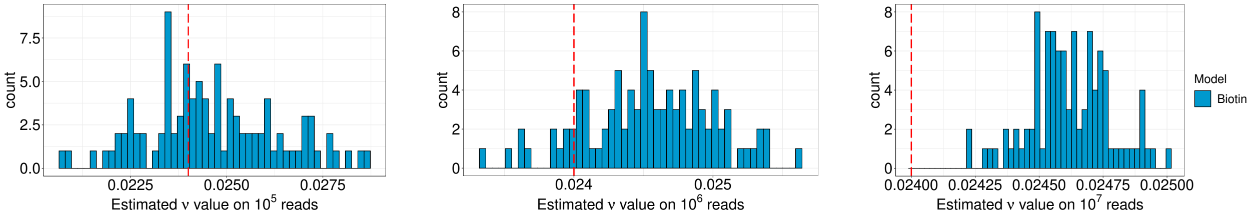

**(d)** $10^5$ reads, ngsBriggs Biotin (RMSE = 0.001811)
**(e)** $10^6$ reads, ngsBriggs Biotin (RMSE = 0.000706)
**(f)** $10^7$ reads, ngsBriggs Biotin (RMSE = 0.000637)

**Figure 10:** Inferred $\nu$ value using the ngsBriggs Biotin model for the different number of reads.

##### 6.1.2 Non-Biotin Model

To make an equivalent benchmarking of the non-biotin model, we performed an identical analysis as presented in the previous section.

For the inferred parameter the same overall trend, as observed for the biotin model, becomes clear. The consistent pattern of reduced variation, increased model disparity between mapDamage 2.0 and ngsBriggs non-biotin and decreasing RMSE values when increasing the number of reads (**Figure** 11 to 13). However, the ngsBriggs non-biotin parameter estimates appear to be situated more closely to the true value, albeit with low RMSE values comparable to those of the biotin model, suggesting both ngsBriggs are equally accurate.

#### Inferring $\lambda$ parameter

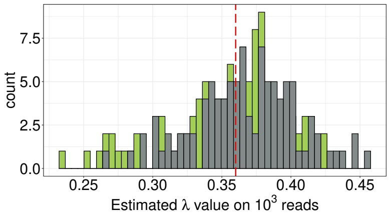

**(a)** $10^3$ reads, ngsBriggs Non-Biotin (RMSE = 0.042362) and mapDamage 2.0 (RMSE = 0.038119)

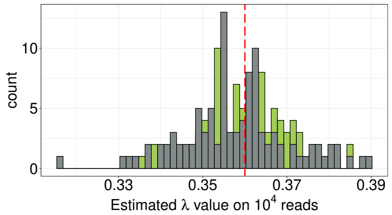

**(b)** $10^4$ reads, ngsBriggs Non-Biotin (RMSE = 0.009788) and mapDamage 2.0 (RMSE = 0.012993)

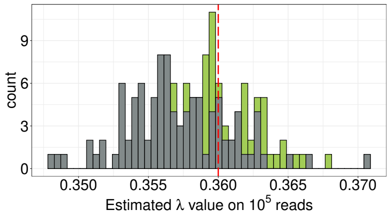

**(c)** $10^5$ reads, ngsBriggs Non-Biotin (RMSE = 0.003065) and mapDamage 2.0 (RMSE = 0.004819)

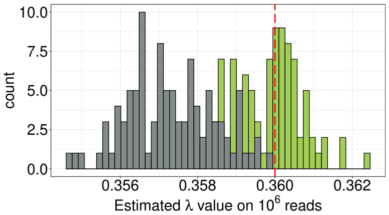

**(d)** $10^6$ reads, ngsBriggs Non-Biotin (RMSE = 0.001001) and mapDamage 2.0 (RMSE = 0.002923)

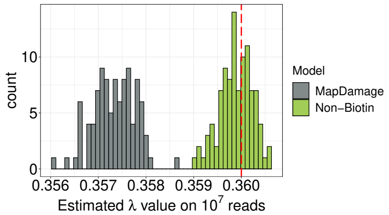

**(e)** $10^7$ reads, ngsBriggs Non-Biotin (RMSE = 0.000354) and mapDamage 2.0 (RMSE = 0.002694)

**Figure 11:** Inferred $\lambda$ values using ngsBriggs non-Biotin and mapDamage 2.0 model for the different number of reads.

#### Inferring $\delta_d$ parameter

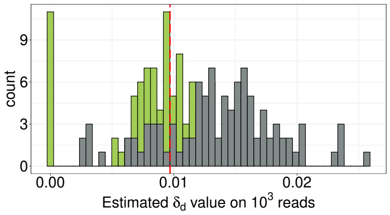

**(a)** $10^3$ reads, ngsBriggs Non-Biotin (RMSE = 0.004062) and mapDamage 2.0 (RMSE = 0.006213)

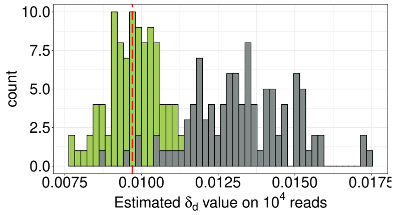

**(b)** $10^4$ reads,ngsBriggs Non-Biotin (RMSE = 0.000840) and mapDamage 2.0 (RMSE = 0.003928)

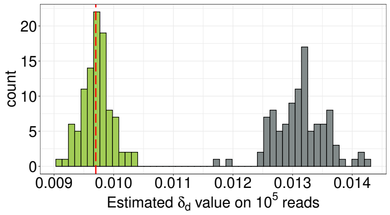

**(c)** $10^5$ reads, ngsBriggs Non-Biotin (RMSE = 0.000237) and mapDamage 2.0 (RMSE = 0.003456)

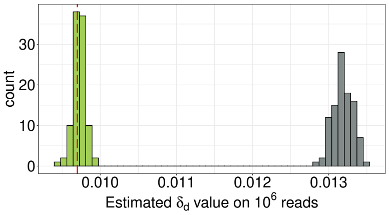

**(d)** $10^6$ reads, ngsBriggs Non-Biotin (RMSE = 0.000085) and mapDamage 2.0 (RMSE = 0.003487)

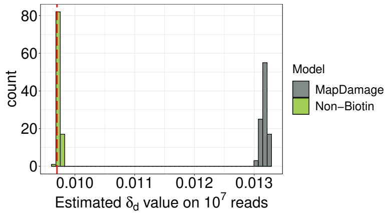

**(e)** $10^7$ reads, ngsBriggs Non-Biotin (RMSE = 0.000039) and mapDamage 2.0 (RMSE = 0.003474)

**Figure 12:** Inferred $\delta_d$ values using ngsBriggs non-Biotin and mapDamage 2.0 model for the different number of reads.

#### Inferring $\delta_s$ parameter

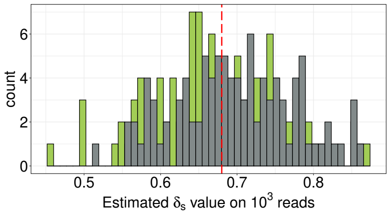

**(a)** $10^3$ reads, ngsBriggs Non-Biotin (RMSE = 0.083731) and mapDamage 2.0 (RMSE = 0.079868)

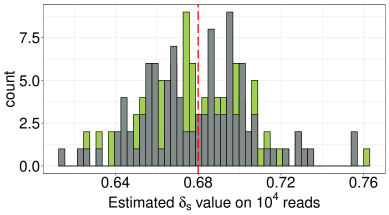

**(b)** $10^4$ reads, ngsBriggs Non-Biotin (RMSE = 0.024856) and mapDamage 2.0 (RMSE = 0.026483)

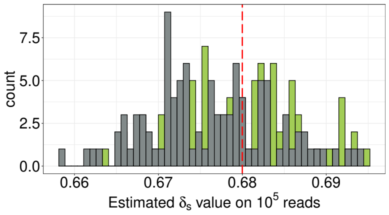

**(c)** $10^5$ reads, ngsBriggs Non-Biotin (RMSE = 0.007384) and mapDamage 2.0 (RMSE = 0.008346)

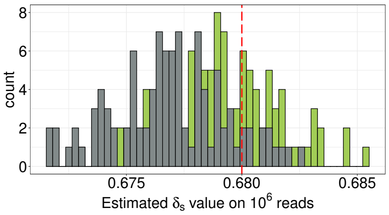

**(d)** $10^6$ reads, ngsBriggs Non-Biotin (RMSE = 0.002580) and mapDamage 2.0 (RMSE = 0.003964)

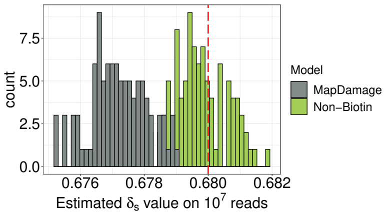

**(e)** $10^7$ reads, ngsBriggs Non-Biotin (RMSE = 0.000905) and mapDamage 2.0 (RMSE = 0.002971)

**Figure 13:** Inferred $\delta_s$ values using ngsBriggs non-Biotin and mapDamage 2.0 model for the different number of reads.

**Inferring $\nu$ parameter**

The distributions observed in **Figure** 14 reveal a similar pattern to the comparable biotin inferred $\nu$ presented in **Figure** 10, with considerably higher NRMSE value across all simulations when compared to the other parameters. However an interesting distinction emerges between the two ngsBriggs models, with the biotin model having a slight tendency to overestimate $\nu$, whereas the non-biotin model underestimates the parameter. Additionally, we observe a pronounced difference between our two deamination models. For a low amount of simulated reads, the ngsBriggs non-biotin model depicts a greater deviation from the true value (**Figure** 14b) when compared to the biotin counterpart (**Figure** 10b), while the opposite is observed for a greater number of reads comparing **Figure** 14f with 10f. This indicates that ngsBriggs non-biotin which is a representative of contemporary sequencing data, provides a more accurate estimate of the nicks occurring within aDNA.

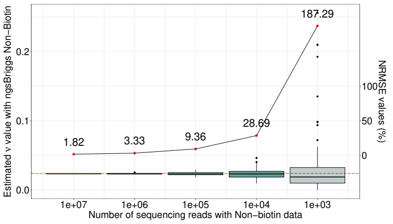

**(a)** ngsBriggs Non-biotin estimation of $\nu$ (0.024)

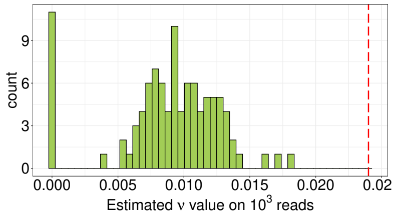

**(b)** $10^3$ reads, ngsBriggs Non-Biotin (RMSE = 0.044950)

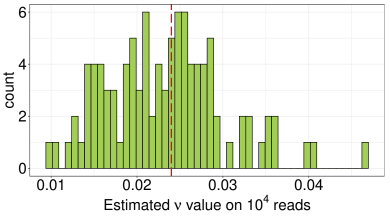

**(c)** $10^4$ reads, ngsBriggs Non-Biotin (RMSE = 0.006885)

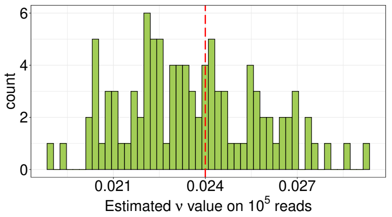

**(d)** $10^5$ reads, ngsBriggs Non-Biotin (RMSE = 0.002246)

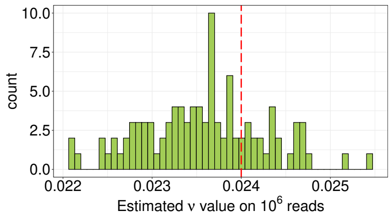

**(e)** $10^6$ reads, ngsBriggs Non-Biotin (RMSE = 0.000799)

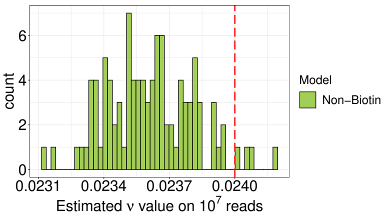

**(f)** $10^7$ reads, ngsBriggs Non-Biotin (RMSE = 0.000437)

**Figure 14:** Inferred $\nu$ values using ngsBriggs non-Biotin and mapDamage 2.0 model for the different number of reads.

#### 6.2 Deamination patterns

The empirical datasets analyzed encompass four distinct ancient populations across a varying age range from different preservation conditions. Therefore we anticipate a discernable deamination pattern. Since the applications of all three tools, ngsBriggs, mapDamage 2.0 and PMDtools are contingent upon the creation of the mismatch matrix, we need to ensure that the divergent results when inferring the parameters (**Section** 6.1), are not a consequence of dissimilar mismatch matrices. To align our expectations to the modern sequencing platforms used for the four distinct populations, we conducted a comparative analysis involving ngsBriggs (non-biotin), mapDamage 2.0, and PMDtools. We proceeded under the assumption, that when providing a reference genome, all tools would construct the mismatch matrix by aligning each position within a read to the corresponding region within the reference genome. We observe, as expected, in **Figure** 15 identical deamination frequencies created from the constructed matrix, with no visual distinction across all three tools. This supports that further potential disparity between inferred parameters for the empirical datasets without a known value is a consequence of the models utilized in each tool, further underlining the need for simulated data to ensure accuracy.

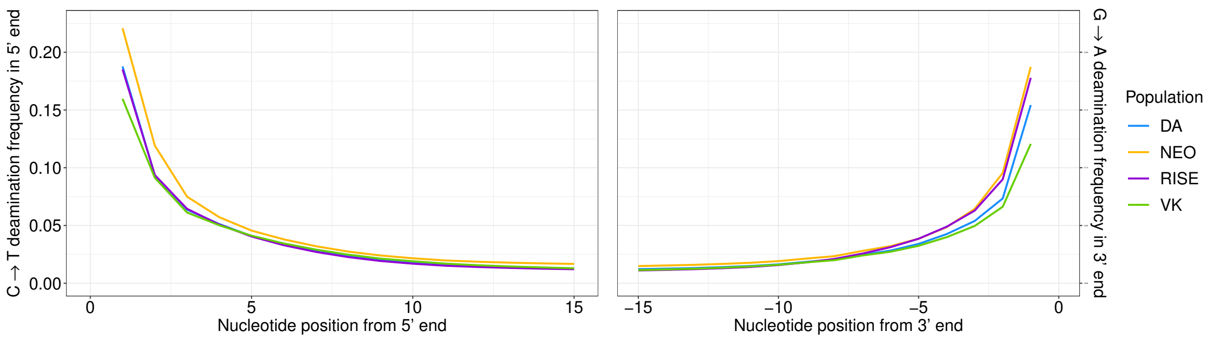

**(a)** 5' C→T frequency.

**(b)** 3' G→A frequency.

**Figure 15:** The average deamination frequencies across the individuals in each population, with identical pattern of ngsBriggs, mapDamage 2.0 and PMDtools for the first 15 base pairs from the 5' and the last 15 base pairs in the 3' end.

#### 6.3 Correlation plots

Following the successful inference of the Briggs parameters of the simulated data with known true values as outlined in **Section** 6.1, we proceeded to quantitatively compare ngsBriggs and mapDamage 2.0. By applying the two tools on empirical data without a known true parameter we aim to discern systematic differences. Subsequently, we quantified the relationship between the two tools across all the shared inferred parameters ($\lambda$, $\delta_d$, $\delta_s$).

##### 6.3.1 Inferred parameters of ngsBriggs vs mapDamage 2.0

From the inferred parameters, it becomes evident that a significant correlation between ngsBriggs and mapDamage 2.0 exists, as indicated by the Pearson-correlations and p-values below the 0.05. Noticeably we see from the black diagonal line representing identical estimates, that mapDamage 2.0 estimates slightly higher values.

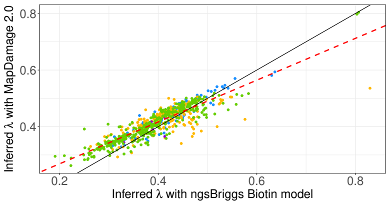

**(a)** Correlation of the $\lambda$ estimations, with Pearson correlation of 0.8881 (p-value $< 10^{-16}$)

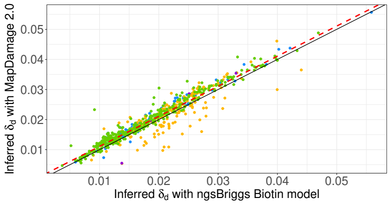

**(b)** Correlation of the $\delta_d$ estimations, with Pearson correlation of 0.9563 (p-value $< 10^{-16}$)

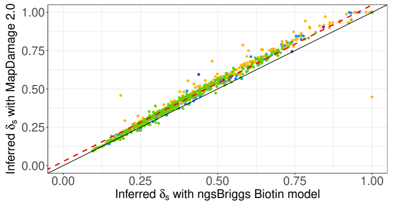

**(c)** Correlation of the $\delta_s$ estimations, with Pearson correlation of 0.9868 (p-value $< 10^{-16}$)

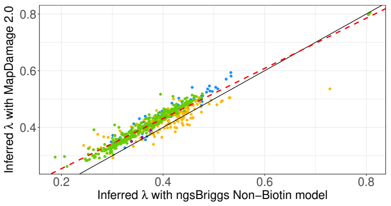

**(d)** Correlation of the $\lambda$ estimations, with Pearson correlation of 0.9388 (p-value $< 10^{-16}$)

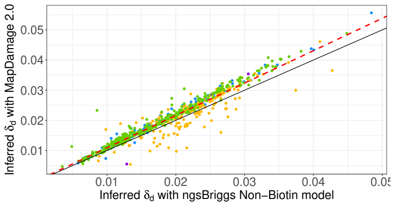

**(e)** Correlation of the $\delta_d$ estimations, with Pearson correlation of 0.9621 (p-value $< 10^{-16}$)

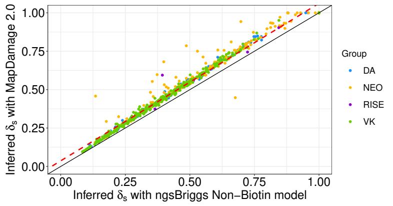

**(f)** Correlation of the $\delta_s$ estimations, with Pearson correlation of 0.9885 (p-value $< 10^{-16}$)

**Figure 16:** Correlation between the inferred parameters of the empirical dataset using ngsBriggs non-biotin model and mapDamage 2.0. The significant correlation is represented with the red-dotted line in all plots.

**Figure** 16c and 16f illustrated how, althought for a small minority of individuals, mapDmage 2.0 estimates to a greater extend parameters with extremes values of 1, while it appears ngsBriggs are more capable of estimating high $\delta_s$ vales. These extreme values suggest a discrepancy between the true PMD and the assumption that the PMD can be described using a Briggs model. However, despite these few instances, we do observe several significant linear relationships on a population level as described (**Table** 3).

| | DA | NEO | RISE | VK |
|---|---|---|---|---|
| *Biotin correlation* | | | | |
| $\lambda$ | 0.9252 (<2.2e-16) | 0.6769 (<2.2e-16) | 0.6903 (0.1290) | 0.9346 (<2.2e-16) |
| $\delta_d$ | 0.9883 (<2.2e-16) | 0.8570 (<2.2e-16) | 0.9475 (0.0041) | 0.9805 (<2.2e-16) |
| $\delta_s$ | 0.9942 (<2.2e-16) | 0.9736 (<2.2e-16) | 0.9484 (0.0039) | 0.9944 (<2.2e-16) |
| *Non-Biotin correlation* | | | | |
| $\lambda$ | 0.9548 (<2.2e-16) | 0.8357 (<2.2e-16) | 0.9341 (0.0064) | 0.9752 (<2.2e-16) |
| $\delta_d$ | 0.9929 (<2.2e-16) | 0.8570 (<2.2e-16) | 0.9564 (0.0028) | 0.9878 (<2.2e-16) |
| $\delta_s$ | 0.9942 (<2.2e-16) | 0.9736 (<2.2e-16) | 0.9304 (0.0071) | 0.9971 (<2.2e-16) |

**Table 3:** Pearson correlation values and corresponding p-values for population-specific comparison of ngsBriggs biotin with mapDamage 2.0 and ngsBriggs non-biotin with mapDamage 2.0.

##### 6.3.2 Differences in estimated parameters between models

Without a known set of PMD parameters, we're unable to use RMSE as the chosen metric while assessing the quality of our Briggs parameter estimations. Therefore we use the variation of absolute difference (quantified by its standard derivation) between estimations of corresponding parameters from mapDamage 2.0 and ngsBriggs as a metric to assess potential improvement (**Figure** 17).

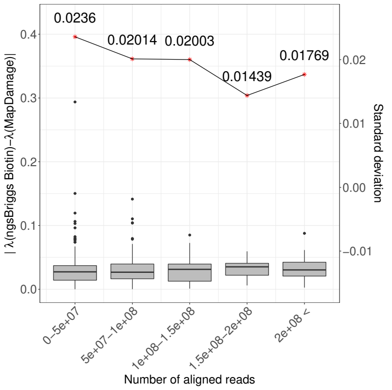

**(a)** Difference in inffered $\lambda$ for ngsBriggs Biotin and mapDamage 2.0

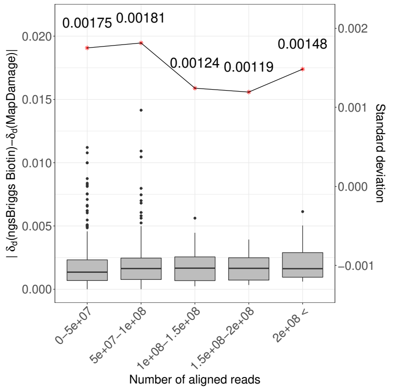

**(b)** Difference in inferred $\delta_d$ for ngsBriggs Biotin and mapDamage 2.0

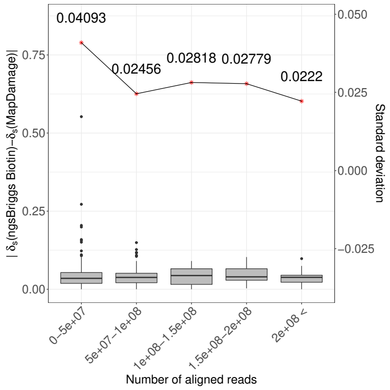

**(c)** Difference in inferred $\delta_s$ for ngsBriggs Biotin and mapDamage 2.0

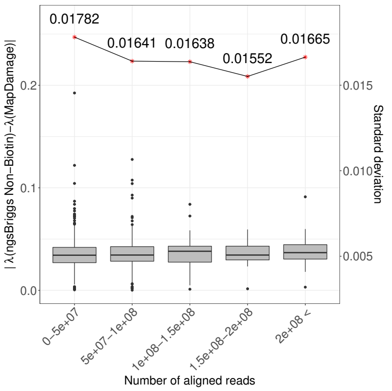

**(d)** Difference in inferred $\lambda$ for ngsBriggs Non-Biotin and mapDamage 2.0

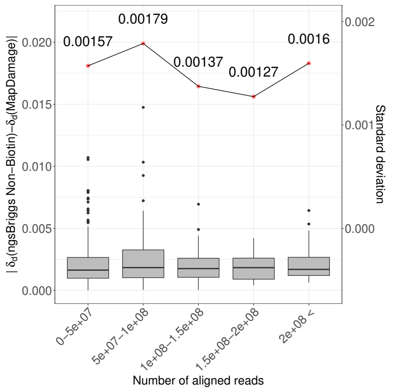

**(e)** Difference in inferred $\delta_d$ for ngsBriggs Non-Biotin and mapDamage 2.0

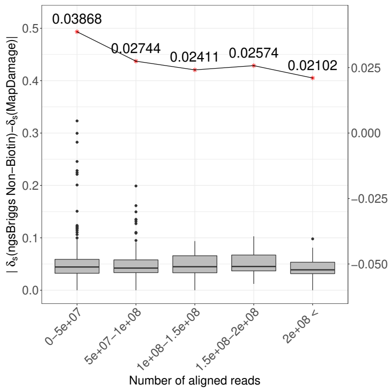

**(f)** Difference in inferred $\delta_s$ for ngsBriggs Non-Biotin and mapDamage 2.0

**Figure 17:** The difference in the three estimated parameters obtained from mapDamage 2.0 and our Briggs biotin or Briggs non-biotin model and the correlation given the number of sequencing reads.

As the number of reads increases, we observe in **Figure** 17 that the absolute differences between parameter estimates from ngsBriggs and mapDamage 2.0 consistently have stable (non-zero) means, while exhibiting relatively smaller standard deviation. This trend supports the observations depicted in **Figure** 2 in the main article, with the inferred parameters with ngsBriggs remains stable, whether we use the resulting estimates from mapDamage 2.0 as a reference, in contrast to the simulated data with known PMD values.

##### 6.3.3 Nick rate per nucleotide

The occurrence of nicks is hypothesized to serve as a potential precursor to aDNA fragmentation (Orlando *et al.*, 2021), with the inferred parameters obtained from the empirical data corroborating this belief. Under both deamination models, once ngsBriggs infers the nick rates, we can identify a negative correlation with the median read lengths (**Figure** 18). This trend could indicate, those samples with more frequent nicks, will likewise contain shorter DNA fragments. These findings lend support to the hypothesis, as nicks could potentially play a vital role in the fragmentation process, making it a likely causal factor of aDNA fragmentation.

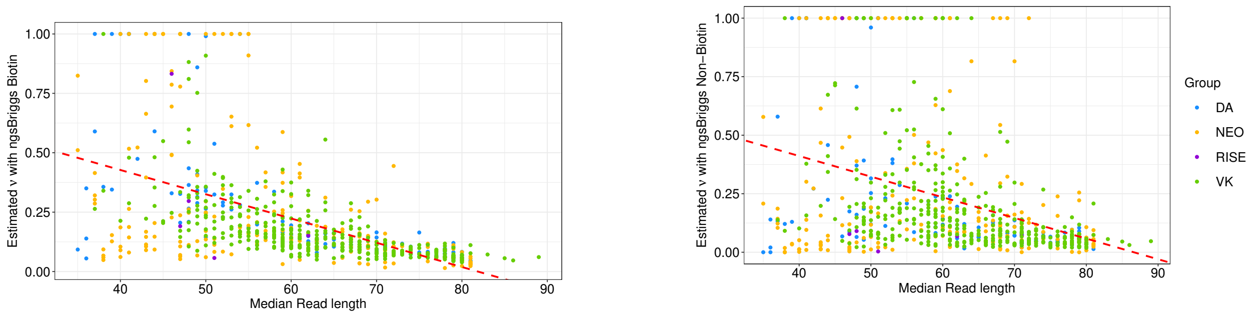

**(a)** Correlation for inferred $\nu$ value with ngsBriggs Biotin, with Pearson correlation of -0.5536 (p-value $< 10^{-16}$).

**(b)** Correlation for inferred $\nu$ value with ngsBriggs non-Biotin, with Pearson correlation of -0.3619 (p-value $< 10^{-16}$).

**Figure 18:** Correlation between the occurrence of nick within a median fragment length across all reads within an individual, identified using both ngsBriggs model, and the relationship between the two.

From **Figure** 18 we can likewise observe some extreme $\nu$ values of 1, which we similar to the extreme values previously observed (**Figure** 16), attribute to a small minority of the samples with potentially PMD patterns which are slightly irregular when assuming the usual Briggs model. Additionally, these extreme values also support the previous result for the simulated data (**Section** 6.1.1 and 6.1.2) where we do observe that the $\nu$ parameter remains the most difficult parameter to infer when compared to the three additional parameters.

#### 6.4 Relationship between inferred parameters and sample age

If the preservation conditions and treatment protocols stay similar, we hypothesized that the severeness of the DNA deamination increases monotonically with the sample age. A test has been conducted by grouping the DA, NEO, RISE and VK populations into four sample age ranges, i.e., 0 to 1250 years BCE, 1250-3750 BCE, 3750-7500 years BCE and 7500 years and older. This random grouping guarantees that within each range, there are samples from at least two populations, thus mitigates a potential location specific bias. And the results under different models are presented in Figure 19 and 20. Interestingly, we have observed that both $\delta_s$ (**Figure** 19b and 20b, both $p$ values of Jonckheere-Terpstra trend test are $< 2.2 \times 10^{-16}$), (**Figure** 19d and 20d, both $p$ values of Jonckheere-Terpstra trend test are $< 2.2 \times 10^{-16}$) $\delta_d$, and $\lambda$ (**Figure** 19a and 20a, $p$ values of Jonckheere-Terpstra trend test are $< 2.2 \times 10^{-16}$ and $= 3.346 \times 10^{-12}$, respectively) significantly increase as the sample age grows older, while we cannot draw the similar significant conclusion on $\nu$ (**Figure** 19c and 20c, $p$ values of Jonckheere-Terpstra trend test are $= 0.117$ and $= 0.0102$, respectively). The lift of the value $\delta_s$ and $\delta_d$ with the increase of sample age seems to support the previous hypothesis and suggests more ancient samples will accumulate C$\rightarrow$T per base, and the increase of $\lambda$ with sample age may be related to the phenomenon that ancient DNA sequences become shorter and shorter, however, we could not exclude the impacts introduced by location-specific preservation/treatment. t test for $\delta_s - \delta_d$ across all the ancient samples (p-values are $< 2.2 \times 10^{-16}$ under both models) could also indicate in the ancient samples, significantly higher C$\rightarrow$T rates in the single-stranded overhang regions than the double-stranded regions are always observed.

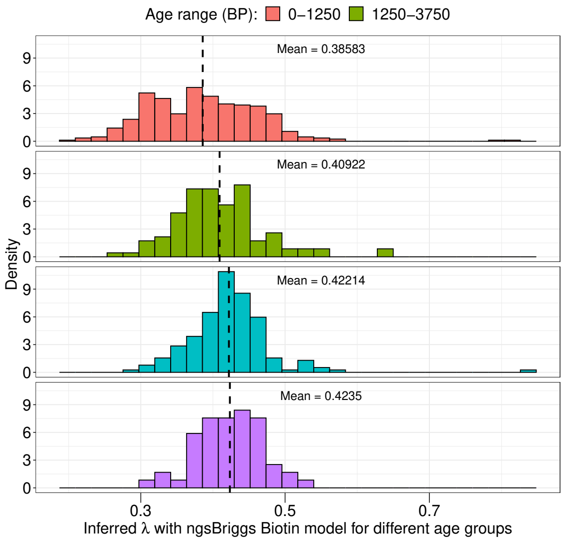

**(a)** Relationship between inferred $\lambda$ with ngsBriggs biotin model and different age ranges across populations.

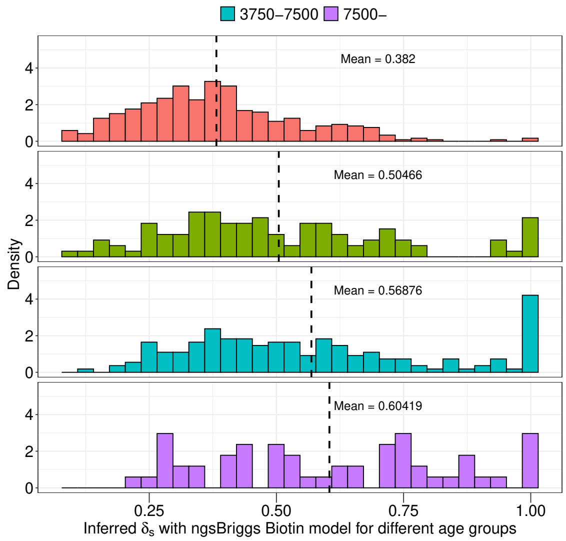

**(b)** Relationship between inferred $\delta_s$ with ngsBriggs biotin model and different age ranges across populations.

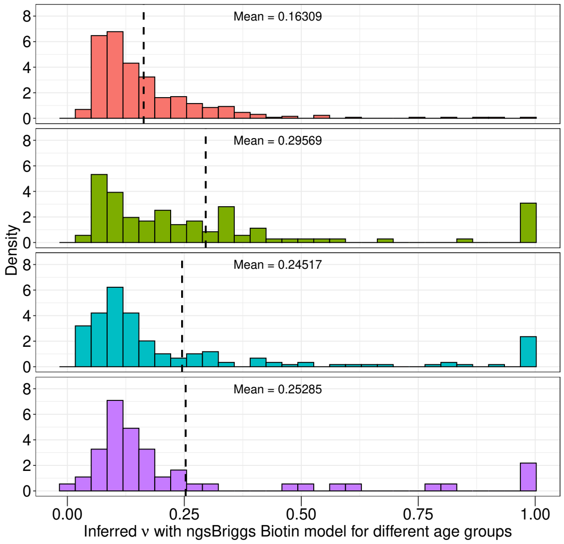

**(c)** Relationship between inferred $\nu$ with ngsBriggs biotin model and different age ranges across populations.

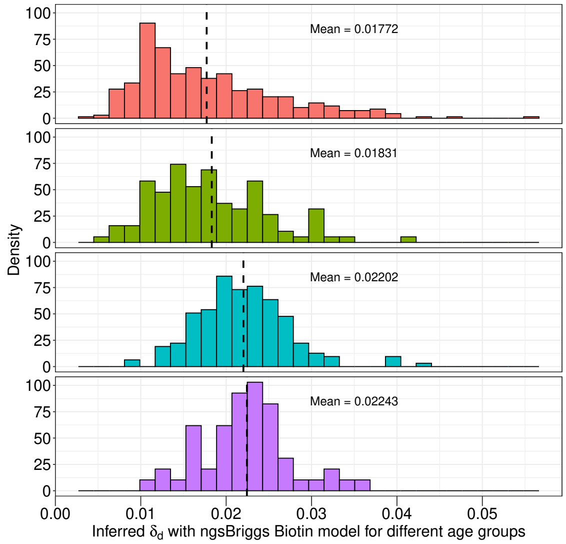

**(d)** Relationship between inferred $\delta_d$ with ngsBriggs biotin model and different age ranges across populations.

**Figure 19:** Relationship between inferred parameters with the ngsBriggs biotin model and grouped ages across the four populations (DA,NEO,RISE and VK). With the mean value of the inferred parameters being highlighted with a vertical black dotted line.

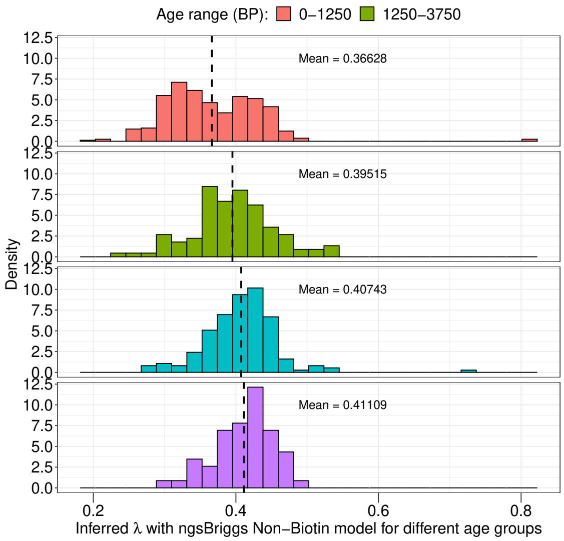

**(a)** Relationship between inferred $\lambda$ with ngsBriggs non-biotin model and different age ranges across populations.

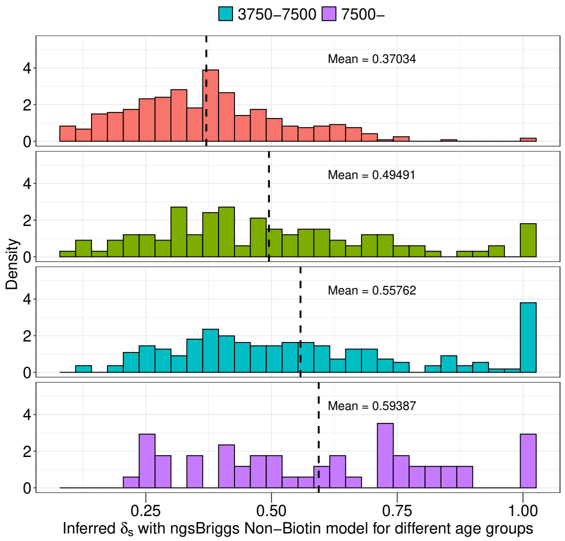

**(b)** Relationship between inferred $\delta_s$ with ngsBriggs non-biotin model and different age ranges across populations.

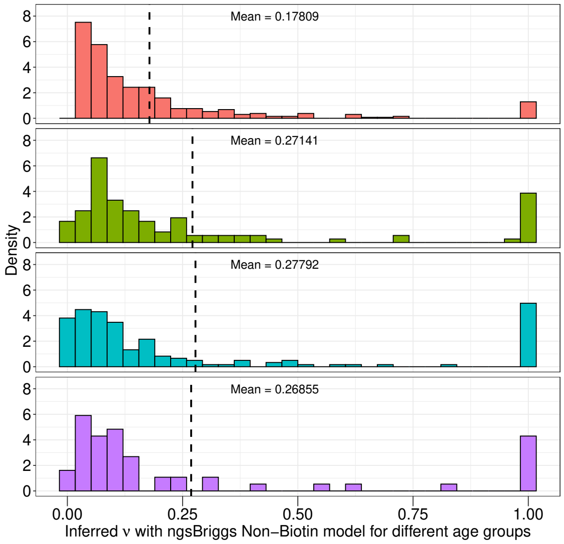

**(c)** Relationship between inferred $\nu$ with ngsBriggs non-biotin model and different age ranges across populations.

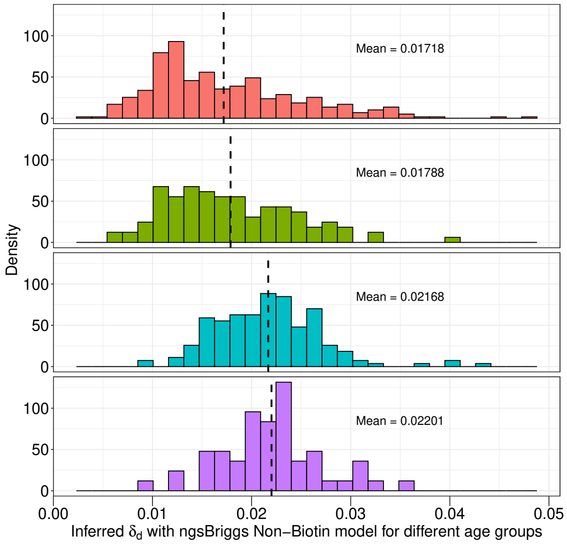

**(d)** Relationship between inferred $\delta_d$ with ngsBriggs non-biotin model and different age ranges across populations.

**Figure 20:** Relationship between inferred parameters with the ngsBriggs non-biotin model and grouped ages across the four populations (DA,NEO,RISE and VK). With the mean value of the inferred parameters being highlighted with a vertical black dotted line.

#### 6.5 Contamination

While mapDamage 2.0 is unable to account for any potential contamination, ngsBriggs offers adaptability, to incorporate a known overall contamination rate before inferring deamination parameters (**Section** 5.5). This prior knowledge can be derived from external tools such as Schmutzi (Renaud *et al.*, 2015) or ContaMix (Fu *et al.*, 2013). To examine the potential impact of contamination on our methodological procedure, we performed a comparative analysis on several contamination scenarios (**Section** 5.2) inferring the parameters with and without the prior contamination rates ($\epsilon$ value) encompassing both the biotin and non-biotin models in relation to mapDamage 2.0.

##### 6.5.1 Contamination rate's influence on parameter inference of 0.024, 0.36, 0.68 and 0.0097

In the initial comparison, we used the native parameters as presented in the original Briggs article (Briggs *et al.*, 2007) as previously described, $\nu$, $\lambda$, $\delta_d$ and $\delta_d$ having values of 0.024, 0.36, 0.68 and 0.0097.

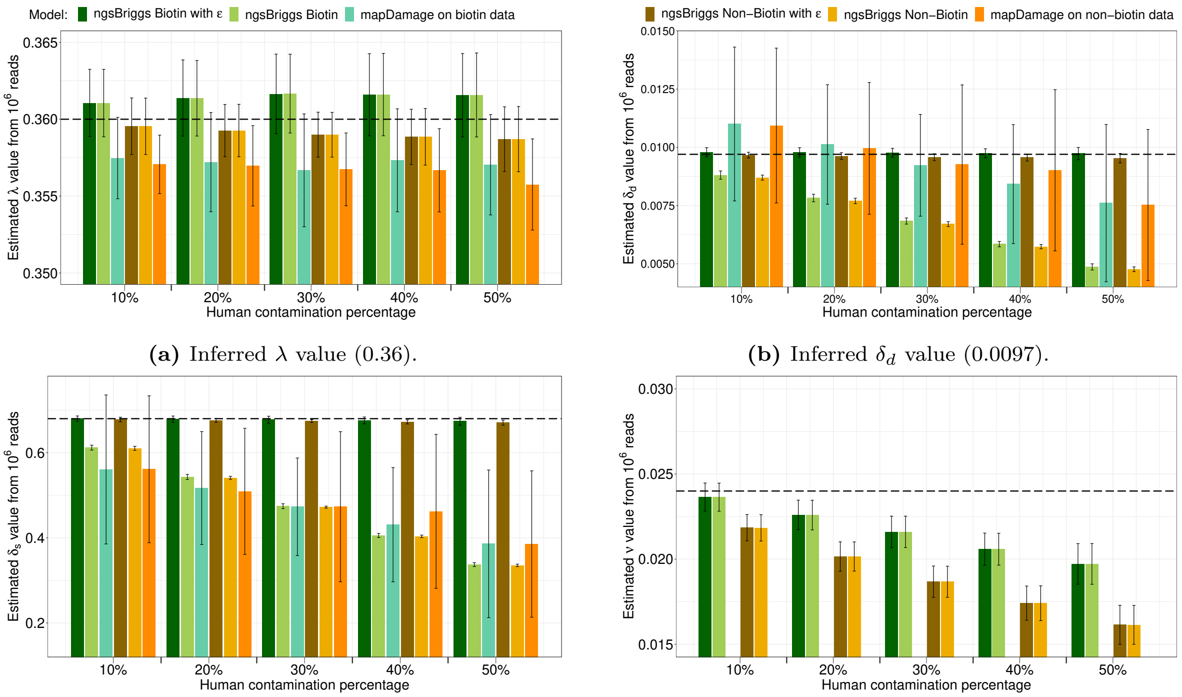

**(a)** Inferred $\lambda$ value (0.36).

**(b)** Inferred $\delta_d$ value (0.0097).

**(c)** Inferred $\delta_s$ value (0.68).

**(d)** Inferred $\nu$ value (0.024).

**Figure 21:** Inferred values of the native briggs deamination parameters for both models across the different contamination rates.

Observing the five different contamination levels, it becomes evident that the $\lambda$ parameter remains the least affected. Both tools estimate near-identical values when compared to themselves across the biotin or non-biotin data. Additionally, we observe no discernible change in either of the ngsBriggs deamination models when supplying an $\epsilon$ ratio. In contrast, the estimated $\delta_d$ and $\delta_s$ parameters exhibit susceptibility to contamination, with a substantial discrepancy between the various levels. This trend is only observable when not factoring in the contamination, i.e. mapDamage 2.0 or ngsBriggs without $\epsilon$. While providing $\epsilon$, ngsBriggs across both models consistently provides accurate estimates for both $\delta_d$ and $\delta_s$. While the $\nu$ estimates seem robust with- or without the $\epsilon$ the inferred parameters decrease with an increasing deamination level.

To verify, that the potential irregular PMD signal causing the extreme values observed during parameter inference from our empirical data are not merely artifacts from contaminants, we performed further testing on simulated data to ensure the reliability of both tools under extreme deamination conditions (**Section** 5.2). The extreme values deliberately influence the single-stranded overhang length, which remains the most susceptible segment of the fragment to undergo deamination.

##### 6.5.2 Contamination rate's influence on parameter inference of 0.024, 0.7, 0.68 and 0.0097

Using a higher $\lambda$ value, such as 0.7, the length of the single-strand overhang region decrease. Consequently, we expect a larger uncertainty- and bias in the estimation of the parameters related to the single-stranded regions, i.e. $\delta_s$.

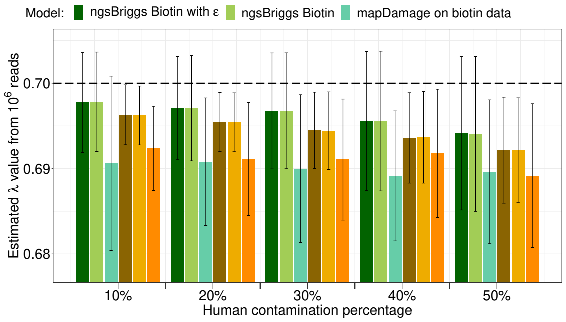

**(a)** Inferred $\lambda$ value (0.7).

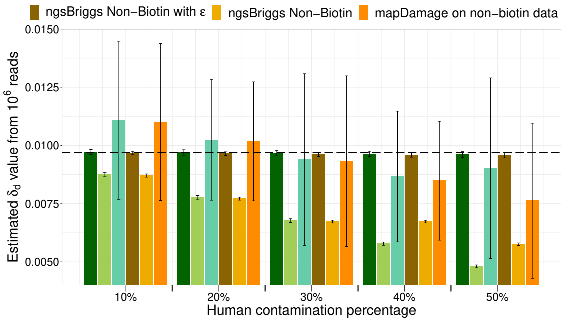

**(b)** Inferred $\delta_d$ value (0.0097).

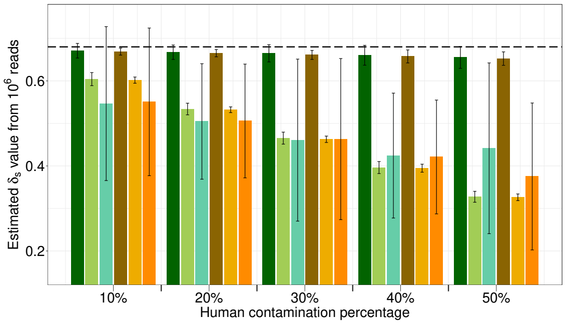

**(c)** Inferred $\delta_s$ value (0.68).

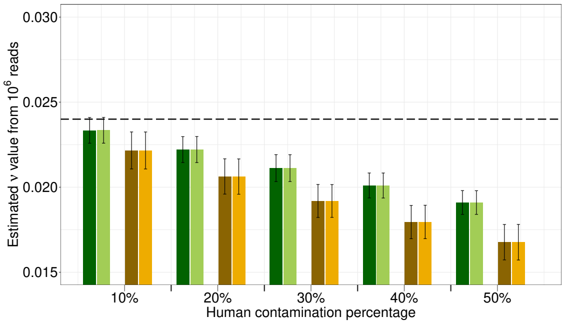

**(d)** Inferred $\nu$ value (0.024)

**Figure 22:** Inferred values of with a higher $\lambda$ parameter for both models across the different contamination rates.

With the reduced overhang length, we observe the same trend across the different scenarios as depicted in **Figure** 21. Although the inferred $\lambda$ parameter is not entirely perfect, it remains consistent for each tool across the contaminations. While slightly less accurate, when compared to **Figure** 21, it is only feasible to estimate a $\delta_s$ and $\delta_d$ close to the true PMD value by using ngsBriggs while factoring in the contamination levels.

##### 6.5.3 Contamination rate's influence on parameter inference of 0.024, 0.1, 0.68 and 0.0097

The smaller value of $\lambda$ (0.1) will increase the single-stranded overhang, and conversely decrease the length of the double-stranded region. While ngsBriggs relies on a multinomial regression approach using the information from the first- and last few cycle positions, mapDamage 2.0 MCMC approach uses all cycle positions within a read. While the distinction appears irrelevant, with a decreased double-stranded region we would expect the contamination levels to have a greater influence on ngsBriggs estimation of the $\lambda$, $\delta_d$ and $\nu$ parameter, whereas mapDamage 2.0 should be less affected as all information within the fragment is utilized.

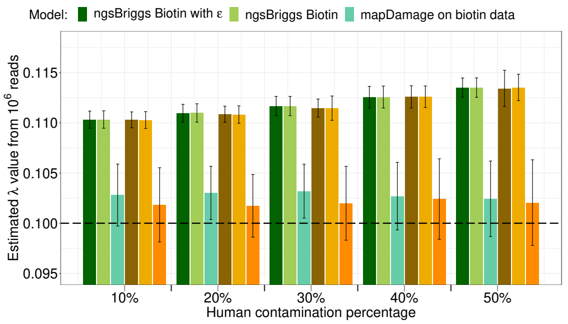

**(a)** Inferred $\lambda$ value (0.1).

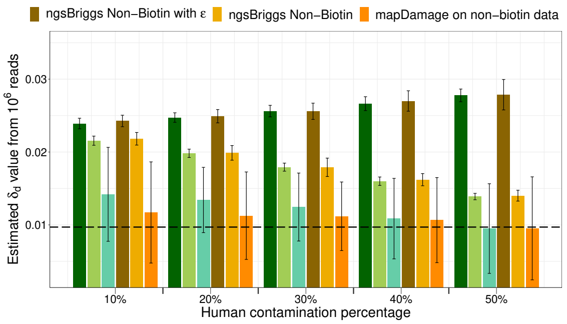

**(b)** Inferred $\delta_d$ value (0.0097).

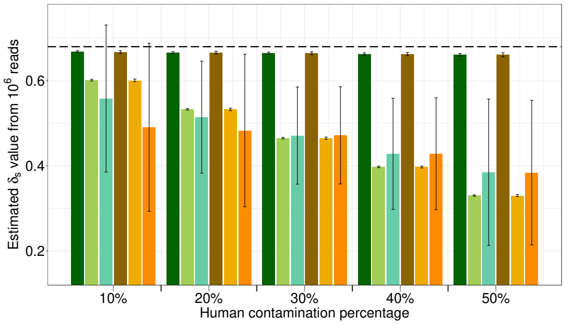

**(c)** Inferred $\delta_s$ value (0.68).

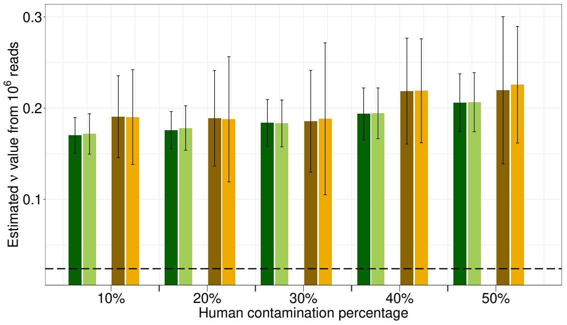

**(d)** Inferred $\nu$ value (0.024).

**Figure 23:** Inferred values of with a lower $\lambda$ parameter for both models across the different contamination rates.

The trends observed in **Figure** 23 fit our initial expectation, with the most accurate estimations, although still imperfect, of $\lambda$ and $\delta_d$ are obtainable from mapDamage 2.0, with no improvement present when factoring in the contamination for the ngsBriggs models. Similar to the previous $\delta_s$ estimates (**Figure** 21c and 22c) it becomes evident that $\delta_s$ remains the only parameter that can across all scenarios be accurately obtained when providing $\epsilon$. The $\nu$ frequency would be the most influenced by the reduced double-strand region, as this remains the sole region of the fragment it can be observed, which caused the inaccurate results.

##### 6.5.4 The overall contamination rate influence on simulated data

The visualized results described in previous sections, all support the knowledge of the well-known issue of contamination influencing analyses, in this instance the deamination parameter estimation, especially $\delta_s$ and $\delta_d$. However, we also show, given the differences within our ngsBriggs models, that this issue can be rectified by providing the prior contamination knowledge ($\epsilon$). As shown in **Figure** 21-23, since mapDamage 2.0 does not consider modern contamination, it always gives inaccurate estimations on $\lambda$, $\delta_d$ and $\delta_s$. And for ngsBriggs, it includes the effects of modern contamination and can adjust its parameter estimation according to the provided overall contamination rate $\epsilon$. More specifically, there is no difference in ngsBriggs $\lambda$ estimation (See **Figure** 21a, 22a, and 23a) with or without $\epsilon$ provided (By default if $\epsilon$ is not provided, $\epsilon$ will be set as 0 and no modern contamination will be considered) under a chosen model (the biotin model or the non-biotin model). However, if the right contamination rate $\epsilon$ is provided, the estimation of both deamination rates, i.e., $\delta_d$ and $\delta_s$ will be largely improved (See **Figure** 21b, 21c, 22b and 22c).

Though considering the effects of contamination, ngsBriggs is not perfect.

1. It assumes the fragment length distributions of ancient and modern strands are identical when calculating the likelihoods, which is not true. And this explains the ngsBriggs estimations also get worse as the overall contamination rate becomes larger (See **Figure** 21a, 21d, 22a and 22d). The length

distribution difference will be considered in the coming **Subsection** 6.5.5.

2. We note that the estimations of ngsBriggs become less accurate compared with those of mapDamage 2.0 in the case where $\lambda = 0.1$ (**Figure** 23) compared with the cases where $\lambda = 0.36$ or $\lambda = 0.7$ (**Figure** 21 and 22). The reason is as follows: When $\lambda$ is smaller, which indicates a longer overhang (single-stranded region) on average, there will be fewer double-stranded regions of aDNA that ngsBriggs could make use of, as for the sake of efficiency, it only focuses on a limited number of cyclic positions near both ends of each fragment, and the parameter estimations of the ngsBriggs related to the deamination features of double-stranded aDNA, e.g., $\lambda$, $\delta_d$ and $\nu$, will be influenced. But the accuracy of mapDamage 2.0 will not be affected by $\lambda$ values, since it always goes through all positions along each read.

##### 6.5.5 Assignment of ancient probability

With the potential impact of contamination, its become a necessity to identify which sequence reads originate from potential modern-day contaminants. Once identified, by excluding them the sample is effectively decontaminated and subsequent analysis won't be influenced. As previously described the ancient- and modern components of the simulated data have a slight overlap of the simulated fragment lengths, making it more challenging to distinguish between the endo- and exogenous content. In the other scenario with completely disjointed distributions (e.g. sequencing-by-synthesis or long-read sequencing) the decontamination could simply proceed by imposing a length threshold for read filtering.

With the default deamination parameters for the PMD signal of the ancient component, we quantified both the true- and false positive rate (TPR and FPR) when classifying the reads as ancient or modern, as presented in **Figure** 3. While ngsBriggs takes the fragment length distributions into consideration, we observe in the ROC curves (**Figure** 24), despite extreme PMD parameters (**Section** 5.3, an identical pattern as main manuscript **Figure** 3.

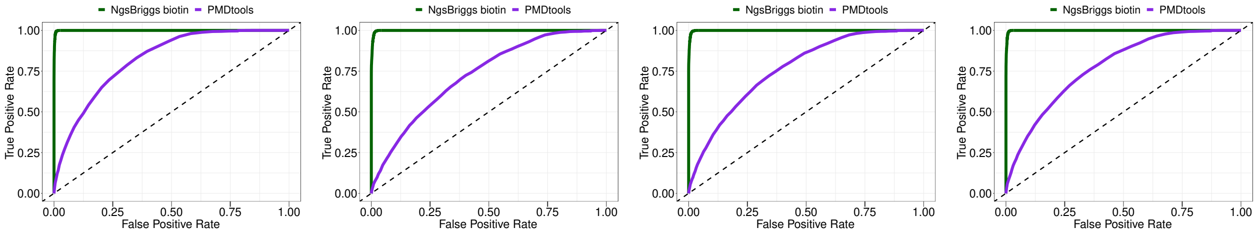

**(a)** Biotin data, $\lambda$ value of 0.1 **(b)** Biotin data, $\lambda$ value of 0.7 **(c)** Biotin data, $\delta_s$ value of 0.3 **(d)** Biotin data, $\delta_s$ value of 0.9

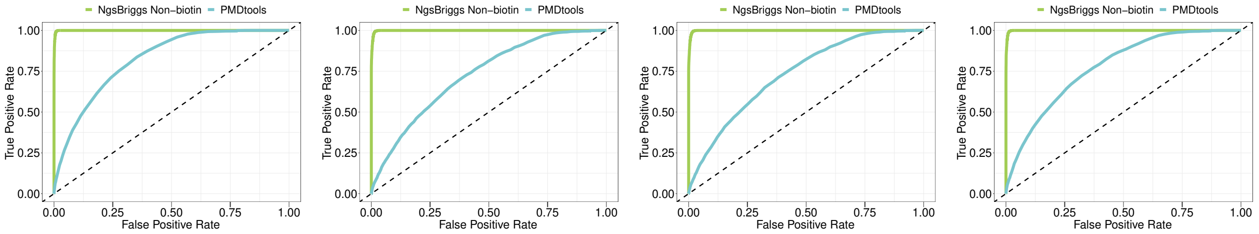

**(e)** Non-Biotin data, $\lambda$ value of 0.1 **(f)** Non-Biotin data, $\lambda$ value of 0.7 **(g)** Non-Biotin data, $\delta_s$ value of 0.3 **(h)** Non-Biotin data, $\delta_s$ value of 0.9

**Figure 24:** Measured performance test of PMDtools and both ngsBriggs models across several parameters, increasing or decreasing the deamination pattern. For the biotin data, the dark green curve corresponds to the ngsBriggs biotin and purple equalts PMDtools. The light green in the non-biotin data corresponds to the ngsBriggs non-biotin model and the light blue the PMDtools.

Across all scenarios ngsBriggs depicts a high TPR value for the lower classification thresholds, exhibiting better performance than that of PMDtools. Suggesting ngsBriggs, for both the biotin- and non-biotin model is an almost perfect classifier, as such being the optimal models to discriminate and decontaminate samples.

#### 6.6 Wall clock time - nucleotide matrix and deamination frequency

In the initial time measurement, we observe a lower mean wall-clock time when running ngsBriggs, as presented in **Table** 1 in the main article. One distinction between the ngsBriggs comparison with mapDamage 2.0 and PMDtools, is the broader range of measured wall-clock times. Most time measurements for ngsBriggs are situated within a range of 500 seconds, with a minority of outliers reaching 2007 seconds. Whereas mapDamage 2.0 and PMDtools are evenly distributed across 10.000- and 15.000 seconds respectively. Making the depiction of the wall-clock measurements more comparable on a log-scale as seen in **Figure** 25

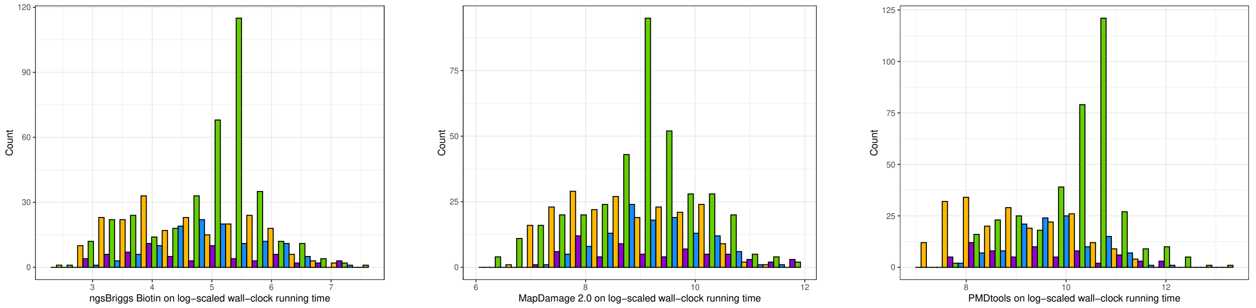

**(a)** ngsBriggs Biotin **(b)** mapDamage 2.0 **(c)** PMDtools

**Figure 25:** Wall clock time on log-scale for deamination frequency estimation using ngsBriggs, mapDamage 2.0 and PMDtools

Similarly, the disparity between the range of time measurements might make the median wall-clock a better representation, for point of comparison (**Table** 4).

| Population / Application | DA | NEO | RISE | VK |
|---|---|---|---|---|
| *Mismatch Matrix* | | | | |
| mapDamage 2.0 | 11603.20 | 6052.36 | 8072.52 | 8441.08 |
| PMDtools | 19441.30 | 7001.02 | 10315.80 | 31006.90 |
| ngsBriggs | 168.00 | 87.00 | 82.00 | 169.50 |
| *Parameter inference* | | | | |
| mapDamage 2.0 | 1117.38 | 1475.23 | 1122.19 | 782.52 |
| ngsBriggs | 2.00 | 2.00 | 2.00 | 1.00 |

**Table 4:** The first three lines represents the mean wall-clock running time of the generated nucleotide matrices used to calculate the 5' C→T and 3' G→A deamination frequencies. The next three lines represent the wall clock running time of the briggs parameter inference.

Across the two applications compared, we observe a significant increase when using ngsBriggs as opposed to either mapDamge 2.0 and PMDtools. The large difference when constructing the mismatch matrix is a consequence of each tools distinct procedure. As previously mentioned ngsBriggs solely relies on the first- and last number of cycles, which is the regions relevant for deamination, while mapDamage during the construction of the mismatch matrix uses the information to generate multiple output files, increasing the Output bottleneck. PMDtools as presented in **Section** 5.4 use the standard output once, *samtools* (Li *et al.*, 2009; CB *et al.*, 2022) have opened the entire alignment file, which with an increasing number of NGS reads generated will likewise increase the Input bottleneck.

##### 6.6.1 Nucleotide correlations

While mapDamage 2.0 employs a MCMC approach across all nucleotides when inferring the parameters, ngsBriggs as mentioned rely on a multinomial regression of the first- and last few cycle positions. Thus we measure the number of nucleotides impact on the wall-clock running time.

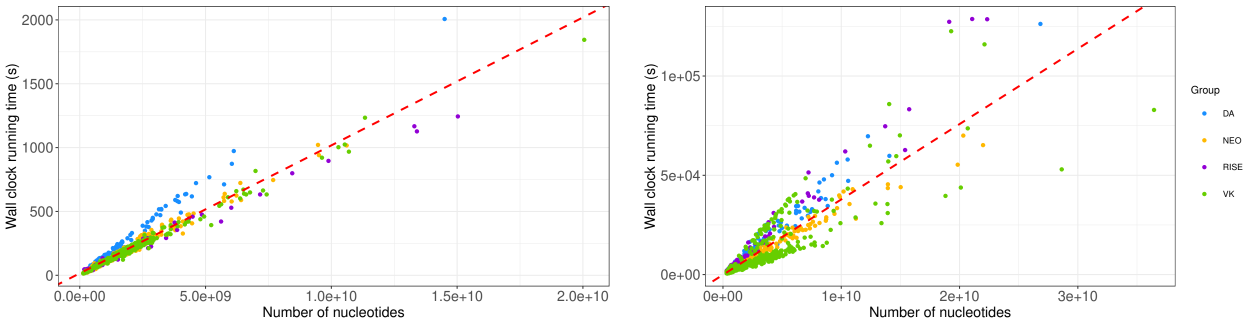

**(a)** NgsBriggs biotin model, using 15 nucleotides in both 5' and 3' end, with Pearson correlation of 0.9707 (p-value $< 10^{-16}$).

**(b)** mapDamage 2.0, using all nucleotides within a sequence read, with Pearson correlation of 0.8840 (p-value $< 10^{-16}$).

**Figure 26:** Relationship between the wall-clock running time and number of nucleotides used to count the PMD pattern.

For both tools, we see a linear relationship, indicating, as expected with a greater number of reads and nucleotides the wall-clock running time increases. We do observe that our tool (**Figure** 26a) has a smaller slope, which further supports that our tool is more time-efficient.

### 7 Future persepectives

The presented tool, ngsBriggs is implemented as a stand-alone multi-threaded tool that can be used to determine the parameters accurately describing the PMD signal and decontaminate single genome data. Furthermore, as stated in the main article, the implementation and applications of ngsBriggs are developed within a framework designed to analyse ancient environmental samples (aeDNA) with the toolkit metaDMG (Michelsen *et al.*, 2022). Within metagenomic analysis, the stand-alone ngsLCA (Wang *et al.*, 2022) obtains taxonomic profiles for all taxonomical ranks from the NCBI taxonomy database by inferring the lowest common ancestor for multiple aligned reads. Each taxonomical profile contains a mismatch matrix and mean read length amongst other alignment statistics. While, a stand-alone tool, the applications of ngsLCA is similarly part of the metaDMG toolkit. As such the information stored in the taxonomical profiles, i.e. mismatch matrix and the internal read lengths distribution (used for the various computed metrics) can be utilized as input to ngsBriggs (illustrated with the **green** box, in **Figure** 1) to infer the parameters of the PMD signal across all taxonomical ranks, to accurately determine the ancient organism present in the increasingly used environmental samples.
